# Global impact of aberrant splicing on human gene expression levels

**DOI:** 10.1101/2023.09.13.557588

**Authors:** Benjamin Fair, Carlos Buen Abad Najar, Junxing Zhao, Stephanie Lozano, Austin Reilly, Gabriela Mossian, Jonathan P Staley, Jingxin Wang, Yang I Li

## Abstract

Alternative splicing (AS) is pervasive in human genes, yet the specific function of most AS events remains unknown. It is widely assumed that the primary function of AS is to diversify the proteome, however AS can also influence gene expression levels by producing transcripts rapidly degraded by nonsense-mediated decay (NMD). Currently, there are no precise estimates for how often the coupling of AS and NMD (AS-NMD) impacts gene expression levels because rapidly degraded NMD transcripts are challenging to capture. To better understand the impact of AS on gene expression levels, we analyzed population-scale genomic data in lymphoblastoid cell lines across eight molecular assays that capture gene regulation before, during, and after transcription and cytoplasmic decay. Sequencing nascent mRNA transcripts revealed frequent aberrant splicing of human introns, which results in remarkably high levels of mRNA transcripts subject to NMD. We estimate that ∼15% of all protein-coding transcripts are degraded by NMD, and this estimate increases to nearly half of all transcripts for lowly-expressed genes with many introns. Leveraging genetic variation across cell lines, we find that GWAS trait-associated loci explained by AS are similarly likely to associate with NMD-induced expression level differences as with differences in protein isoform usage. Additionally, we used the splice-switching drug risdiplam to perturb AS at hundreds of genes, finding that ∼3/4 of the splicing perturbations induce NMD. Thus, we conclude that AS-NMD substantially impacts the expression levels of most human genes. Our work further suggests that much of the molecular impact of AS is mediated by changes in protein expression levels rather than diversification of the proteome.

## Background

Alternative splicing (AS) has the potential to dramatically expand the number of functional peptides encoded in mRNA. Large-scale transcriptomics studies have confirmed that nearly all protein coding genes generate multiple – sometimes dozens – of distinct mRNA isoforms. This finding is often interpreted as supporting the role of AS in diversifying the proteome; however, most alternatively spliced isoforms are lowly expressed and lack cross-species conservation^1–6^. To explain these observations, multiple studies have suggested that the vast majority of mRNA isoforms are nonfunctional transcripts that result from mis-splicing rather than regulated AS^5–10^.

Mis-splicing events, such as those resulting from aberrant activation of unconserved cryptic splice sites, often introduce frameshifts in the mRNA coding sequence, resulting in premature termination codons (PTCs). Consequently, downstream exon junction complexes – which would normally be displaced by translating ribosomes – recruit nonsense-mediated decay (NMD) machinery to the mRNA for degradation. Thus, most transcripts with one or more aberrant splicing events are considered to be ‘unproductive’, as they are expected to undergo rapid NMD.

Unproductive transcripts can also result from regulated AS^11–23^. For example, splicing factors have been found to control AS of their own pre-mRNA, relying on the coupling between alternative splicing and NMD (AS-NMD) to autoregulate their expression levels^13–15,18^. However, regulated AS-NMD has only been documented in a handful of genes, suggesting that AS-NMD has a very modest impact on the expression levels of most human genes. In support of this view, early studies of AS-NMD inhibited core NMD factors such as *UPF1* or *UPF2* and observed an effect on expression levels for only ∼10–30% of genes^24–27^, suggesting that the vast majority of transcript molecules are spliced into productive transcripts^25^. However, more recent studies^28–34^ find that partial redundancy can obscure knockdowns of single NMD factors, possibly masking the full impact of AS-NMD on mRNA expression levels. Thus, the main mechanism by which AS impacts molecular and organismal phenotypes – whether by altering mRNA abundance or protein isoform diversity – remains unclear^11,16,35–38^.

In this study, we use molecular data collected before, during, and after transcription and mRNA decay to estimate the effect of AS-NMD on gene expression levels. We discover that splicing-mediated effects on gene expression levels and organism-level phenotypes are considerable and pervasive, substantially affecting nearly all human genes. We find that the majority of these AS-NMD events are consistent with aberrant splicing and likely result from unexpectedly high rates of errors in the splicing of human introns. Thus, the diversity of mRNA isoforms detected in existing studies is eclipsed by an even larger number of unannotated, unproductive mRNA isoforms that are subject to rapid degradation by NMD. We conclude that the impact of AS on gene expression levels has been greatly underestimated, and that the importance of AS-NMD in explaining human phenotypic variation may even exceed that mediated by the protein-diversifying function of AS.

## Results

### High-throughput measurements of alternative splicing before mRNA decay

To assess the impact of AS on steady-state gene expression levels, we must jointly consider multiple stages of gene regulation that reflect mRNA before and after cytoplasmic degradation. To do this, we leveraged a large collection of molecular assays in lymphoblastoid cell lines (LCLs) derived primarily from 40–86 Yoruba individuals from Ibadan, Nigeria (**Figure 1A**, **Figure S1)**. These datasets^39–42^ have been widely used to study the impact of genetic variants on molecular phenotypes, and consists of measurements tracking major steps of mRNA biogenesis including chromatin activity at enhancers (H3K4me1 and H3K27Ac ChIP-seq) and promoters (H3K27Ac and H3K4me3 ChIP-seq), newly transcribed polyA RNAs (4sU pulse-labeled for 30- or 60-minutes), and steady-state mRNA levels (RNA-seq). However, these data fail to capture spliced mRNA before potential cytoplasmic degradation, preventing us from capturing rapidly degraded mRNA transcripts.

**Figure 1:**
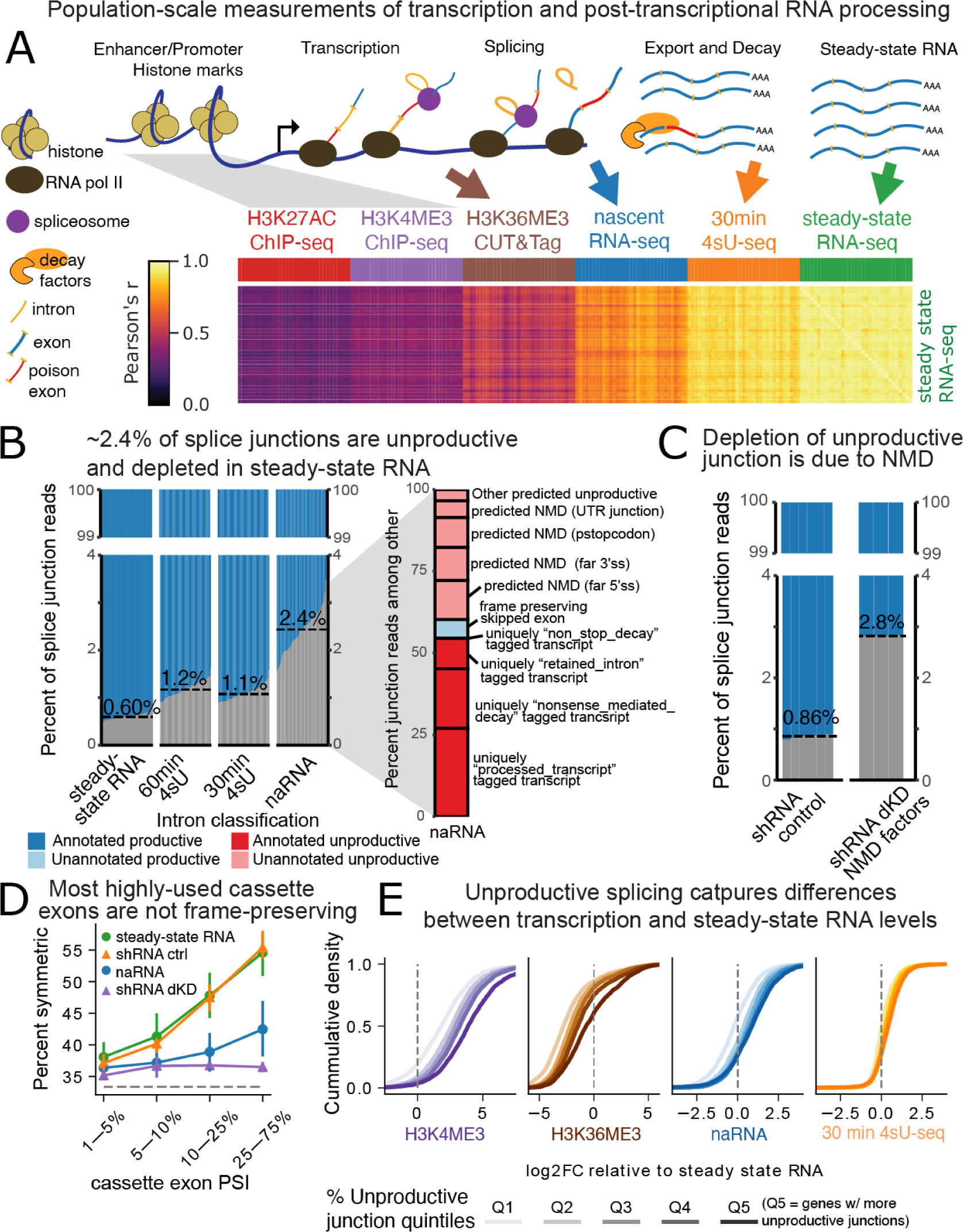
Genomic data captured before, during, and after transcription reveal an abundance of NMD isoforms. (A) A subset of the population-scale datasets we analyzed, covering stages of mRNA biogenesis, from activation of enhancers/promoters (i.e. H3K27ac and H3K4me3 ChIP-seq), to steady-state RNA (polyA-RNA-seq). Gene expression correlation matrix relative to steady-state RNA samples shown as heatmap (B) Left: Fraction of splice junction reads in each RNA-seq sample that are in Gencode-annotated productive transcript structures (blue), versus unannotated or annotated unproductive transcript structures (gray). Dashed lines indicate median for each dataset. Right: For the 2.4% of splice junctions in naRNA-seq data that are not in annotated productive transcript structures, we checked for their unique presence in annotated unproductive transcript structures (e.g., transcripts tagged by Gencode as “retained_intron”), or if unannotated, we attempted to translate sequences surrounding the splice sites (STAR Methods). Stacked bar plot indicates the fraction of naRNA-seq splice junctions in each sub-category. (C) Similar to (B), comparing steady-state RNA from shRNA scramble control (n=6) and shRNA double knockdown (dKD, n=3) of *SMG6* and *SMG7* in HeLa cells^28^. (D) Fraction of cassette exons that are symmetric (i.e. length divisible by three) as a function of their usage, estimated as percent spliced in (PSI). Error bars represent the 5-95% percentile range of values for LCL lines treated as replicates (circular markers, same dataset as (B)), and the full range of values for replicate shRNA knockdown experiments (triangular markers, same dataset as (C)). (E) Cumulative distribution of log fold differences in steady-state gene expression versus gene transcription (measured by H3K4me3 promoter activity, naRNA, H3K36me3, or 30m 4sU-labeled RNA), a proxy for degradation rate. Genes are grouped by quintiles based on percent of unproductive junction reads.

To measure AS of pre-mRNA splicing before mRNA decay, we used nascent RNA-seq (naRNA-seq), which captures chromatin-associated RNA before it can be degraded in the cytoplasm. We obtained a total of 22.4 billion naRNA-seq reads across 86 LCLs. To further increase the temporal resolution of our dataset, we also collected CUT&Tag data from 95 LCLs to profile H3K36me3, a mark associated with active transcription elongation (**Figure S2**). These data are critical for discriminating between transcriptional and splicing-mediated effects on gene expression levels.

The correlation between gene expression measurements in steady-state RNA and gene expression measurement at previous stages of RNA processing reveals a clear temporal pattern, matching our expectation that our naRNA-seq data captures mRNA at an earlier stage of maturation than 4sU-seq labeled for 30 minutes (**Figure 1A**). Together, all our molecular measurements capture the flow and loss of information across regulatory processes from chromatin to steady-state mRNA expression with unprecedented resolution.

We confirmed that naRNA-seq captures RNA associated with chromatin and nascent pre-mRNA in several ways. First, we observed that non-coding RNAs known to be depleted in steady state polyA RNA, such as long non-coding RNAs, upstream-antisense RNAs, and small nucleolar RNAs, were present in our naRNA-seq data, but not in metabolic labeled RNA (30 min 4sU-seq) or steady-state RNA-seq data, which were collected from the same LCLs in previous studies (**Figure S3A**). Second, we found that a much larger fraction of naRNA-seq reads mapped to the introns of protein coding genes, which is reflected by marked differences in intron persistence across the three datasets (**Figure S3B**). Co-transcriptional splicing is evident in the aligned naRNA-seq data, as is a 5’ bias in intronic reads, together producing a clear sawtooth pattern at long introns that has been documented^43–45^ to reflect co-transcriptional RNA processing (**Figures S3C,D**). The abundance of co-transcriptional splicing, together with the deep coverage of our naRNA dataset (**Figure S3E**), allows high resolution analysis of splicing outcomes prior to cytoplasmic degradation.

### Unproductive mRNA splicing is pervasive

We used naRNA-seq data to estimate the prevalence of ‘unproductive splicing’, i.e. splicing outcomes that are expected to induce NMD of the host transcript. In contrast, ‘productive splicing’ is expected to preserve the proper reading frame of the host transcript. Thus, exon-exon junction reads from unproductive transcripts are expected to be depleted in steady-state RNA-seq, while the junction reads identified in naRNA would reflect unbiased rates of unproductive splicing. We find 0.44% of all junction reads overlapping protein-coding genes from naRNA-seq map to splice junctions uniquely annotated in NMD-targeted transcripts (GENCODE annotations^46^), compared to 0.15% in steady-state RNA (**Figure S4A**). Moreover, we find that 2.4% of junction reads in naRNA are not attributable to annotated junctions of stable protein-coding transcripts, compared to 1.1% in 30min labeled 4sU RNA and 0.60% in steady-state RNA (Figure 1B). As only a small fraction of these 2.4% of junction reads can be attributed to annotated NMD transcripts, we sought to understand whether the remaining represent unproductive isoforms. We find that about half of these remaining junctions are annotated in transcripts that are expected to be NMD substrates, albeit not explicitly defined as so by Gencode (e.g., transcripts associated with a retained intron) (**Figure 1B**, **Figures S4B,C**, STAR Methods). Most of these splice junctions are greatly depleted in steady-state RNA (**Figure S5A**), consistent with their rapid decay. To categorize the completely unannotated splice junctions, we developed a method to predict their effect on transcript coding potential (STAR Methods). We find that these unannotated junctions are overwhelmingly expected to either frameshift or introduce a PTC in the coding sequence, thus resulting in a transcript targeted by NMD. To confirm the quality of our productive/unproductive categorizations of splice junctions, we assessed the change in abundance of splice junctions in shRNA-induced knockdowns of core NMD components, including single knockdowns of *UPF1*, *SMG6*, and *SMG7*, and a double knockdown of *SMG6* and *SMG7* ^28^. As expected, annotated and unannotated splice junctions that we classify as productive are relatively unchanged upon NMD knockdown, while unproductive junctions increase in abundance (**Figure S5**). Importantly, single knockdown of core NMD factors displayed much smaller abundances of unproductive splice junctions compared to double *SMG6* and *SMG7* knockdowns (**Figure S6**), suggesting functional redundancy between these NMD factors. Overall, we find a similar enrichment of unproductive splice junctions in the double knockdown experiments (**Figure 1C**) as in our naRNA (**Figure 1B**), suggesting NMD is the primary mechanism explaining the abundance of unannotated junctions in naRNA. Thus, we estimate that ∼2.3% of splicing events target transcripts for NMD, as measured in naRNA, compared to ∼0.55% in steady-state (**Figure S4C**). Our estimate of ∼0.55% unproductive splice junctions in steady-state RNA, which is robust across our panel of LCLs (**Figure S4C**), is consistent with a previous report that uses steady-state RNA to estimate a ∼0.7% splicing error rate for the typical human intron^6^. However, our observation in nascent RNA that ∼2.3% of splicing events are unproductive (**Figure 1B, Figure S4C**) is unprecedentedly high.

We next sought to further verify that unproductive isoforms are abundant without relying on our annotation of productive/unproductive junctions. To this end, we analyzed exons whose length is divisible by three (“symmetric exons”), and thus are frame-preserving whether the exon is skipped or included. Previous studies using steady-state RNA-seq data found that highly-included and conserved alternatively spliced exons (“cassette exons”) are biased towards being symmetric^1–4,47^, suggesting that there is selective pressure to maintain coding frame for highly-included cassette exons. Indeed, ∼55% of highly-included cassette exons are symmetric compared to only ∼35% of rarely-used cassette exons (Percent spliced-in, PSI < 1%). However, we find that in naRNA the fraction of symmetric exons is low, under 40%, even for the most highly-included cassette exons (**Figure 1D**). We find similar results in steady-state RNA after knockdown of NMD factors (**Figure 1D**). These observations indicate that the enrichment for frame-preserving exons in steady-state RNA is largely a product of NMD surveillance, instead of supporting the widely-held belief that the main function of AS is protein diversification.

Having observed such abundant unproductive splicing, we wondered to what extent AS-NMD influences gene expression levels genome-wide. Specifically, we binned protein coding genes into quintiles based on the genewise percent of junctions that are predicted to be unproductive and compared estimates of mRNA degradation across the quintiles. The percent of unproductive splicing varies widely across genes, with hundreds of genes containing >20% unproductive junctions (**Figure S7A**). Degradation was estimated as the difference in expression observed in steady-state RNA, versus various measures of transcriptional activity, including promoter activity (H3K27ac) and transcription elongation (H3K36me3) (**Figure 1E**). Indeed, quintiles of higher percent unproductive junctions correspond to higher degradation rates, though this effect largely disappears when using degradation rates calculated from polyA-selected 30min 4sU labeled RNA, consistent with previous reports^49,50^ that suggest most NMD occurs quickly after mRNA maturation. Using linear modeling, we conservatively estimate that AS-NMD explains at least 9% of post-transcriptional gene expression regulation variance across genes (Supplemental Note 1). Though, it is not clear to what extent AS-NMD is functionally regulated and optimized through selection, versus a result of ‘noisy’ splicing mistakes which nonetheless impact gene expression.

To better understand whether AS-NMD is generally functionally regulated versus noisy, we explored which classes of genes and introns are most prone to AS-NMD. We find that highly expressed (**Figure S7**) and evolutionarily constrained genes (**Figure S8A**) have among the lowest rates of unproductive splicing. Because evolutionarily constrained genes are expected to be under strong selective pressure to be tightly and accurately regulated^51^, the fact that we see the least unproductive splicing in them suggests that unproductive splicing generally represents molecular noise, rather than a form of regulation. However, we find exceptions – for example, the highly-conserved SR proteins have among the highest rates of unproductive splicing even when controlling for expression (**Figure S7B**, Supplemental Note 2), consistent with previous reports of conserved AS-NMD-based auto-regulatory loops in these genes^13–15,18,20^. Still, splicing regulators represent only a small fraction of genes with the highest rates of unproductive splicing (Supplemental Note 2). Further, we observe that introns with weak splice sites (**Figure S8B**,**C**) and long introns (**Figure 2A**) have higher unproductive splicing rates, consistent with splicing errors caused by appreciable competition between *bona fide* functional splice sites and cryptic splice sites. We also reasoned that if unproductive splicing is regulated and optimized, most unproductive splicing would be derived from a single alternative event that could be efficiently regulated by cis-regulatory elements, as is the case for SR protein autoregulatory loops. Yet, for most genes, we were unable to attribute a majority of unproductive splice events to a single splice junction (**Figure 2B**). Instead, we observed that unproductive splicing events tend to be distributed across a collection of rarely used junctions (**Figure S9**), with some genes requiring more than five unique splice junctions to explain the majority of unproductive junction reads (**Figure 2B**).

**Figure 2:**
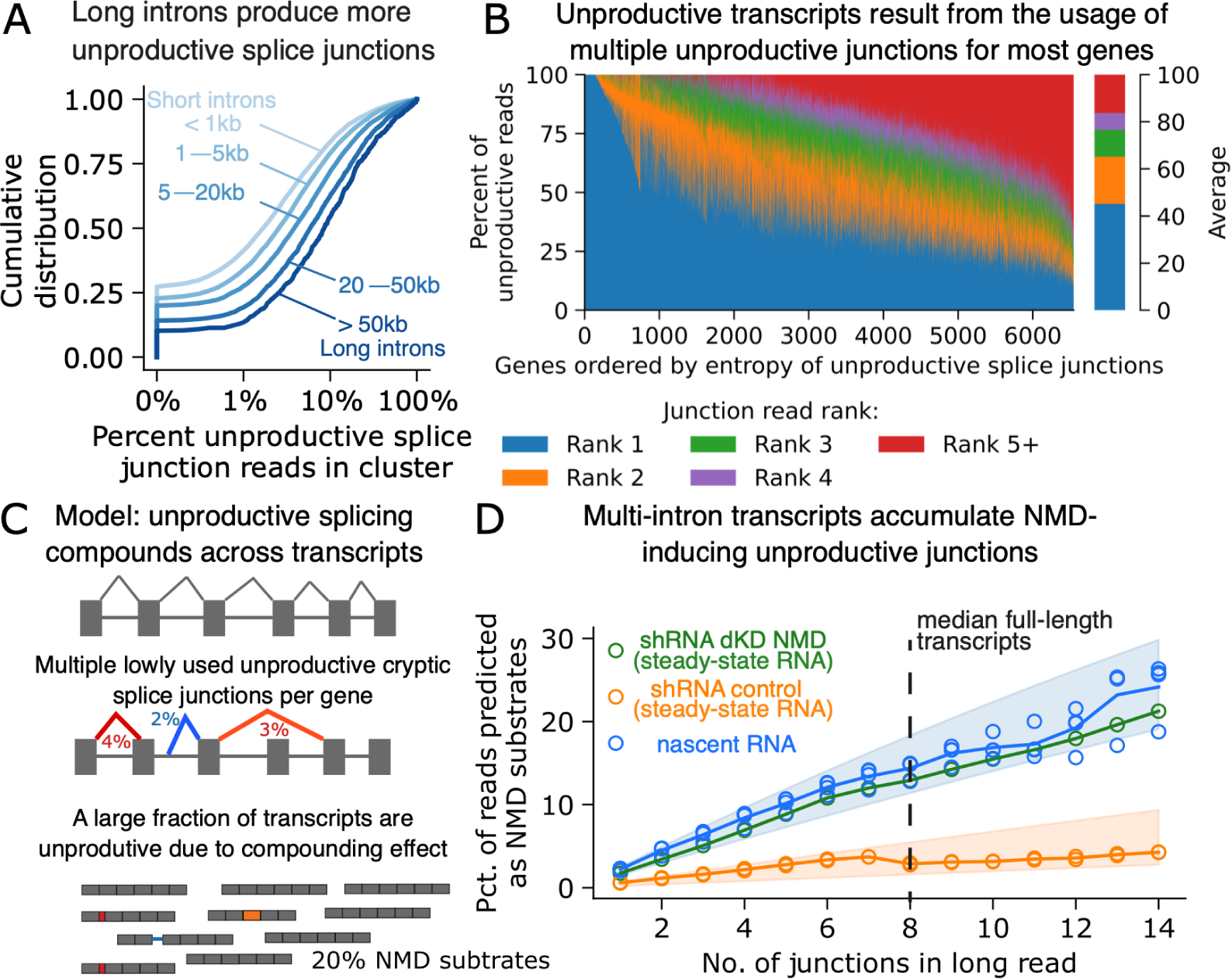
Unproductive mis-splicing accumulates across transcripts. (A) Correlation between percent of unproductive junction reads in LeafCutter clusters versus length of the most used productive intron in the cluster. Spearman correlation coefficient of 0.19 (p-value of 1.3×10^-243^). Correlation presented as cumulative distribution of percent unproductive splice junctions, for increasing groups of intron length. (B) Percent unproductive splice junctions for each gene that are attributed to the most common (Rank 1) unproductive junction, the second most common unproductive junction, etc… (C) Model of unproductive splicing compounding across multi-intronic transcripts. A low unproductive rate at many independently spliced introns produces a high rate of unproductive molecules at the transcript level. (D) Nanopore long-read sequencing quantifies the percent of full-length reads that are targeted by NMD, defined as containing at least one unproductive junction, as a function of the number of splice junctions in the read. Vertical dashed line marks 8 splice junctions, corresponding to a typical full-length human transcript. Total RNA isolated from shRNA-mediated double knockdown (dKD) of NMD factors *SMG6* and *SMG7* or shRNA scramble control^29^ in HeLa cells. naRNA data from K562 cells^48^. Multiple points of the same color indicate replicate experiments. Blue and orange shaded area represents the binomial expectation when assuming 1.5-2.5% and 0.2-0.7% of unproductive junction reads at each independent junction.

Based on these observations, we propose a model in which every protein coding intron of a gene can generate unproductive junctions at a specific “error” rate, which depends on intron length, splice site strength, and other genomic features. This model predicts that unproductive junctions will compound along the entire length of pre-mRNA, making long genes – or genes with many introns – likely to produce more unproductive transcripts than those with fewer introns (**Figure 2C**). Under the simplest binomial model, assuming that 1.5-2.5% of splicing events are unproductive, we predicted that 11–18% of mRNA transcripts would be unproductive for a typical human gene with 8 introns. To test this prediction, we analyzed long-read RNA sequencing (LRS) data from datasets collected using Oxford Nanopore technology^29,48^. We investigated the number of transcripts containing one or more unproductive junctions and only considered LRS junctions that were also detected in short-read RNA-seq to rule out any bias caused by sequencing errors from LRS. Additionally, since most reads from LRS do not span the entire length of an mRNA transcript, we binned LRS reads by their number of junctions, and then calculated the fraction of reads with an unproductive junction for each bin to estimate the abundance of unproductive transcripts for genes with different numbers of introns. We first analyzed LRS naRNA from chromatin cell fractions from K562 cells^48^. We found that ∼15% of reads spanning a typical human gene contained one or more unproductive junctions (**Figure 2D**). Importantly, we next investigated LRS of steady-state mRNA following shRNA knockdown of NMD machinery^29^. Again, we find that single NMD factor knockdowns displayed modest increases in unproductive transcripts (**Figure S6B**) compared to double knockdown of *SMG6* and *SMG7*, which revealed that ∼15% of reads spanning a typical human gene had one or more unproductive junctions. This estimate is far above the ∼5% observed in control experiments and is remarkably consistent both with the estimate from LRS of chromatin mRNA and our prediction based on a simple binomial model of compounding splicing errors. Taken together, we estimate that ∼15% of all transcript molecules generated are NMD targets, though rates of unproductive splicing vary greatly across genes. For example, we estimate that 25% of mRNA transcribed from transcripts with 10 exons and over 50% of transcripts with 15 exons or more at lowly expressed genes (bottom quartile of protein coding genes) are NMD targets (**Figure S10**).

### Pervasive effects of sQTLs on gene expression levels

The prevalence of unproductive splicing, along with the observation that unproductive splicing anti-correlates with gene expression levels genomewide, predicts that genetic effects on RNA splicing would often impact RNA expression levels. To test this prediction, we used quantitative trait loci (QTL) mapping (**Figure 3A**) to identify genetic variants associated with expression (eQTLs) and splicing (splice junction abundance, sQTLs) in naRNA-seq, 4sU-seq and steady-state RNA-seq data. To better distinguish splicing-mediated expression effects from transcriptional effects, we mapped histone QTLs (hQTLs), reflective of variants impacting promoter and enhancer activity (H3K27ac, H3K4me1, and H3K4me3) and transcription across gene bodies (H3K36me3). In total, we identified 57,981 QTLs for 620,020 tested molecular traits (**Figure S11**). Consistent with previous work, we find a large fraction of eQTLs are explained by transcriptional regulation, as indicated by the high degree of sharing between eQTL and hQTL signals. For example, we estimate that 67% of eQTLs (steady-state RNA) contain hQTL effects at the gene’s promoter (Storey’s *π*_1_ =0.67, Figure S12A), consistent with transcriptional regulation. As expected, we observe a strong concordance in the direction of hQTL and eQTL effects, wherein hQTL alleles that increase H3K27ac signal at the promoter overwhelmingly have corresponding up-regulating signals at the level of mRNA, as measured in naRNA, 30min or 60min labeled 4sU RNA, or steady-state RNA **(Figure 3B**). The remaining steady-state eQTLs that do not have hQTL signal likely function during or shortly after transcription, as the effects of an additional 24% of steady-state eQTLs are also detected in 30min 4sU-labeled RNA (*π*_1_=0.91, **Figure S12A**). Thus, we estimate that at least 24% of eQTLs function post-transcriptionally. This estimate is robust to various statistical thresholds for identification of eQTLs (**Figure S12**), though we acknowledge that interpretations of QTL sharing may be overestimated if distinct signals are in linkage disequilibrium.

**Figure 3:**
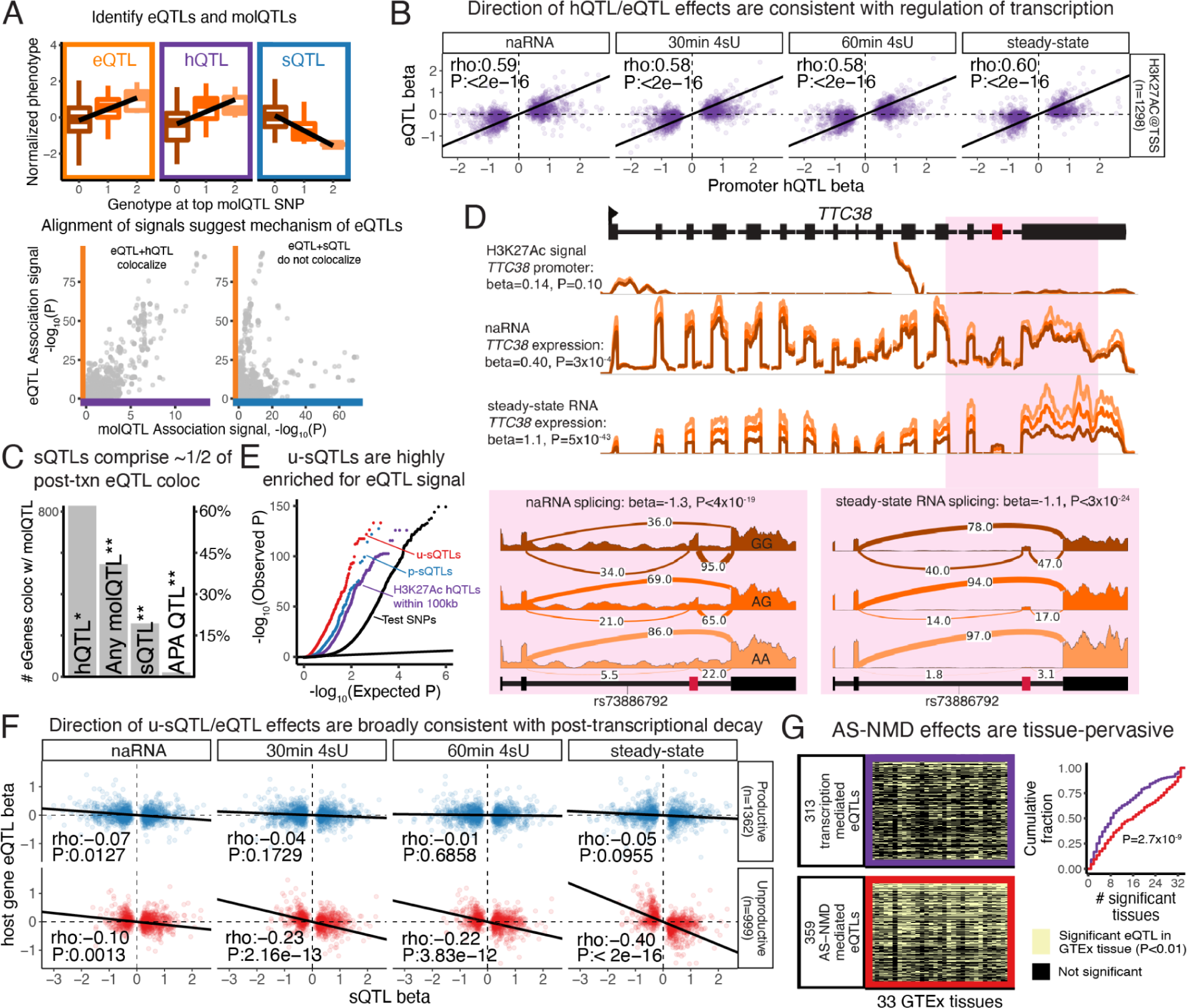
Genetic variants that alter expression post-transcriptionally are enriched in splice-altering variants that are predicted to induce NMD. (A) Approach for identifying mechanisms of gene regulation, using data from the *CCHCR1* locus as an illustrative example. Genetic variants that associate with chromatin peak height, RNA expression, splicing, etc… (molecular QTLs, or molQTL) are identified (boxplots). Multi-trait colocalization compares alignment of molQTL signals (scatter plots) to identify molQTLs that likely share a causal SNP. (B) Effect size (beta) of H3K27ac hQTLs at promoter is correlated with eQTL effect sizes. eQTL effect sizes estimated in each RNA-seq dataset relative to lead hQTL SNP. Correlation summarized with Spearman’s rho coefficient and significance test. (C) Tally of eGenes by their colocalizing molQTLs. 831 eGenes colocalize with an hQTL (*, H3K27ac, H3K4me1, H3K4me3, or H3K36me3). 518 eGenes colocalize with other molQTLs (** indicates eGenes that do not colocalize with any hQTL), suggesting post-transcriptional regulation, some of which colocalize with an sQTL, or alternative polyadenylation (apaQTL). (D) Example eQTL for the *TTC38* gene functioning through AS-NMD caused by inclusion of a poison exon (red). H3K27ac ChIP-seq and RNA-seq coverage grouped by genotype of the lead eQTL SNP. Pink inset region shows effect on splicing of poison exon, depicted in detail as sashimi plots with relative usage (intronic PSI) of splice junctions as arcs. (E) eQTL QQ-plot shows inflation of eQTL signal among various groups of SNPs (lead SNPs for p-sQTLs and u-sQTLs within the host-gene, H3K27ac QTLs within 100kb of test gene, or random test SNPs). (F) (Top row) Effect size of p-sQTLs (all sQTL introns in LeafCutter cluster are productive splice junctions) versus the effect on host gene expression (eQTL beta). (Bottom row) Effect size of u-sQTL (sQTLs which significantly influence at least one unproductive splice junction) versus effect on host gene expression. Similar to (B), effects assessed in each RNA-seq dataset relative to the top sQTL SNP, using the unproductive splice junction for u-sQTLs. (G) eQTLs consistent with regulation by transcription (eQTL/hQTLs, purple) in discovery dataset in LCLs were compared to eQTLs consistent with AS-NMD (eQTL/u-sQTLs, red). Left: Each eQTL (SNP:gene pair, rows) assessed for effects in GTEx tissues (columns). Right: Cumulative distribution for number of tissues with significant effects. P value corresponds to a two-sided Mann-Whitney test.

To better interpret molecular mechanisms of eQTLs while accounting for linkage disequilibrium, we performed a multi-trait colocalization analysis to identify molecular QTLs that likely share causal variants (**Figure 3A**, **Figure S13A**, STAR Methods). For example, at the *CCHRC1* locus, we identify an eQTL, an hQTL and an sQTL (**Figure 3A**). While both the sQTL and hQTL are nominally significant for an eQTL association, colocalization analysis reveals that only the hQTL and eQTL are likely caused by the same genetic variant (posterior probability of full colocalization, PPFC > 0.5). This suggests that transcriptional regulation explains the eQTL effect, while the sQTL constitutes a distinct genetic effect that does not explain the primary eQTL signal. Across all 3,970 steady-state RNA eQTLs, we find 831 (∼20%) colocalized with an hQTL (PPFC>0.5) (**Figure 3C**, **Figure S13A**). We do not interpret this to suggest that only 20% of eQTLs are caused by transcriptional regulation, but rather that 20% represents a lower limit of the degree of transcriptional regulation. Nevertheless, we reasoned that the abundance of eQTL colocalizations with hQTLs relative to their colocalizations with other molecular QTLs may be informative. For example, when considering eQTLs that colocalized with any molQTL we tested, 62% colocalized to an hQTL. Among the remaining 38%, which we hereafter refer to as “post-transcriptional eQTLs”, nearly half colocalized with an sQTL (**Figure 3C**, **Figure S13B**) suggesting that AS may be a major contributor to inter-individual variation in gene expression levels. By comparison, alternative polyadenylation QTLs (apaQTLs) colocalized with only ∼5% of the non-hQTL colocalizations (**Figure 3C**, **Figure S13B**), suggesting that alternative polyadenylation plays a comparatively minor role.

The degree of sQTL-eQTL colocalization suggests that a large number of genetic variants may impact gene expression levels through primary effects on AS. However, we and others have noted that alternative transcription initiation or polyadenylation can alter splicing quantifications, and vice versa^42,52–54^. Furthermore, preferential decay of specific mRNA isoforms may manifest as sQTL-eQTL colocalizations^55^, without being mediated by AS. To better assess whether splicing changes causally drive the observed abundance of eQTL-sQTL colocalizations, we interrogated the genetic variants underlying post-transcriptional eQTLs. Specifically, we asked which genomic annotations are most enriched among post-transcriptional eQTL SNPs, compared to transcriptional eQTLs as controls. As expected, post-transcriptional eQTLs are strongly depleted in enhancers and promoters. In terms of enriched genomic regions, we find significant enrichment of post-transcriptional eQTLs near polyadenylation sites, and in splice donor, branch site, and splice acceptor regions (**Figure S13C**). These observations indicate that changes in RNA splicing causally drive many of these post-transcriptional eQTLs.

As an illustrative example of a splicing-mediated eQTL, we highlight the *TTC38* gene (**Figure 3D**) for which the lead eQTL variant has no detectable effect on promoter activity. Rather, we observed a clear effect of the lead eQTL variant on splicing. The allele associated with decreased expression level is also associated with an increase in splicing of unproductive introns. These unproductive splice junctions are substantially more abundant in naRNA than steady-state RNA, consistent with their rapid degradation, though the sQTL effect size is similar in naRNA and steady-state RNA (**Figure 3D**). More generally, when we stratify sQTLs by whether they affect an unproductive splice junction (henceforth “u-sQTLs”), versus merely switching between alternate productive protein coding isoforms (henceforth “p-sQTLs”), we find that u-sQTLs are particularly enriched in eQTL signal for the host gene (**Figure 3E, Figure S14A**), with an even stronger eQTL signal than H3K27ac QTLs within 100kb of the gene (**Figure 3E**). Furthermore, though we identify a similar number of p-sQTLs and u-sQTLs overall, u-sQTLs explain 77% of sQTL colocalizations with post-transcriptional eQTLs, compared to 23% for p-sQTLs (P=5.9×10^-12^, hypergeometric test, **Figure S14B**). These results again suggest that AS-NMD is a major contributor to the regulation of gene expression levels.

To further validate that u-sQTLs affect gene expression levels through post-transcriptional decay, we considered the concordance of effect sizes between u-sQTLs and eQTLs. As gene expression traits often contain multiple independently acting eQTL signals, we considered every detected lead sQTL SNP and assessed its eQTL effect size, rather than considering only the top sQTL or top eQTL per gene. To remove possible confounding effects from alternative promoter usage, which may manifest as both an sQTL and an eQTL, we filtered out sQTLs that have significant hQTL signal from this analysis (STAR Methods). As expected, p-sQTL variants are largely inert with respect to expression levels. In contrast, u-sQTL effects are strongly anti-correlated with expression, such that alleles increasing the unproductive junction are associated with decreased expression of the host gene (**Figure 3F**, steady-state facet). These effects on expression are most apparent in steady-state RNA, weaker in our 4sU RNA-seq data, and largely absent in naRNA (**Figure 3F**), which is expected as NMD occurs post-transcriptionally and in the cytoplasm. Though we identify similar numbers of p-sQTLs and u-sQTLs overall, u-sQTLs tend to have stronger and more predictable effects on mRNA levels (**Figure 3F**).

While we observed an overwhelming enrichment of eQTLs among u-sQTL SNPs (**Figure 3E**), previous studies have found that the leading eQTL and sQTL signals for a given gene tend to be independent. To reconcile these observations, we provide two mutually compatible explanations. Firstly, we note while many genes have multiple sQTLs (**Figure S15B**), the strongest sQTL for a gene is often a p-sQTL, which is generally not expected to trigger NMD. In contrast, u-sQTLs are less likely to be the strongest sQTL for the gene (**Figure S15C**, Odds ratio 1.4, P=6.1×10^-11^, hypergeometric test), likely because the unproductive isoform is rapidly degraded to a low abundance in steady-state RNA. Secondly, we note that genes often have multiple detectable independent eQTL signals that contribute to the gene’s regulation. Thus, while the lead eQTL is more often explained by transcription-based mechanisms (**Figure 3C**), sQTLs may still pervasively contribute to gene regulation as secondary or tertiary eQTLs.

Given previous reports that lead eQTLs can sometimes change across tissues^56,57^, and our previous observation that sQTL effects are generally more stable across tissues than eQTLs^58^, we wondered about the tissue-pervasiveness of u-sQTL effects on expression. We hypothesized that the effects of splicing-mediated eQTLs would be more consistent across tissues than the effects of transcription-mediated eQTLs. To test this, we assessed the eQTL effects of u-sQTLs across a wide range of tissues collected through the GTEx consortium. For comparison, we also assessed eQTL effects across tissues for the hQTLs identified in our data (**Figure 3G**). Consistent with cell-type-specific activity of enhancers, we find that eQTLs mediated by splicing mechanisms are significant eQTLs in a larger number of GTEx tissues than eQTLs mediated by transcription regulation (**Figure 3G**). Accordingly, sQTL/eQTLs also have a larger median effect size and smaller standard deviation of effects across GTEx tissues, despite having slightly weaker effects than hQTL-eQTLs in our discovery dataset (**Figure S16**). Thus, the regulatory impact of variants that function through AS-NMD are more tissue-pervasive than those that function through transcription. Such variants could be relevant for interpreting genome-wide-association (GWAS) signals that have yet to be explained by eQTLs in known tissue or cell-types.

### Phenotypic impact of AS-NMD

Given the pervasive effects of AS-NMD on gene expression, we wondered to what extent the AS that affects organism-level phenotypes are mediated by NMD, versus the protein-diversifying function of AS. Previous works have identified examples of trait-associated variants which are clearly mediated by AS-NMD. For example, GWAS variants that affect ALS risk and COVID-19-susceptibility increase the inclusion of a cryptic exon in *UNC13A* and *OAS1*, respectively, leading to lower protein expression^59–61^. More generally, recent studies^62–65^ have found that approximately half of GWAS loci that share a causal variant (colocalize) with splicing quantitative trait loci (sQTLs) also colocalize with an eQTL, suggesting that the impact of AS is often mediated by gene expression levels rather than alternate protein isoforms. However, the overlap of sQTLs, eQTLs, and GWAS does not immediately imply AS-NMD mechanisms because alternative promoters, isoform-specific mRNA decay, and other molecular processes can similarly affect both expression and splicing measurements^52–55^. To better resolve how these sQTLs may be functioning, we compiled summary statistics from 45 GWAS for blood and immune-related traits and evaluated the enrichment of GWAS signal among various classes of molecular QTLs: eQTLs, H3K27Ac QTLs, p-sQTLs, and u-sQTLs For example, using multiple sclerosis GWAS summary statistics, we find enrichment of u-sQTLs on par with that of p-sQTLs, hQTLs, and eQTLs (**Figure 4A**). Similar results are found in most complex traits we examined (**Figure S17**).

**Figure 4:**
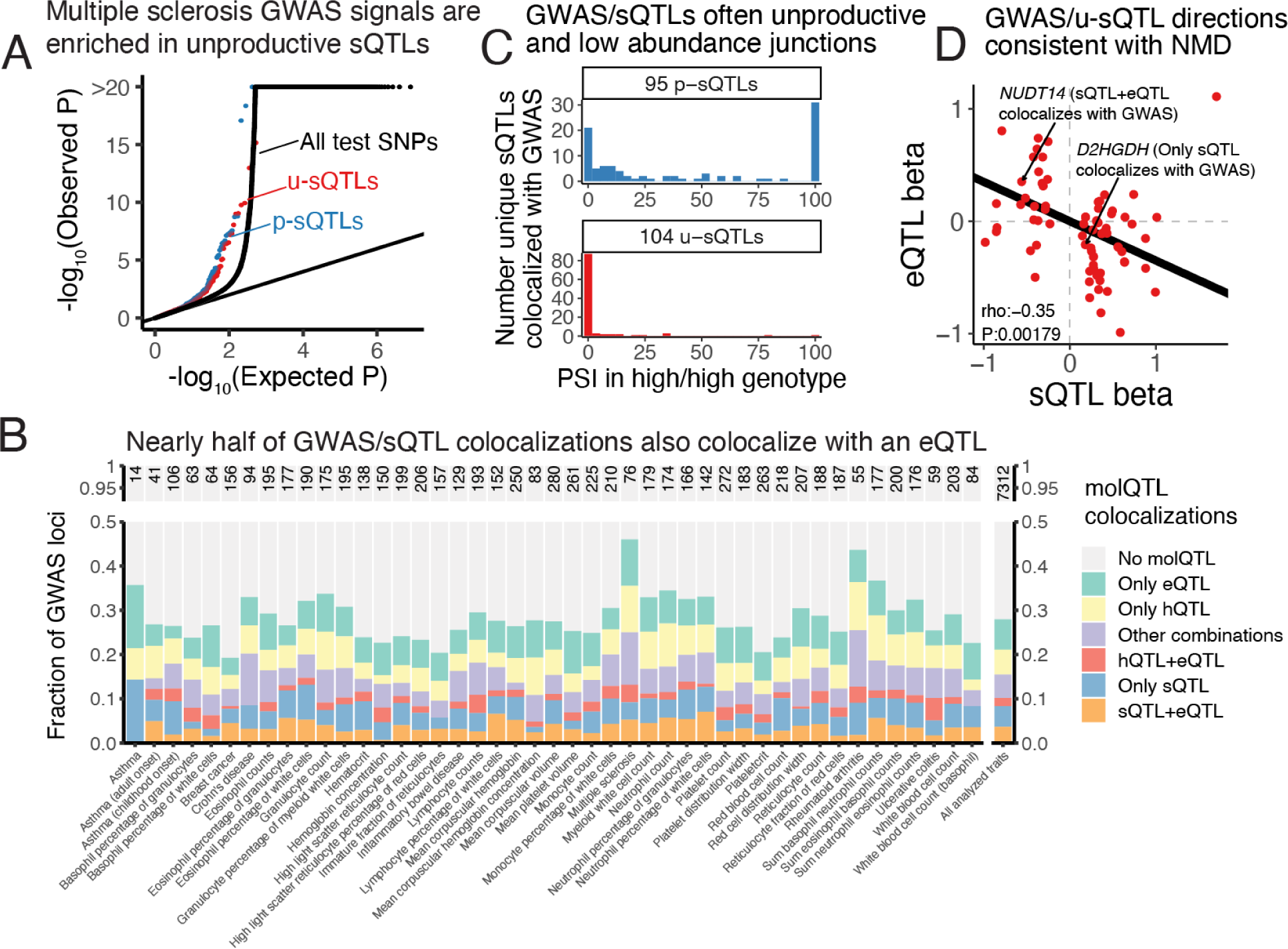
Splicing-mediated NMD contributes to complex trait biology. (A) QQ-plot of multiple sclerosis GWAS signal, grouped by categories of SNPs. p-sQTLs that impact the balance of protein coding isoforms and u-sQTLs that impact usage of unproductive splice junctions are similarly inflated for GWAS signals. (B) Fraction of GWAS loci that colocalize with various sets of molecular QTLs (molQTLs) in each of 45 blood or immune-related traits. Number of loci for which colocalization was attempted is indicated at the top of each column. “Other combinations” includes loci that colocalize with alternative polyadenylation QTLs, or hQTLs and sQTLs, or other combinations which may include sQTLs and other molQTLs, and are difficult to interpret mechanistically. (C) Histogram of usage of unique sQTL junctions that colocalize with a GWAS signal, grouped by sQTL type. Intronic PSI (junction read count divided by most abundant junction in LeafCutter cluster) for each junction was summarized as the median from steady-state RNA samples which are homozygous for the PSI-increasing allele. (D) Effect size (beta) of sQTLs and eQTLs for distinct u-sQTLs that colocalize with a GWAS signal. Correlation summarized with Spearman’s rho coefficient and significance test.

To better resolve the mechanisms at these GWAS loci, we used multi-trait colocalization to identify GWAS signals that colocalize with hQTLs, eQTLs, sQTLs, or various combinations of QTLs. Across all complex traits, approximately 70% of GWAS loci could not be colocalized with any molecular QTL (**Figure 4B**), in line with previous studies with similar sample sizes^64,65^. Approximately 18% of GWAS loci colocalize with either an hQTL, eQTL, or some combination of molecular QTLs (**Figure 4B**). The remaining 12% of loci colocalize with an sQTL, but not an hQTL, consistent with splicing-mediated impacts on traits. We next sought to assess whether these GWAS/sQTL loci possess characteristics consistent with an AS-NMD mechanism, versus protein-diversification.

Notably, we find that these sQTLs largely affect low-usage splice junctions, such that 57% of these splice junctions are spliced-in at PSI<5% (**Figure 4C**). This observation is consistent with that of a recent study^64^ from the HipSci consortium, and naturally poses questions as to how such low-usage isoforms might impact traits. We found that most of these low-usage sQTLs are u-sQTLs that alter the balance of unproductive and productive isoforms, consistent with AS-NMD mechanisms mediating these loci. We also found that the allelic effects of u-sQTLs that colocalize with GWAS loci, like that of u-sQTLs in general (**Figure 3F**), were anti-correlated with their effects on gene expression levels, again consistent with NMD (**Figure 4D**). These u-sQTLs are also enriched among the GWAS/sQTL loci that also colocalize with eQTL signal (**Figure S18A**). For example, a reticulocyte-count-associated GWAS signal colocalizes with a u-sQTL in *NUDT14* gene, as well as *NUDT14* eQTL signal (**Figures S19A**,**B**). As expected, the allele that increases usage of the unproductive splice junction is associated with *NUDT14* downregulation (**Figure 4C**). While the effect on splicing is present in both steady-state RNA-seq and naRNA-seq (**Figure S19C**), the effect on *NUDT14* expression is only apparent in steady-state RNA (**Figure S19D**), again consistent with AS-NMD.

More generally, u-sQTLs that colocalize with GWAS and eQTL signals tend to also display lower usage in steady-state RNA than naRNA (**Figure S18B**), with sQTL and eQTL effects that are also consistent with AS-NMD (**Figure S18C**). In contrast, the sQTLs that colocalize with GWAS but not eQTL do not share these characteristics (**Figures S18B,C**), and likely function by tuning the expression levels of alternative protein coding isoforms. Given that there are a similar number of these GWAS loci that colocalize with both sQTL and eQTL (and not hQTL), as compared to just sQTLs (**Figure 4B**), we conclude that AS-NMD carries similar importance as splicing-mediated protein diversification for complex organism-level traits.

While the sQTLs that colocalize with GWAS but not eQTL signals may generally function by impacting the balance of protein coding isoforms, we highlight an interesting exception which does show characteristics consistent with AS-NMD. An asthma-associated GWAS signal colocalizes with a u-sQTL in the *D2HGDH* gene (lead variant rs34290285), for which the protective-allele produces an unproductive isoform with a PTC (**Figures S20A**,**B**), and subsequent downregulation of *D2HGDH* expression levels (**Figure 4D**). Although this is consistent with AS-NMD, we found no GWAS/eQTL colocalization at this locus. Inspection of colocalization scatter plots of eQTL, sQTL, and GWAS signals revealed a stronger, independent eQTL signal (lead variant rs71430382) that is well aligned with weaker GWAS signals but confounds GWAS-eQTL colocalization (**Figure S20C**). Given the general tissue-pervasive effects of AS-NMD mediated eQTLs compared to transcription mediated eQTLs (**Figure 3G**), we wondered about the relative effects of these eQTLs in different tissues. Indeed, across GTEx tissues, we find other tissues, such as thyroid, where rs34290285 eQTL signal is as strong or stronger than rs71430382 (**Figure S20D**), raising the possibility that the AS-NMD-mediated rs34290285 eQTL has the largest impact on *D2HGDH* expression levels in cell-types most relevant to asthma. Accordingly, colocalization scatter plots better support convergence of *D2HGDH* eQTL and GWAS signals in thyroid (**Figure S20E**). Thus, even some of the low-usage sQTLs that that colocalize with GWAS signals but not eQTL may still function through AS-NMD.

### Broad impact of AS on gene expression levels revealed by therapeutic perturbation

Our findings collectively suggest that most AS events result in unproductive transcripts that are targeted by NMD for rapid degradation. To validate this finding further, we measured the effects of a non-specific splice-switching drug on gene expression levels in LCLs (**Figure 5A**). Risdiplam, a small molecule used to treat spinal muscular atrophy, increases the usage of non-canonical GA|GU 5’ splice sites (5’ss) by stabilizing the spliceosome at U1:pre-mRNA duplexes with bulged adenosines at the −1 position in the 5’ss^66–68^. At low doses, risdiplam selectively promotes the inclusion of exon 7 into *SMN2*, allowing its protein to compensate for SMN1 loss-of-function. Recent studies^68,69^ find that at higher doses, risdiplam’s effects are less selective, extending to GA|GU 5’ss in many other genes. We predicted that a risdiplam-induced increase in noncanonical GA|GU splice site usage would lead to an abundance of unproductive, NMD-targeted transcripts and consequently, decreased expression of the affected genes.

**Figure 5:**
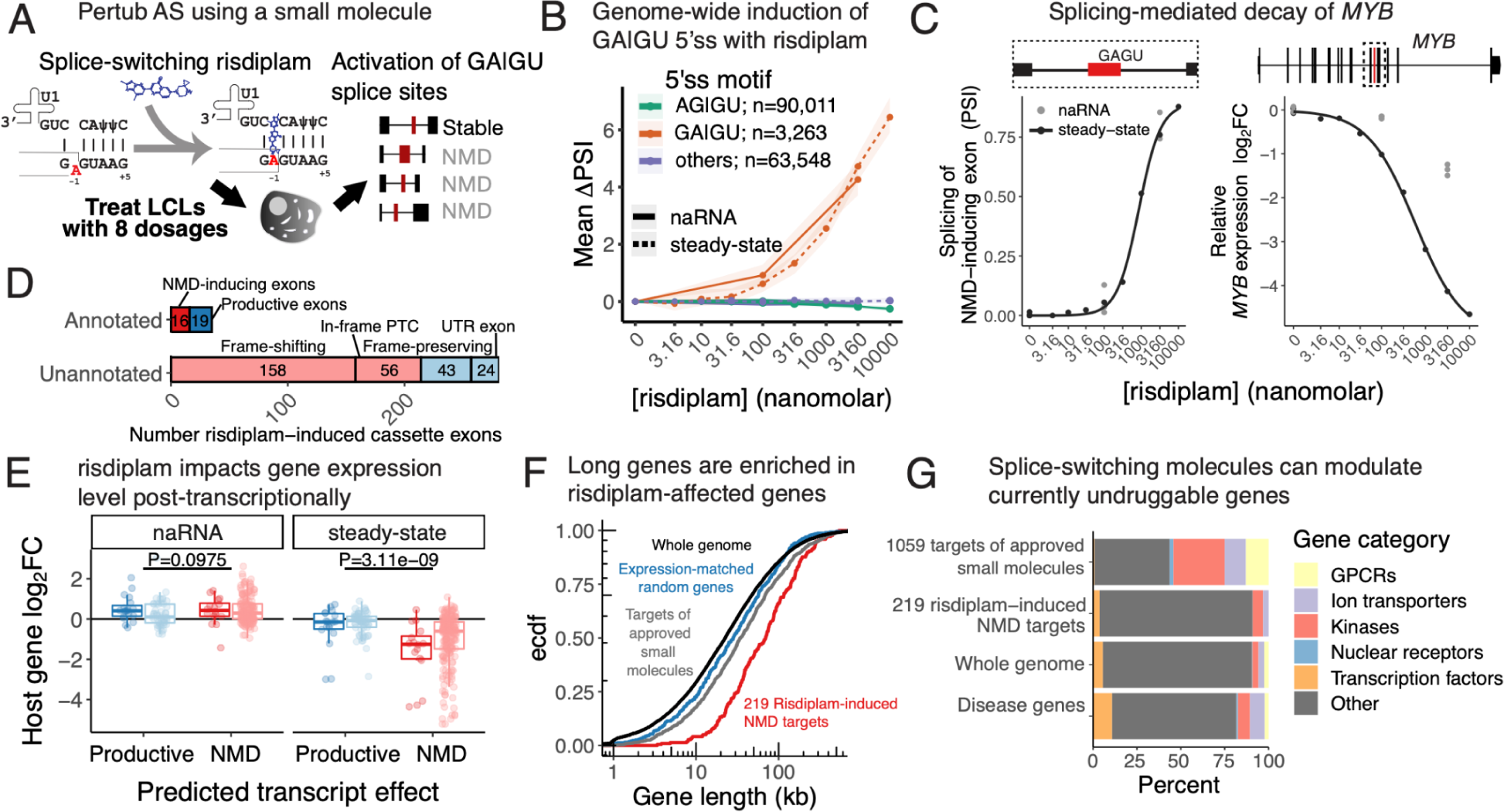
Risdiplam-induced splicing alterations mediate expression changes at hundreds of genes. (A) Overview of risdiplam-based approach to assess pervasiveness of NMD after splicing perturbations. LCLs treated with 8 doses of risdiplam. Splicing changes at cryptic exons assessed for NMD-potential, and gene expression changes were estimated. (B) Genome-wide mean splicing dose-dependent changes at various classes of 5’ss in naRNA and steady-state RNA. Bootstrapped 95% confidence intervals shaded around the mean activation level across n introns in each group. (C) Left: Dose-dependent splicing response (left) at a risdiplam-targeted exon in *MYB*. Right: Dose-dependent expression response of *MYB*. (D) Predicted translation result of 305 risdiplam-induced exons. Exons expected to induce NMD versus those that maintain transcript stability are red and blue, respectively. Annotated and unannotated exons are dark and light colors, respectively. (E) Empirically measured effect of host gene expression as measured in steady-state RNA and naRNA in the presence of risdiplam at 3.16*μ*M. Each point is an induced exon/host-gene, colored the same as in (D). (F) Cumulative distribution of gene length for genes with predicted NMD-induced exons, a similarly sized set of expression-matched genes, all genes, or a set of gene targets for FDA-approved small molecules. (G) Conventional small molecule drug targets are disproportionately skewed for particular classes of ‘druggable’ genes (e.g. G-protein-coupled receptors, GPCRs) that operate at the level of protein binding. Risdiplam-induced NMD targets are more representative of all genes. Disease genes with therapeutic potential by down-regulation (OMIM dominant negative genes) are similarly distributed across categories of previously ‘druggable’ genes.

To test this prediction, we treated LCLs with risdiplam and sequenced steady-state RNA at 8 increasing doses of risdiplam, and naRNA at 2 doses. We observed widespread, genome-wide induction of GA|GU 5’ ss in a dose-dependent manner, but no effect on canonical AG|GU 5’ss or other non-canonical splice sites (**Figure 5B**). In total, 641 out of 3,263 observed non-canonical GA|GU splice sites were significantly activated with a dose-dependent response (FDR<10%, **Figure S21**). Of those, 305 were involved in inclusion of cassette exons (hereafter referred to as risdiplam-induced exons), as opposed to other forms of AS. For example, *MYB*, a transcription factor proto-oncogene targeted by a number of anti-tumor therapies, contains a risdiplam-induced exon (**Figure 5C**). This exon is uniquely annotated in an unproductive transcript isoform, and *MYB* expression is down-regulated in a dose-dependent manner. As expected, the decrease in *MYB* expression is more pronounced in steady-state RNA than in naRNA, consistent with splicing-induced post-transcriptional decay.

Unlike the exon in *MYB*, 267/305 (88%) of all risdiplam-induced exons are unannotated. We used the annotated productive transcript structure as a reference to classify whether these unannotated exons are predicted to introduce PTCs and trigger NMD. We found that 246/267 (92%) of the unannotated risdiplam-induced exons are within the coding region of the transcript (i.e., not in UTRs). Of these, 158/246 (64%) result in a frameshift, matching our naïve expectation that 2/3 of random splicing changes should produce a frameshift. An additional 48/246 (19%) of the novel exons contain an in-frame PTC, and are also expected to induce NMD. The remaining unannotated exons preserve the reading frame and are not expected to induce NMD (**Figure 5D**). Overall, the effects of these predicted stable- and NMD-inducing-exons on gene expression matched our expectations. That is, inclusion of predicted frame-shifting- and in-frame-PTC-containing exons results in down-regulation of the affected gene in steady-state steady RNA, but not naRNA (**Figure 5E**, **Figure S22**). While it is possible for splice junctions in the 3’UTR to also induce NMD, provided that the UTR splice junction complex is sufficiently downstream of the stop codon, we note that all of the UTR exons induced by risdiplam in this study emerged at UTRs that already contain a splice junction in the annotated transcript structure. These additional UTR junctions generally do not appear to induce additional decay (**Figure S22**).

As it is well-established that differential splicing analyses are limited in power at currently achievable sequencing depths compared to differential expression analysis^70–72^, we wondered whether we could find additional AS-NMD effects by examining gene-expression changes for genes in which we could not detect differential splicing. When considering all genes genome-wide, we find that risdiplam overwhelmingly induces down-regulation rather than up-regulation of genes in steady-state RNA (**Figure S23A**). Though we have limited power to identify the unproductive splice junctions underlying these changes, we surmise that hundreds of additional genes are subject to similar induction of splicing-mediated decay. To further explore this possibility, we sought to determine whether the relative abundance of down-versus up-regulated genes is due to post-transcriptional mechanisms, which would be consistent with AS-NMD. We compared risdiplam-induced expression changes in naRNA to risdiplam-induced changes in steady-state RNA. Given that risdipam’s known direct molecular targets involve splicing machinery^67,68,73^, we reasoned that changes in naRNA reflect transcriptional changes that result from secondary effects of the drug. In contrast, changes in steady-state RNA should reflect the sum of transcriptional and post-transcriptional changes. Guided by this intuition, we classified risdiplam-induced expression changes as transcriptional, post-transcriptionally down-regulated, or post-transcriptionally up-regulated. We find nearly twice as many post-transcriptionally down-regulated genes as post-transcriptionally up-regulated genes (**Figure S23B**). Post-transcriptionally down-regulated genes are greatly enriched for genes with predicted NMD-inducing exons, which we identified through independent analysis of splicing effects (**Figure S23B**, Odds Ratio=14.0, P=2×10^-16^). While splicing perturbations can also promote productive isoforms at the expense of unproductive isoforms, resulting in up-regulation of the affected gene as in the case of *SMN2*^66^, our data suggests that random splicing perturbations, such as those introduced by non-specific splice-switching drugs or *de novo* mutations, are more likely to result in NMD-targeted transcripts and subsequent down-regulation.

Splice-switching drugs such as risdiplam and the antisense oligo nusinersen have proven that AS has great potential as a therapeutic target^74–78^. Genes that are subject to splicing-induced down-regulation may be particularly amenable as targets of splice-switching drugs. We therefore sought to describe the characteristics of genes that were targeted by risdiplam for downregulation. In accordance with our previous observation that unproductive junctions accumulate across the length of the transcript, and that long introns and long genes are most susceptible to AS-NMD, we found that the 214 genes we identified with risdiplam-induced NMD-exons tend to be longer and contain more exons (median gene length 60kb, median of 14 exons) than a set of expression-matched random genes (median gene length 20kb, median of 9 introns, **Figure 5F**, **Figure S24A**). Similarly, the introns that host these risdiplam-induced NMD-exons tend to be longer than other introns within the same set of genes (**Figure S24B**). Unlike most existing small molecule drugs, which function by protein binding and are highly enriched for a small subset of ‘druggable’ protein families (e.g., G-protein coupled receptors, kinases, etc.), risdiplam targets are not strongly enriched for any particular gene families (**Figure 5G**, **Figure S25**). Rather, we find that risdiplam targets are more or less random, with the exception that long genes are more likely than short genes to contain a cryptic splice site (**Figure 5F**, **Figure S24**, **Figure S25**). This suggests that risdiplam and related splice-modulators could be leveraged to target ‘undruggable’ disease genes. To nominate candidate gene targets for therapeutic treatment by close derivatives of risdiplam, we curated a list of disease genes from Online Mendelian Inheritance of Man (OMIM) database that contain dominant negative alleles, as these may have therapeutic potential when down-regulated. We identified 11 disease genes with an identifiable NMD exon that are down-regulated by risdiplam (**Figure S26**). Among these, we recovered the *HTT* gene, for which dominant mutations cause Huntington’s disease^79^. Several experimental splice-modifying drugs targeting *HTT* are currently being investigated as potential treatments in pre-clinical and clinical trials^76,77,80^. Among the other candidate targets are transcription factors and other protein families for which drug development by protein-binding small molecules is notoriously difficult, suggesting that perturbation of AS-NMD greatly expands the utility of splice modulators for treating human diseases.

## Discussion

The molecular impact of AS has been challenging to verify experimentally, as it has been difficult to study the function of individual isoforms at the protein level, and the rapid decay of unproductive isoforms obscure their quantification at the mRNA level. Through detailed analysis of molecular measurements that capture the major steps of RNA maturation, we found that aberrant splicing produces remarkably high levels of unproductive transcripts bearing a premature termination codon. Unproductive mRNAs account for around 15% of all mRNA transcripts from the average human gene, even exceeding 50% for many long genes expressed at low levels. These estimates may be surprising given earlier studies that utilized single knockdown of *UPF1* or *UPF2* to identify a relatively small subset of AS-NMD-regulated genes^24–27^, suggesting that unproductive isoforms are produced at such low rates that expression levels of most genes are unaffected by AS-NMD^25^. However, we find that early estimates are consistent with incomplete inhibition of NMD due to partial redundancy between core NMD factors. Indeed, the levels of unproductive splicing in steady-state mRNA stabilized from double *SMG6* and *SMG7* knockdowns – but not that of single *UPF1* knockdowns – were nearly identical with that estimated in our nascent mRNA dataset. Thus, previous studies using knockdowns of single NMD factors appear to have substantially underestimated the impact of AS-NMD.

Notably, we also show that an important previous observation - that highly used AS exons are enriched for frame-preserving alternate coding isoforms - is largely due to NMD surveillance rather than selection at the level of splicing regulation. Thus, our study suggests that the molecular impact of AS is largely shouldered by NMD, which regulates protein output by targeting unproductive transcripts for degradation. Supporting this view, we identified nearly as many genetic variants that impact production of these unproductive transcripts as compared to those that tune the balance of stable mRNA isoforms. Importantly, this observation also holds for sQTLs that colocalize with GWAS signals to influence organism-level traits. The tissue-pervasive nature of AS-NMD-mediated eQTLs compared to transcription-mediated eQTLs may have particular relevance both in enhancing the phenotypic impact of AS-NMD-based regulation and in mapping the regulatory mechanisms underlying genetic associations for complex traits.

What fraction of mRNA isoforms encode functionally diverse peptides has long been under debate^7,25,36,38,81^. Novel isoforms continue to be discovered as RNA-sequencing experiments increase in depth, making this question particularly timely. Our observation that introns are often mis-spliced into substrates of NMD indicates that splicing is inherently noisy and that novel isoforms uncovered by RNA-seq generally do not encode functional proteins. This view is consistent with previous findings that most AS events are lowly used, and do not show cross-species conservation.

Even for mRNA isoforms that are highly expressed and conserved, it remains unclear how often their primary function is to encode functionally diverse proteins. Previous comparative analyses of AS identified hundreds of exons that are alternatively spliced across mammals, and found that ∼70% of them preserved coding frame^1,2^. While this may suggest that functional AS tends to work through protein diversification, we note two limitations to this interpretation: Firstly, the reliance on steady-state RNA-seq introduces an ascertainment bias against identification of conserved AS-NMD events. And secondly, like other sequence elements that impact gene expression levels (e.g., enhancers, or miRNA binding sites), functionally important AS-NMD splice junctions need not be maintained to optimize gene expression levels, as other cis-elements can compensate for the loss or gain of AS-NMD. In contrast, it is more difficult to envision how protein-diversifying functions can be compensated.

We posit that future research will reveal a preponderance of cases where regulated AS functions by tuning protein expression levels rather than by creating protein diversity, as the sheer abundance of AS-NMD events presents opportunities for evolution to co-opt AS-NMD as a functional regulatory mechanism. Regulated AS-NMD has been identified in the past but it has largely been found in genes encoding splicing regulators. While we confirmed that splicing regulators are enriched among genes with high levels of unproductive transcripts, splicing regulators represent only a small fraction of all genes with very high levels of unproductive transcripts. Although the abundance of AS-NMD does not necessarily imply functional regulation, our observations raise the possibility that functionally regulated AS-NMD may be much more common than was previously appreciated. We predict that future work using long-read sequencing of RNA across multiple species, stages of maturation, and biological systems will provide us with a much more complete understanding of the mechanisms by which AS functionally impacts cellular function, organismal phenotypes, and evolution.

Finally, irrespective of functionality, the existence of multiple cryptic splice sites in the vast majority of human genes implies that there are abundant targets that can be manipulated by splice-switching molecules for therapeutic effects. More research is needed to find out how to activate these cryptic splice sites in a selective manner, but when accomplished, we foresee targeting RNA splicing as a widespread modality by which we can therapeutically control the expression level of disease-causing genes.

## Acknowledgements

We thank members of the Li and Gilad labs for discussions and support. We thank Jonathan Pritchard, Yoav Gilad, Xuanyao Liu, Natalia Gonzales, and Christian Jones for their careful reading of our manuscript and their insightful comments. We thank Alex Ruthenburg and Yichen Hou for sharing their protocol for nascent RNA isolation and performing preliminary experiments with this protocol. This work was supported by NIH grants R01GM130738, R01HG011067, R35GM147498, and W. M. Keck Foundation. The naRNA-seq and H3K36me3 CUT&Tag data have been deposited in the Gene Expression Omnibus (www.ncbi.nlm.nih.gov/geo/) and will be available shortly; other data sources can be found in Figure S1.

## Methods

### Publicly available data

We obtained FASTQ files of standard short read RNA-sequencing data from 465 lymphoblastoid cell lines (LCLs) of samples from the 1000 Genomes project produced by the GEUVADIS consortium (ArrayExpress accession number E-GEUV-1).

We obtained the FASTQ files of short read RNA-sequencing data of shRNA double knockdown of *SMG6* and *SMG7* in HeLa cells, and shRNA controls from a previous study^28^ (SRA accession SRP083135).

### Illumina short read RNA-sequencing data

We aligned the Illumina short read RNA-sequencing datasets from LCL and HeLa cells to the human genome version GRCh38 with Gencode v34 annotation using STAR^82^ version 2.7.7a. We used the same parameters as the ENCODE project for Illumina short read sequencing:

--outFilterType BySJout

--outFilterMultimapNmax 20

--alignSJoverhangMin 8

--alignSJDBoverhangMin 1

--outFilterMismatchNmax 999

--outFilterMismatchNoverReadLmax 0.04

--alignIntronMin 20

--alignIntronMax 1000000

--alignMatesGapMax 1000000

For the LCL lines, we used STAR’s WASP mode^83^ with the genotype data from the VCF files of the 1000 Genomes Project. In addition to the previously outlined parameters, we added the following:

--waspOutputMode SAMtag

--outSAMattributes NH HI AS nM XS vW

After alignment, we filtered the BAM files to retain primarily aligned reads. For the HeLa cell lines, which were mapped with standard mode for STAR, we used samtools with the flag -F256 to remove all non-primary alignments. For the LCL cell lines mapped with STAR WASP mode, we kept the alignments that passed the WASP filtering, by retaining those with the vW:i:1 tag.

### Histone modification ChIP-seq and CUT&Tag data

To analyze ChIP-seq and CUT&Tag data, we mapped the reads to the human genome using HISAT^84^ version 2.2.1. We used the --no-spliced-alignment flag to prevent the insertion of junction reads. For the paired end ChIP-seq datasets (H3K27ac, H3K4me1 and H3K4me3), we used the --no-discordant flag to avoid discordant alignments of the paired reads, and allowed a maximum insert size of 1000 bp. We used Hornet’s find_intersecting_snps.py script to find reads that overlap with SNPs for remapping. These reads were realigned using HISAT with the parameters outlined above, and the reads that mapped to different locations were discarded. For the ChIP-seq datasets, we used MACS2^85^ version 2.2.7.1 to call peaks, using default parameters for paired-end reads. We called narrow peaks for H3K27ac and H3K3me1 data, and broad peaks for H3K4me3.

### Molecular trait quantification

We quantified gene expression and histone modification coverage of H3K27ac, H3K4me1 and H3K4me3 using featureCounts^86^ version 2.0.3. For quantifying gene expression in the LCL RNA-seq datasets, we used the Gencode v34 primary assembly annotation. We used the --ignoreDup flag to ignore duplicate reads, and the --primary flag to limit the counts to primary alignments only. For steady-state RNA-seq and naRNA-seq we used the -p flag for paired end reads. For naRNA-seq data we additionally used the -s 2 parameter to signal that the data is reversely stranded. For quantifying ChIP-seq coverage of histone modifications, we used the peaks called by MACS2 as the annotation for featureCounts. Unlike other histone modifications, H3K36me3 covers the entire gene body instead of being concentrated at peaks. We used bedtools multicov to count the number of H3K36me3 CUT&Tag reads overlapping the entire gene body of the primary assembly of protein coding genes.

To quantify splicing in the RNA-seq datasets, we extracted junction reads by running regtools version 0.5.2^87^ on the filtered BAM files, requiring a minimum intron length of 20 bp. We merged the resulting .junc files into a single database of observed splice junctions and splice junction read counts across all samples. For alternative splicing analysis, we used the .junc files from the LCL samples to obtain intron clusters using Leafcutter’s^58^ leafcutter_cluster_regtools_py3.py script. We used the read counts for each intron to quantify the percent spliced-in (PSI) of each splice junction in a cluster.

To quantify splicing efficiency of introns, we used SPLICE-q^88^ version 1.0.0 to calculate the reverse intron expression ratio (IER) in protein coding genes using the Gencode v34 chromosomal annotation. By default, SPLICE-q uses the highest filtering settings for IER quantification, which selects only introns that do not overlap any exons of the same gene or of any other gene.

### Scoring splice site strengths with MaxEntScan

We scored the 5’ and 3’ splice site strength ends of all splice junctions in our data using maxentpy, the python wrapper of MaxEntScan^89^. For each 5’ splice sites, we used a 9 nucleotide sequence consisting in 3 nucleotides in the exon and 6 nucleotides in the intron at the splice site. For 3’ sites, we used 23 nucleotide sequences, with 3 nucleotides in the exon and 20 nucleotides in the intron.

### Oxford Nanopore Technologies long read RNA-sequencing data

Long-read Oxford Nanopore Technology sequencing data was obtained from published sources. The total RNA data from double knockdown of *SMG6* and *SMG7* in HeLa cells (SRA accession SAMEA8691113), as well as two control experiments (SRA accessions SAMEA8691110 and SAMEA8691111), was obtained from a previous publication^29^. Long-read naRNA-seq data in K562 cells (SRA accession SRP171702) was obtained from Drexler et al^48^. Reads were mapped to the human genome version GRCh38 using minimap 2.24^90^ preset parameters for spliced long reads: -x splice. We used the flag -a to produce BAM files with CIGAR strings for downstream analysis.

We used a custom python script to extract the splice junctions from each read using the CIGAR string from the BAM files produced by minimap2. Long-read sequencing is prone to high rate of sequencing errors and misalignments. Accordingly, we only considered splice junctions that matched splice junctions observed in protein coding genes in our short-read analysis. We removed reads that did not match any junctions. Two out of six of the 4sU naRNA samples from Drexler et al had fewer than 10,000 unique long reads left and were removed from downstream analysis. The remaining naRNA 4sU samples had ∼20,000 unique long reads or more matching at least one short-read junction.

Reads with at least one NMD-associated splice junction were considered NMD substrates. Due to the variable length of long-reads, not all reads cover a full-length transcript. As a result, the lack of NMD-associated splice junctions in one read does not guarantee that the read comes from a protein coding transcript. For this reason, we calculated the percent of transcripts confirmed to be NMD substrates (i.e., the percent of transcripts with at least one NMD junction) for reads with 1 to 14 junctions. For any k number of junctions per read, we bootstrapped the reads with k or more junctions to get an estimate of observed NMD transcripts.

We compared the observed percent of NMD transcripts with the theoretical probability of a read presenting an NMD-associated splice junction. The theoretical probability is:

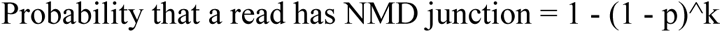

Where k is the total number of junctions in the read, and p is the proportion of splice junctions that are NMD-associated.

### Data normalization

To normalize the LCL RNA-seq data, we first selected the read counts from featureCounts in protein coding genes from Gencode v34’s primary assembly, and for each LCL sample, we kept only the first replicate. We normalized the raw counts into counts per million (CPM) and obtained the log2 of the CPM, defined as:

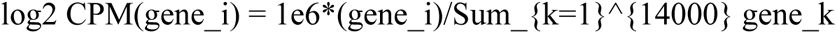

We selected the top 14000 autosomal genes with the highest median CPM in standard RNA-seq data. We utilized these selected genes for all per-gene downstream normalization and analysis for all RNA-seq datasets.

For comparison of expression between the different RNA-seq assays, we applied the log2 of the RPKM normalization on the 14000 selected protein coding genes, using the length of the exonic regions estimated by featureCounts. For H3K36me3, we applied log2 RPKM normalization using the length of the entire gene body. For coverage of the other histone modification marks at the MACS2 peaks, we used the log2 CPM normalization. For alternative splicing, we used the PSI of introns using the rations from LeafCutter without adding a pseudocount.

### Ranking of NMD junctions by contribution and entropy calculation

Out of the 14000 protein coding genes that we selected for downstream analysis, 11563 had NMD associated junction reads in our naRNA-seq data. From these, we selected the 6549 protein coding genes with an average percent of NMD junction reads between 1 and 20% across all naRNA-seq samples. From these, we selected the genes with a total of one hundred or more NMD junction reads aggregated from the 86 naRNA-seq samples (average of at least 1.16 reads per sample). This resulted in a subset of 6537 protein coding genes. For a given gene, we obtained the percent of NMD junction reads contributed by each unique NMD splice junction as follows:

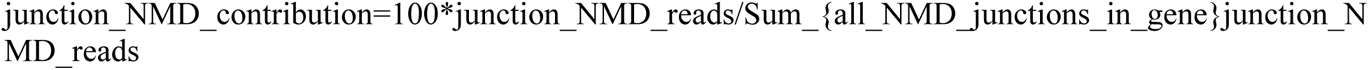

We ranked the junctions by their percent of NMD junction reads contribution. For each gene, we calculated the NMD junction read’s entropy as the Shannon entropy of the fraction of the contribution of NMD reads from each junction:

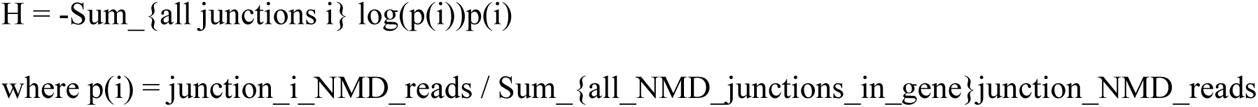

### Analysis of alternative splicing and symmetry of cassette exons

We downloaded the annotation of alternative splicing events for the human hg38 genome from VastDB^91^. We selected cassette exon events that fall within the coding region of protein coding genes. We used VastDB’s annotation of the cassette exons to determine whether each exon is symmetric (i.e., if the length of the exon in bp is a multiple of 3).

For the Illumina short read RNA-seq datasets, we calculated the cassette exon PSI as follows:

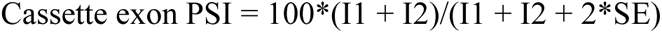

Where I1 and I2 are the splice junction read counts overlapping the first and second splice junction supporting cassette exon inclusion respectively, and SE is the splice junction read counts overlapping the splice junction supporting cassette exon exclusion. For each dataset independently, we retained the cassette exons that have at least one read overlapping any of the three junctions in at least 50% of the samples. To ensure that both junction reads are similarly used in the cassette exons, we discarded the exons in which the average splice junction PSI from I1 and I2 (defined as the average of 100*I1/(I1+I2+SE) and 100*I2/(I1+I2+SE) across all samples respectively; not to be confused with the cassette exon PSI) differs by more than 33%.

For the ONT RNA-seq data, we calculated the cassette exon PSI of each exon as:

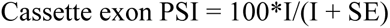

Where I is the total number of reads that contain both splice junctions supporting inclusion of the cassette exon on the same read, and SE is the total number of reads containing the splice junction supporting exclusion of the cassette exon.

We used these cassette exon PSI calculations to estimate the percent of symmetric cassette exons at different PSI ranges.

### Measurements of unproductive junctions in Illumina short-read data

Through this paper we used multiple measurements of alternative splicing for different purposes:

● The **percent of unproductive splice junction reads per gene** correspond to the total number of unproductive splice junction reads in a gene divided by the total number of splice junction reads in the gene.
● The **PSI of a junction at the gene level** is the total number of reads mapping to that junction divided by the maximum number of reads mapping to any junction on the same gene.
● The **PSI of a junction at the Leafcutter intron cluster level** is the total number of reads mapping to that junction divided by the total number of reads mapping to any junction on the same intron cluster.
● The **PSI of cassette exons**, as described in the previous section.
● The **percent of unproductive transcripts** in long read sequencing data, calculated as the percent of transcripts that present one or more unproductive splice junctions.

### Data normalization for QTL mapping

To prepare the LCL RNA-seq data for eQTL mapping, we applied the standard normalization to each gene across all samples. Finally, we enforced normality of the data by applying the rank-based inverse normal transform to each sample across all genes. We utilized the same procedure to normalize the gene coverage quantification for H3K36me3 in the 14000 selected protein coding genes, using the read counts quantified by bedtools multicov.

For ChIP-seq data from H3K27ac, H3K4me1 and H3K4me3, we utilized the read counts quantified by featureCounts at the peaks called by MACS2. If the number of peaks in a dataset exceeded one million, we limited the analysis to the top one million peaks. The rest of the normalization procedure for this data is identical to the normalization that we used for gene expression in RNA-seq data.

To prepare the alternative splicing data in RNA-seq for sQTL mapping, we used the prepare_phenotype_table.py script from Leafcutter. This script selects the intron clusters that have missing data in less than 40% of the samples. For each intron, it then calculates the percent spliced-in (PSI) of each junction while adding a pseudocount of 0.5 to the numerator (total junction read counts in the intron) and to the denominator (total junction read counts in the intron cluster). For each intron, it applies the standard normalization to each gene across all samples. Finally, it enforces normality by rank-normalizing each sample across all introns.

Splicing efficiency ratios were normalized with a custom made script that selects the introns with data missing in less than 40% of the samples and implements the same normalization procedure of applying the standard normalization across the samples followed by rank normalization across the features.

### Principal component analysis for QTL mapping

To build a covariate matrix for QTL mapping for each dataset, we applied principal component analysis (PCA) on the standard and rank-normalized log2 CPM data. We selected the number of principal components that explain more variance than in a version of the data matrix obtained by randomly permuting each feature across the samples. This resulted in 14, 12, 13 and 33 principal components (PCs) for the gene expression data for naRNA-seq, 4sU labeled (30 minutes) RNA-seq, 4sU labeled (60 minutes) RNA-seq, and standard RNA-seq respectively. We also obtained 14, 13, 14 and 11 PCs respectively for H3K27ac, H3K4me1, H3K4me3 and H3K36me3 data. Applying the same procedure on the splicing and IER standard and rank-normalized data, we obtained 11, 8, 8 and 33 PCs for the alternative splicing data matrices of naRNA-seq, 4sU labeled (30 minutes) RNA-seq, 4sU labeled (60 minutes) RNA-seq, and standard RNA-seq respectively; as well as 12, 9, 9 and 24 PCs for IER data.

### Molecular cis-QTL mapping

For molecular cis-QTL calling, we used as input for each dataset the standard and rank-normalized data matrices, the PCs obtained as previously described, and the corresponding VCF files from the 1000 Genomes Project.

We ran QTLTools version 1.3.1 for molecular cis-QTL mapping, using both the permutation and nominal pass versions. For the permutation pass, we used 1000 permutations using the --permute 1000 flag. For the nominal pass, we used the --nominal 1 flag to obtain the QTL statistics of all SNPs irrespective of their p-value. For eQTLs and hQTLs, we used a cis window of 100,000 bp. For sQTLs and splicing efficiency QTLs, we used a cis window of 10,000 bp. For the permutation pass of sQTLs, we used the --grp-best flag to get only the statistics of the best hit per intron cluster. For the permutation pass, we applied the Benjamini-Hochberg correction to the adjusted beta distribution p-values to account for the false discovery rate.

### eQTL calling on GTEx gene expression data

To map eQTLs in GTEx data^57^, we downloaded the publicly available raw count matrices of gene expression in Illumina short read RNA-seq data from the GTEx consortium. We normalized the gene counts for each tissue gene independently, using the standard and rank-normalization of the log2 CPM data described above. For each tissue, we ran QTLtools in nominal pass with the parameters outlined above.

### Nascent RNA-seq experimental methods

#### Cell growth

LCLs were grown in RPMI + glutamine + Penn/Strep + 20% FBS. Cells were grown in 4 batches, with approximately 20-35 cell lines per batch. The day before collection, cell cultures were counted and normalized across cell lines to approximately 35 million live cells, supplemented with media to 50mL, such that cells were in log-phase growth with approximately 50 million cells per culture at the time of collection.

#### Isolation of naRNA

We collected nascent RNA from LCLs by first isolating nuclei through a sucrose cushion followed by high-salt washes to dissociate nucleoplasm and weakly bound RNAs and proteins from chromatin with slight modifications from previous protocols for cellular fractionation^92^, as detailed below.

Cells were collected by centrifugation (300xg for 3min) in 50mL conical tubes. Cells were washed twice in cold PBS with 1mM EDTA. We reasoned that the inclusion of EDTA (and exclusion of magnesium) in buffers used during cellular fractionation would inhibit splicing during sample processing. However, we have noted that chelation of magnesium yields fragile nuclei that are more prone to premature bursting, and care should be taken to pipet gently in subsequent steps. Washed cell pellets were resuspended in 400uL BufferA (10mM pH 7.5 HEPES, 10mM KCl, 10% (v/v) glycerol, 11.6% (w/v) sucrose, 1mM DTT, 1x ROCHE complete protease inhibitor) and transferred to 2mL 96-well plates for convenience. An equal volume of BufferA supplemented with 0.2% TritonX was gently mixed to the resuspended cells, bringing the final TritonX concentration to 0.1% for a 12 minute incubation on ice with periodic inversion. Nuclei were isolated by centrifugation (1200xg, 5min). Supernatant was discarded, and the nuclei pellet was washed with 500uL BufferA, followed by centrifugation (1200xg, 5min), removal of supernatant, and resuspension in 250uL Nuclear resuspension buffer (20mM pH 7.5 HEPES, 50% (v/v) glycerol, 75mM NaCl, 1mM DTT, 0.5mM EDTA, 1x protease inhibitor). An equal volume of NUN (high salt) buffer (50mM pH 7.5 HEPES, 1M urea, 300mM NaCl, 1mM DTT, 1x protease inhibitor) was added to nuclei pellets and gently mixed, following by 5min incubation on ice and periodic inversion. Chromatin pellets were isolated by centrifugation (1200xg, 5min). Non-chromatin-bound supernatant was removed. Chromatin pellets were washed with 500uL BufferA supplemented with 0.2% NP-40, followed by centrifugation, and removal of supernatant. The usually insoluble chromatin pellet was resuspended in 100uL Nuclear resuspension buffer, added to 1mL Trizol, and stored in 1.5mL centrifuge tubes at −20C for further processing.

Trizol samples were vigorously mixed with periodic heating at 50°C until the pellet dissolved. After adding 200uL chloroform, vigorously mixing, and centrifugation (16000xg, 15min), the aqueous phase was transferred to clean tubes or 96-well plates. An equal volume of ethanol was mixed, and samples were bound to Zymo spin I-96-XL plate (Catalog no. C2010) or individual Zymo Spin ii (Catalog no. C1008-50) columns by centrifugation (2000xg, 2min). Columns were washed twice with 500uL wash buffer (80% ethanol, 10mM pH 8.0 Tris). Samples were treated with DNAseI (2.5uL RQ1 Promega DNAseI, 1x DNAse buffer in 25uL total volume) while bound to columns and incubated at room temperature for 15min. Columns were washed with RNA binding buffer (2M guanidinium, 75% isopropanol), followed by two washes with 80uL wash buffer and a dry-spin. RNA was eluted by incubation with 25uL water for 5min followed by centrifugation. RNA yield was quantified by NanoDrop, typically yielding 5-20ug naRNA per sample.

#### RNA-seq library preparation

rRNA was depleted using Lexogen ribocop v2 kit according to the manufacturer’s protocol. RNA was eluted in 8uL water, all of which was used as input for NEB Ultra Directional II RNA-seq kits according to manufacturer’s protocol. Fragmentation time was adjusted from the recommended 15min to 5min to obtain larger insert sizes. Samples were pooled and sequenced on a single NovaSeq flow-cell (2×150bp paired end reads) by UChicago sequencing core.

### H3K36me3 CUT&Tag

Cells were grown as described above (naRNA-seq experimental methods) and 100,000 cells were frozen in 10% DMSO. CUT&Tag was performed as previously described^93^ (detailed protocol DOI: dx.doi.org/10.17504/protocols.io.z6hf9b6) using (Abcam catalog no. ab9050, lot GR3386101-2) polyclonal antibody, with the following modifications. Rather than 13 PCR cycles, we used 14 cycles. We determined this based on a test qPCR with ∼10% of the pre-PCR library, estimating that 14 PCR cycles with the remaining 90% would yield fluorescent signal at about halfway to the plateau, ensuring we have enough DNA material to quantify and sequence.

### Classification of unannotated splice junctions

We developed a method to predict the effect of splice junctions on transcript coding potential. Our method attempts to reconcile junctions identified from short-read RNA-seq with introns of annotated transcripts and predicts whether the junction is compatible with the open reading frame (ORF) of the annotated protein coding transcript (GENCODE v37). Specifically, we classify every annotated intron into one category in the following order of priority: protein_coding > processed_transcript > lncRNA > unprocessed_pseudogene > retained_intron > nonsense_mediated_decay, such that only introns that are uniquely used within transcripts labeled as nonsense_mediated_decay are classified to belong in that category. If an intron belongs to more than one category, it is classified to be in the category with the highest priority.

For splice junctions with coordinates do not match exactly that of an annotated intron, we separated junctions into four different categories for classification:

1. Junctions that only overlap with introns within the 5’ or 3’ UTRs are classified as UTR junctions and are not classified as NMD inducing (though in theory, new junctions in the 3’UTRs may trigger NMD). Additionally, junctions that only overlap with introns from transcripts classified as processed_transcripts, retained_intron or nonsense_mediated_decay are classified as the category with the highest priority. All remaining unannotated junctions that overlap with an intron that flanks an annotated coding exon are classified according to the methods described in (2), (3), or (4).
2. Junctions for which both 5’ and 3’ splice sites (ss) are annotated but are not used by a single intron in an annotated transcript. For these junctions, we translate the resulting mRNA using the frame from the upstream annotated exon CDS until the annotated end of the downstream annotated exon and classify the junction as NMD-inducing if an in-frame stop codon is observed.
3. Junctions for which only the 5’ss or 3’ss is annotated. For junctions with annotated 5’ss, we find all overlapping introns of annotated protein coding transcripts for which the 3’ss is within 60 nt of the junction 3’ss. If no such intron/transcript exists, then the junction is predicted to be NMD inducing. If one or more annotated intron exists, we translate the resulting mRNA up to the end of the annotated downstream exons (or up to an annotated stop codon). We classify the junction as NMD-inducing if all possible resulting mRNA harbor an in-frame stop codon. For junctions with annotated 3’ss, we similarly translate the resulting mRNA, setting the frame from the downstream annotated exon, and up to the start of the upstream exon. Again, we classify the junction as NMD-inducing if all possible resulting mRNA harbor an in-frame stop codon.
4. Junctions for which neither the 5’ss or 3’ss are annotated. Similar to (3), we first attempt to find introns that overlap with these junctions, allowing 60 nt to differ from the 5’ or 3’ ends or junction ends to be within annotated exons. Junctions that do not fit this criteria are classified as NMD-inducing. For the remainder of junctions, we translate the resulting mRNA and classify the junction as NMD-inducing if the resulting mRNA harbors an in-frame stop codon.

### Molecular QTL sharing

#### *π*_1_ sharing of eQTLs between RNA-seq datasets

Storey’s *π*_1_ statistic^94^, an estimate of the fraction of non-null hypothesis from a distribution of p-values, was used to estimate the fraction of features discovered one dataset (the “discovery dataset”) that have non-null effects in another dataset (the “ascertainment dataset”). For example, one may ask, “what fraction of eQTLs discovered in steady state RNA are eQTLs in naRNA?”. The P-value for each discovery eQTL (the nominal P-value for the top SNP:gene pair reported by QTLtools for each gene that passes a false discovery threshold for the genewise permutation test reported by QTLtools) was assessed in the ascertainment dataset to produce a distribution of P-values. *π*_1_ (the complement of *π*_0_), was estimated using the pi0est function from the qvalue package in R:

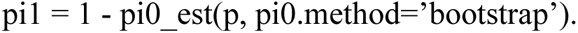

#### *π*_1_ sharing of eQTLs and hQTLs at TSS

For assessing the sharing between hQTLs and eQTLs, we assessed hQTLs at peaks within 500bp of annotated TSS (the 5’ most end of Gencode ‘basic’-tagged transcript structures) as the promoter for each gene. There is often more than one such TSS for each gene. Furthermore, there are multiple hQTL assays (i.e., H3K27ac ChIP-seq, H3K4me3 ChIP-seq) which we collectively consider as an ascertainment feature for each discovery feature. Therefore, there is not a one-to-one mapping of discovery features to ascertainment features for purposes of estimating *π*_1_. To answer the question, “what fraction of eQTLs have at least one hQTL?”, we modified the approach used to estimate *π*_1_ among eQTLs in RNA-seq datasets as follows to obtain a one-to-one mapping of discovery and ascertainment features: The minimum P-value among the n QTL test peaks that correspond to the promoter of a gene was considered a test statistic for obtaining a single ascertainment P-value for each gene. The distribution function of this test statistic under the null was estimated by repeatedly calculating min(x_1_, x_2_, …, x_n_), where x ∼ Uniform(0,1). An empirical cumulative distribution function of this test statistic was calculated after 10,000 repetitions of this process to obtain a single ascertainment P-value for each gene, and *π*_1_ was estimated as above.

#### Colocalization of molQTLs

We simultaneously assessed colocalization of molQTLs around each gene using hyprcoloc^95^. The following molQTL features were jointly considered for colocalization around each gene: Each H3K4me1, H3K4me3, & H3K27ac hQTL within 100kb of the gene, sQTLs introns fully contained in the gene body, H3K36me3 hQTLs (corresponding to the gene body), apaQTLs (features fully contained in the gene body), and eQTLs for the corresponding gene in each RNA-seq dataset (steady state RNA, 4sU 30m, 4sU 60m, naRNA). Summary statistics for a 100kb cis-window surrounding the gene were obtained for each molQTL were obtained using QTLtools nominal pass. Only molQTLs with a permutation pass P value < 0.1 were considered for colocalization. The hyprcoloc::hyprcoloc function was used in R with default settings to report clusters of colocalized molQTLs around each gene that satisfy a regional probability threshold (P_r_*=0.5 by default, corresponding to the probability threshold that all traits in the hyprcoloc iteration/cluster contain an association with a SNP) and regional alignment probability threshold (P_a_*=0.5 by default, corresponding to the probability threshold that all associations in the hyprcoloc iteration/cluster are aligned at a putative single causal SNP).

#### Effect size concordance of sQTLs and eQTLs

For each cluster with a significant sQTL intron, we classified the sQTL as a u-sQTL if it contains at least one sQTL intron (FDR<10%) in an unproductive intron. sQTLs with a nominally significant hQTL (P<0.01, for any H3K4me3, H3K27ac, or H3K36me3 trait) were filtered out. To avoid plotting non-independent sQTLs, we selected only a single sQTL intron per cluster, retaining the intron with the largest absolute value of sQTL beta. The top SNP of remaining sQTLs was used to look up the corresponding eQTL effect size in the host gene.

#### Colocalization of molQTLs with GWAS loci

Summary statistics for GWAS^96–102^ were downloaded from GWAS Catalog, or other source datasets. One mega-base windows centered at lead GWAS SNPs were determined as previously described^65^, using a lead SNP threshold of 5×10^-8^ to consider locus for colocalization. Summary statistics for all molQTL features within each GWAS locus window were obtained using QTLtools nominal pass, after selecting only those with a QTLtools permutation pass P-value < 0.1. Hyprcoloc was used with default settings to colocalize molQTLs and GWAS signals. Loci were categorized as “Only hQTL”, “Only eQTL”, or “Only sQTL”, if the only molQTLs to colocalize with the GWAS signal were either hQTLs (H3K36me3, H3K27ac, H3K4me3, or H3K4me1), eQTLs (in naRNA, 4sU, or steady-state RNA-seq), or sQTLs (in naRNA, 4sU-, or steady-state RNA-seq), respectively. If all of the molQTLs could be classified as eQTLs or hQTLs, or eQTLs and sQTLs, the loci was classified as “hQTL+eQTL” or “sQTL+eQTL”, respectively. All other loci with a molQTL colocalization were classified as “Other combinations”. We chose to classify these loci as “Other combinations” because sQTLs that also colocalize with both eQTL and hQTL, or with apaQTL, or other combinations, are hard to interpret and we wanted to refrain from suggesting these loci may be mediated by AS-NMD or alternative protein isoforms caused by AS.

### Enrichment of genomic annotations amongst QTLs

Genomic annotations include: ChromHMM annotations^103^ downloaded from UCSC genome browser based on ENCODE data from GM12878 LCL cell line, APA test peaks, splice donor regions (−3 to +7 from 5’ss), splice acceptor regions (−10 to 0 from 3’ss) and branchpoint regions (−40 to −10 3’ss) for annotated and unannotated splice sites, and miRNA binding sites (Downloaded from TargetScan v8.0^104^). The fine-mapping posterior inclusion probabilities (PIP) output by hyprcoloc were obtained for clusters containing an eQTL and a hQTL (transcriptional eQTLs), and clusters that contain an eQTL but no hQTL (post transcriptional eQTLs). Fold enrichment was calculated as the total fraction of PIP in a genomic annotation for transcriptional eQTLs, compared to post-transcriptional eQTLs. Confidence intervals were bootstrapped by 1000 resamples of the sets of transcriptional and post-transcriptional eQTLs.

### Risdiplam dosage series experiment

#### Cell growth, library preparation, and RNA-seq

LCLs growth, naRNA isolation, and conversion to sequencing libraries were performed as described above (naRNA-seq experimental methods). Risdiplam was added 24 hours prior to cell collection. Total (“steady-state”) RNA was converted to sequencing libraries using NEB polyA capture kit (product no) followed by NEB Ultra directional II RNA-seq library kits. Steady-state RNA-seq libraries were sequenced by (2×150bp paired end). naRNA-seq libraries were sequenced by UChicago sequencing core (2×150bp paired end).

#### Identification and quantification of risdiplam-induced exons

All splice junctions containing GA|GU in reference genome sequence at the 5’splice site were assessed for a significant positive dose:response correlation using leafcutter’s intron excision ratio as the response. Significance was assessed using R’s ′cor.test(…, method=’spearman’)′ function, and P-values were adjusted for multiple testing with Storey’s qvalue method. All significant (q<0.1) GA|GU introns with a leafcutter-clustered splice junction that has a 3’ss 500bp upstream of the GA|GU splice donor were considered as risdiplam-induced cassette exons. Splicing at these cassette exons was re-quantified using the cassette exon PSI metric (see above).

#### Prediction of transcripts effects

We utilized Gencode transcript structures and their predictions for coding potential (“basic”-tagged transcripts being deemed as productive) to annotate cassette exons that use an annotated GA|GU downstream splice junction. Cassette exons with an unannotated downstream GA|GU splice junction were translated in-frame using the most-expressed “basic”-tagged transcript (transcript quantifications derived from Salmon) as a reference, using custom scripts. For simplicity, we classified unannotated exons as unproductive if the exon-included translation is shorter than the exon-excluded translation.

#### Quantification of expression and splicing

Gene expression was quantified using featureCounts as described above. naRNA-seq samples were analyzed using edgeR^105^, with two contrasts: (1) 100nM risdiplam vs DMSO and (2) 3160nM risdiplam vs DMSO contrast. The unconventional experimental design of the titration series experiment (a single replicate at eight doses) precluded use of many standard differential splicing or differential gene-expression analysis approaches. For those samples, we fit splicing quantifications (in units of cassette exon PSI) and gene-expression quantifications (in units of log2CPM) to a 4-parameter log-logistic curve using the drc package^106^ in R: ′drc::drm(formula = cpm ∼ dose, fact=LL.4(), …)′. Effect size estimates and standard errors at 100nM and 3160nM were extracted from the model fits using ′predict(…, se.fit=T)′. FDR was estimated with Storey’s qvalue. Genes were classified as transcriptionally regulated if they had an absolute log_2_FC between naRNA and steady-state RNA less than 1.5, with FDR<0.1 in both naRNA and steady-state RNA. If genes had FDR<0.1 in steady-state RNA but with an effect size difference between naRNA and steady-state greater than log_2_(1.5), they were classified as post-transcriptionally down-regulated. Similarly, genes had FDR<0.1 in steady-state RNA but with an effect size difference between naRNA and steady-state less than log_2_(1.5), they were classified as post-transcriptionally up-regulated.

#### Gene set enrichment

Gene sets were defined as follows: “Disease genes” are defined as genes from the OMIM genemap database with a string match to ‘dominant’ in the phenotype column. Kinases (GO:0016301), transcription factors (GO:0003700), nuclear hormone receptors (GO:0016301), G-protein coupled receptors [GO:0004930 that are not in the set of olfactory receptors (GO:0004984)], and ion channels (GO:0015075) were downloaded from Msigdb^107^. Gene sets were filtered for tested expressed genes considered in differential splicing and differential expression testing. Significant enrichment of differentially expressed or spliced genes was assessed using a hypergeometric test.

## Supplemental Figures

**Fig S1.**
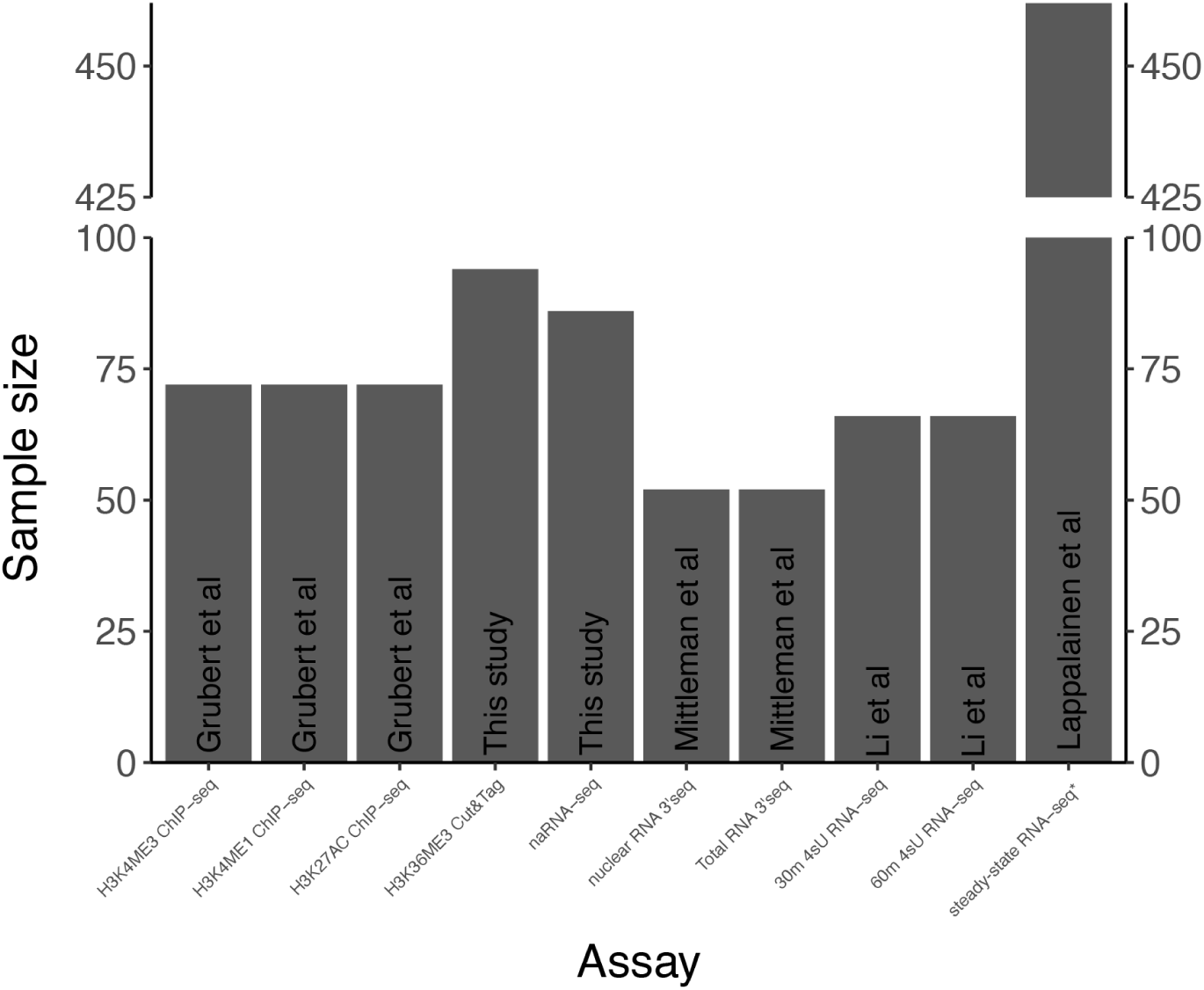
Data sources and sample size. Source publication, assay, and sample size used for QTL analyses^39–42^. All samples are lymphoblastoid cell lines of Yoruba ancestry, except *poly RNA-seq dataset contains 89 Yoruba ancestry cell lines, and 362 non Yoruba ancestry cell lines. Some analyses (STAR Methods) use only Yoruba ancestry cell lines of this dataset.

**Fig S2.**
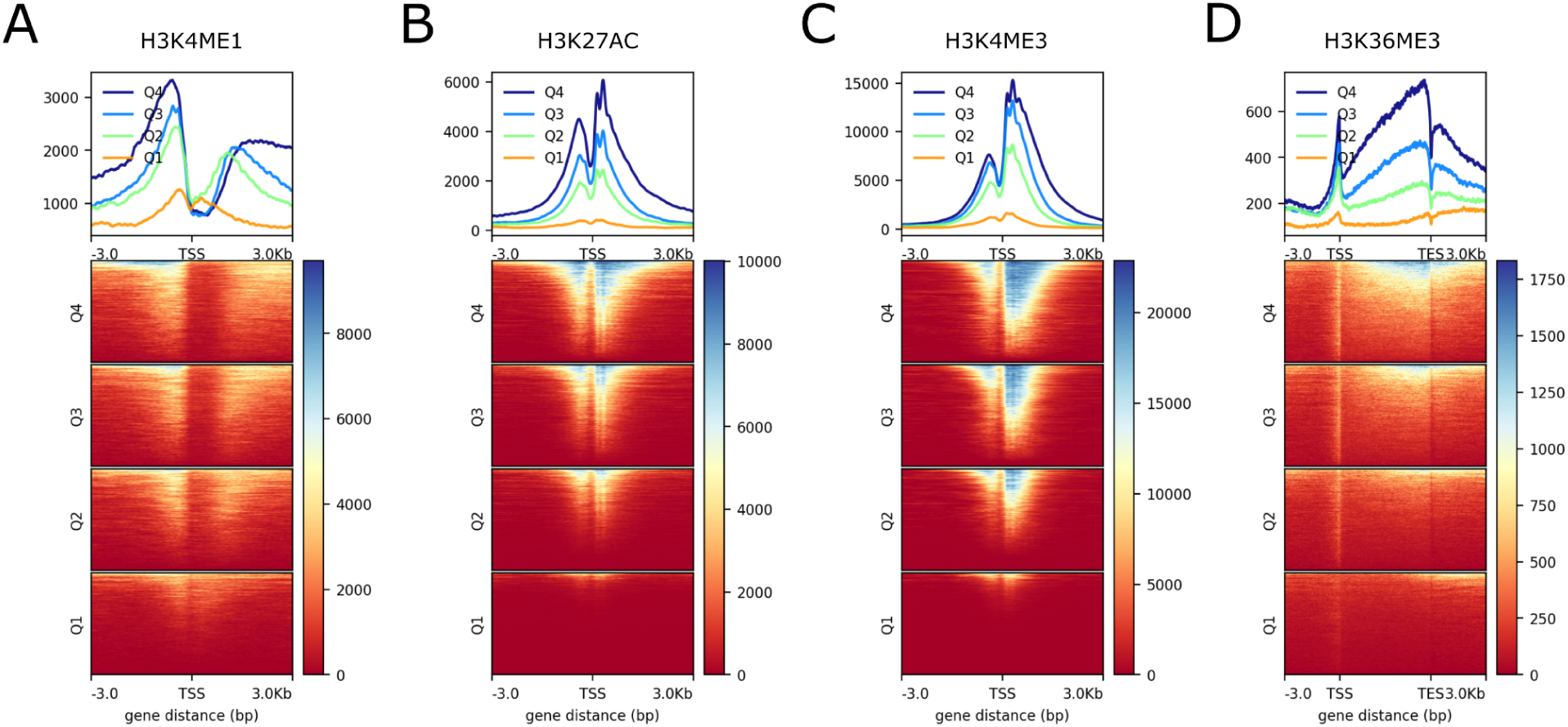
Chromatin profiling metagene plots. (A) H3K4me1 (enhancer mark) ChIP-seq signal. Genes grouped by expression quartile, as determined by polyA-RNA-seq RPKM values of the top 14000 expressed genes. (B) H3K27ac (promoter/enhancer mark) ChIP-seq signal. (C) H3K4me3 (promoter mark) ChIP-seq signal. (D) H3K36me3 CUT&Tag signal. Enrichment at the promoter region may represent some cross-reactivity with other marks.

**Fig S3.**
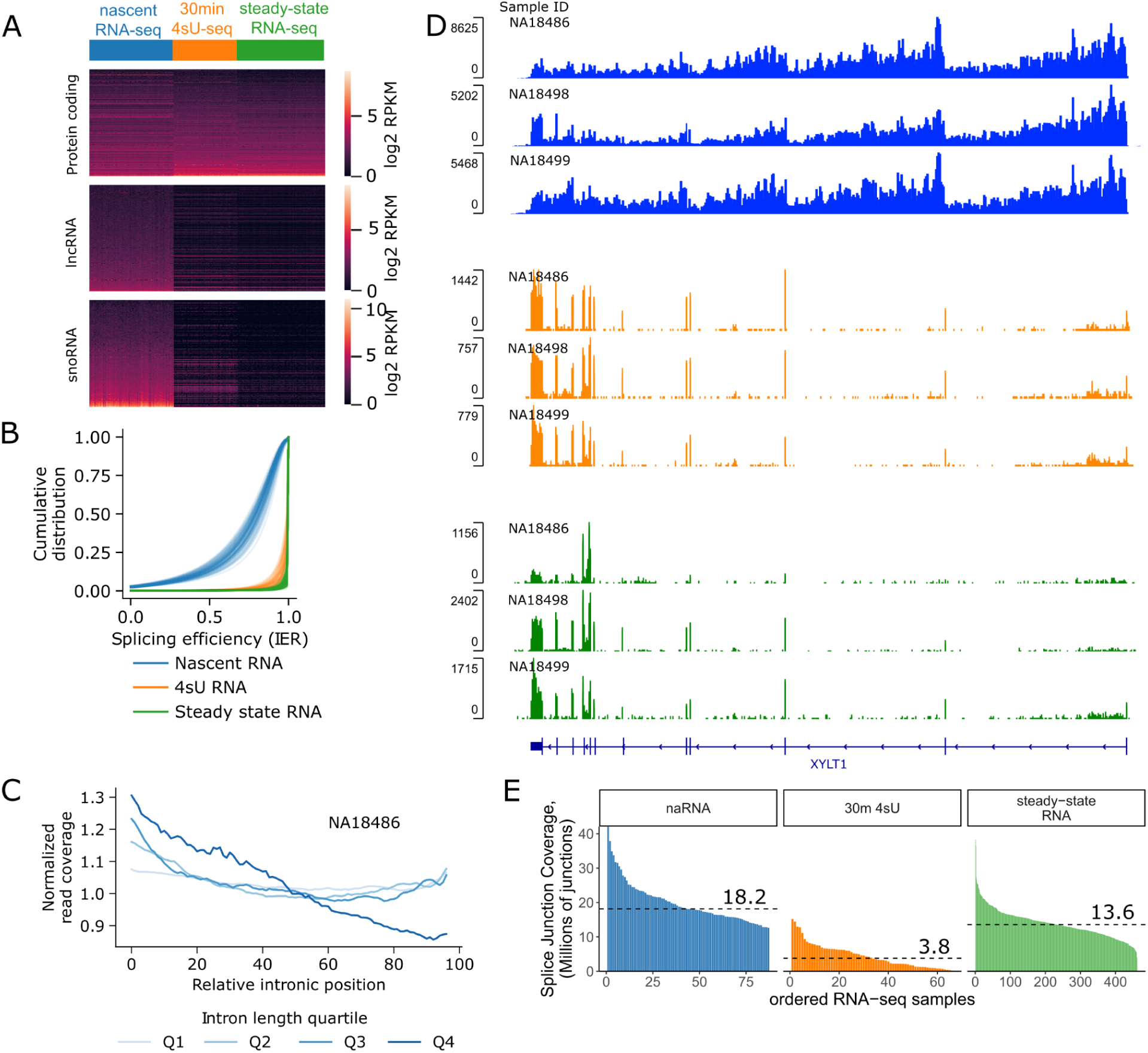
Overview of nascent RNA-seq. (A) Nascent RNA-seq (naRNA-seq) captures nuclear-retained, non-polyadenylated, and rapidly decayed RNAs (snoRNAs, lncRNAs), that are absent from labeled and steady-state RNA-seq datasets. (B) naRNA transcripts are only partially spliced. Example sawtooth pattern in nascent RNA in the gene *XYLT1*. The nascent nature of transcripts in naRNA creates a 5’ bias in coverage, and in combination with co-transcriptional splicing, creates a sawtooth pattern of coverage. (C) Meta-intron coverage plot in LCL naRNA-seq sample NA18486 confirms the 5’ bias in intronic coverage genome-wide, consistent with the nascent nature of naRNA transcripts. Longer introns are naturally expected to have steeper slopes than short introns when intron lengths are rescaled for metaplot. (D) The splicing efficiency metric is based on the ratio of spliced and unspliced (intron:exon junction) reads, and varies between 0 and 1, with 1 indicating all reads are spliced. The cumulative distribution of splicing efficiency across all introns in expressed genes, for each RNA-seq sample from naRNA, recently transcribed RNA (30 min 4sU), and steady-state RNA. (E) Number of exon-exon splice junction reads in RNA-seq samples. The median in each dataset is marked with a labeled dashed line.

**Fig S4.**
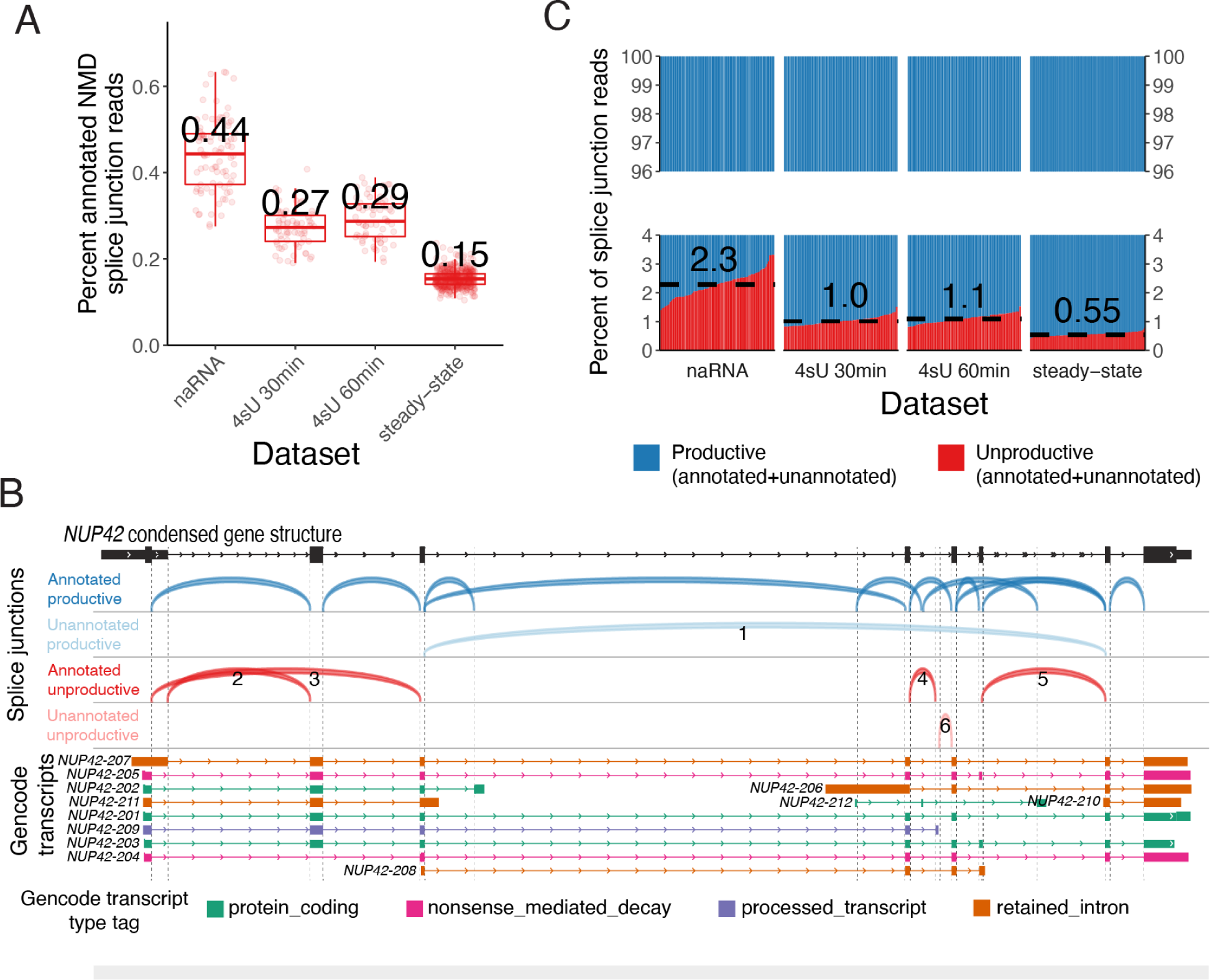
Classification and quantification of splice junction classes across datasets. (A) The percent of splice junctions in each sample (point) that are uniquely attributable to transcripts tagged as “nonsense_mediated_decay” (Gencode v37). Box and whiskers show quartiles for samples in each RNA-seq data-type. Median for each data-type is labeled. (B) Splice junctions (arcs) overlapping the *NUP42* gene illustrate our methods for classifying splice junctions. Annotated splice donors and splice acceptors are marked with vertical dashed lines in dark and light gray, respectively. Annotated productive junctions are defined by their presence in at least one transcript with the value of “protein_coding” in the transcript type tag. Unannotated productive junctions are not in any Gencode transcripts, and skip exons in the principal isoform such that the reading frame is maintained (i.e., splice junction marked with 1). Annotated unproductive junctions are unique to Gencode transcripts not tagged with “protein coding”. Splice junction 2 is unique to *NUP42-207*, a “retained_intron” tagged transcript. This splice junction uses a deep intronic 5’ss, creating a premature termination codon. Junctions 3 and 5 are unique to transcripts tagged as “nonsense_mediated_decay”, and junction 4 is unique to a transcript tagged with “processed_transcript”. All other junctions are classified as Unannotated unproductive. We attempted to translate the resulting transcripts that use these junctions, finding that they overwhelmingly introduce frameshift or in-frame stop codons (STAR Methods), such as the splice junction 6 which we predict to introduce a frameshift. (C) Similar to (B), where sample is represented as a column, and the fraction of splice junction reads that are either productive (annotated or unannotated, classified as in (B), blue) or unproductive (annotated or unannotated, classified as in (B), red). The median in each dataset is marked with a dashed line and labeled.

**Fig S5.**
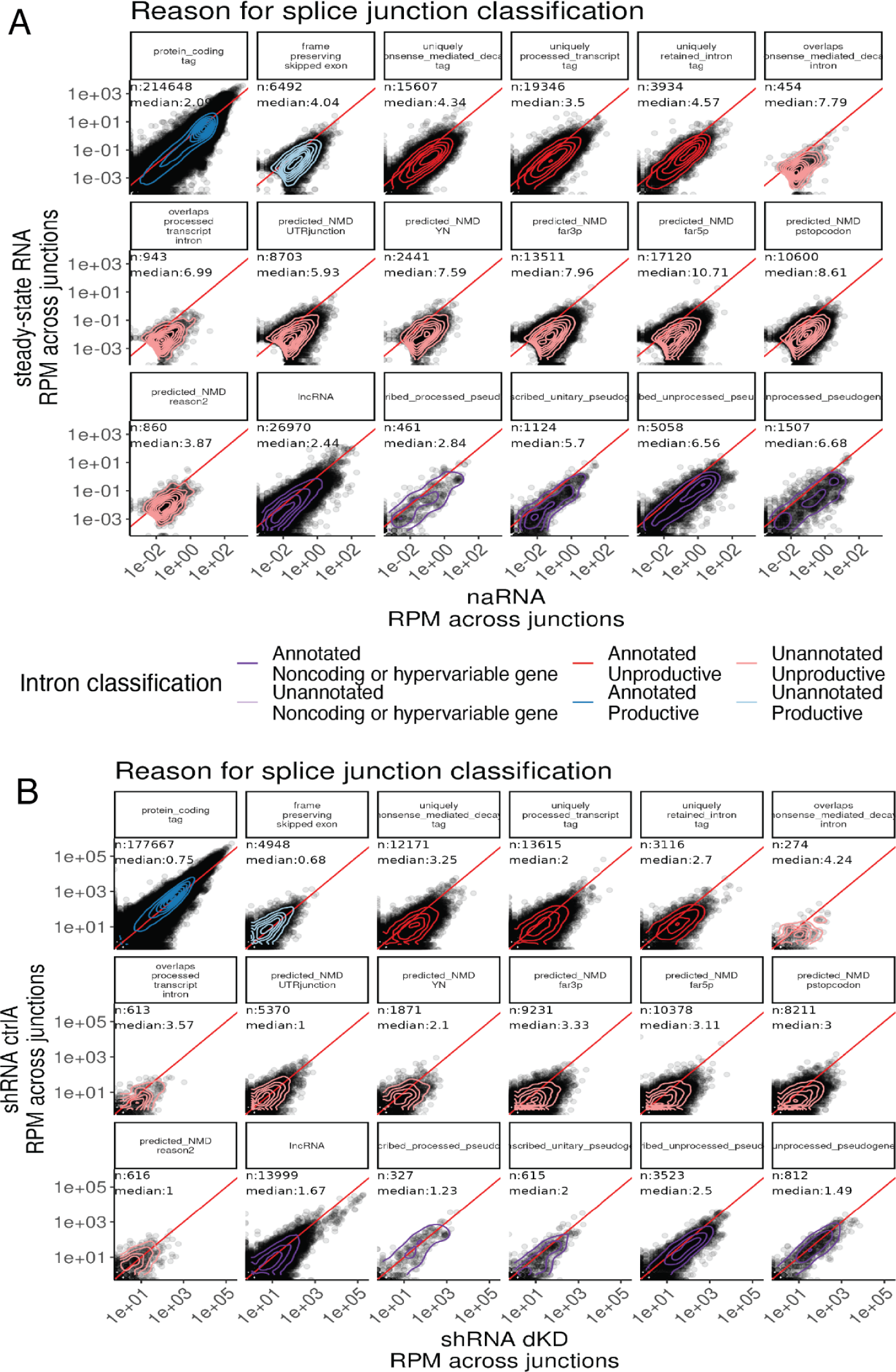
Splice junction abundances across datasets are consistent with classifications of splice junctions as productive or unproductive. (A) Splice junctions (introns) are classified into 6 types based on whether they are annotated, predicted to create a productive transcript, and by whether they are in a protein-coding gene (colors, STAR Methods). For each subcategory of these six classes, a scatter plot depicts relative junction prevalence in naRNA vs steady-state RNA. Each point is a unique splice junction. The median fold changes across all n junctions in each category labeled in each plot. Splice junctions annotated or predicted as productive are generally similarly present in naRNA versus polyA RNA. Splice junctions annotated as unproductive are generally depleted from polyA RNA. (B) Similar to (A), but showing relative change in scramble shRNA control compared to shRNA double knockdown of NMD factors *SMG6* and *SMG7*^28^.

**Fig S6.**
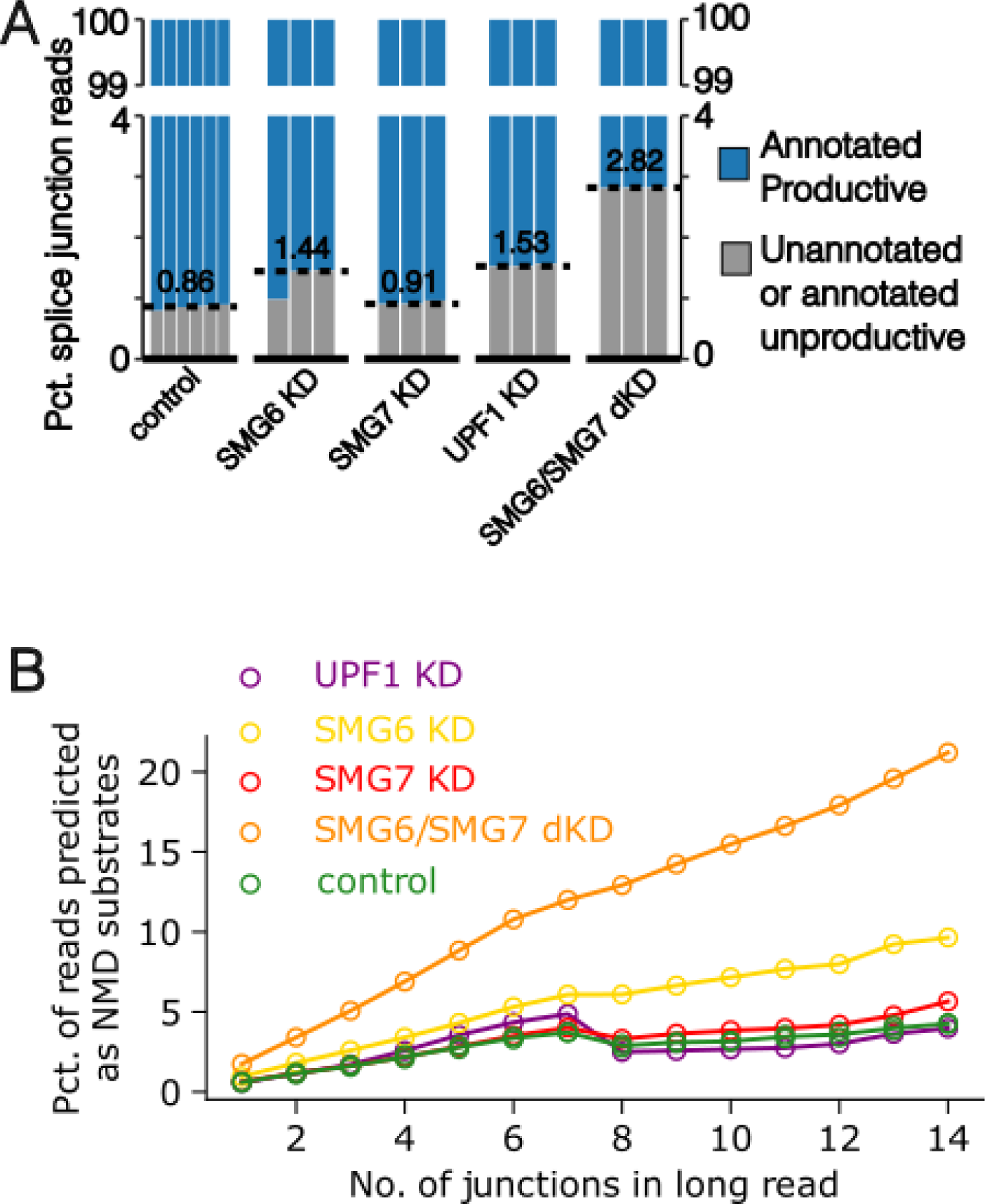
Percentage of unproductively spliced reads upon knockdown of NMD factors. (A) Fraction of splice junction reads in each short read steady-state RNA-seq sample that are in Gencode-annotated productive transcript structures (blue), versus unannotated or annotated unproductive transcript structures (gray). Biological replicates represented by each column, with dashed lines to indicate median for each group. NMD factors were knocked-down (KD) singly or as double knockdown (dKD) with shRNA in HeLa cells with an shRNA scramble control^28^. (B) Nanopore long-read sequencing quantifies the percent of full-length reads that are targeted by NMD, defined as containing at least one unproductive junction, as a function of the number of splice junctions in the read. Knockdown experiments of similar design as in (A)^29^.

**Fig S7.**
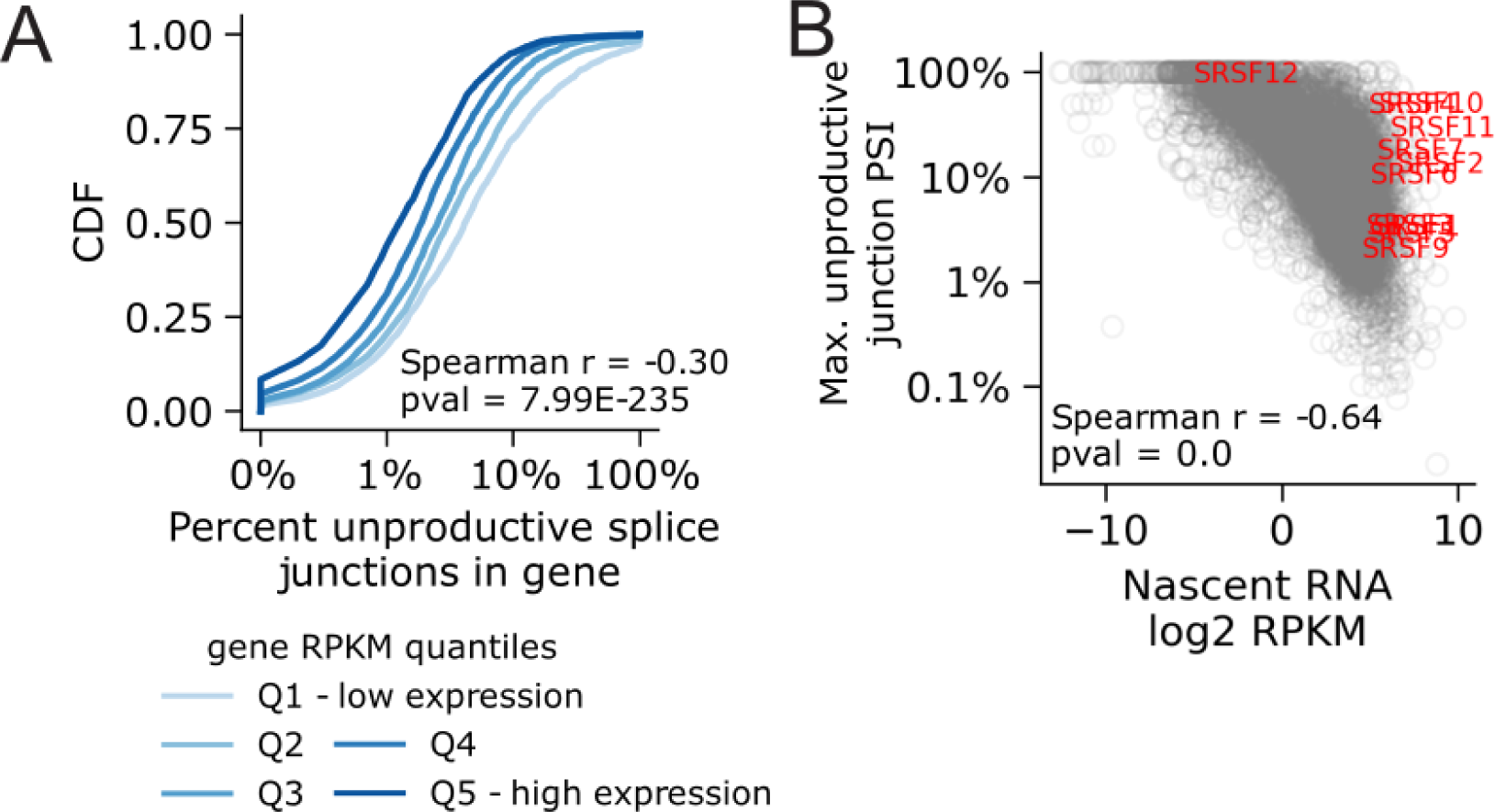
Correlation of unproductive splicing and gene expression in naRNA. (A) Highly expressed genes have a lower unproductive splicing rate, as measured by the genewise percent of splice junction reads that are unproductive. Correlation summarized with spearman correlation coefficient and P value, and visually presented as cumulative distribution of percent unproductive splice junctions, grouped by expression quintiles. (B) Correlation between gene expression and the maximum junction PSI of any unproductive junction in the gene, a proxy for percent of unproductive transcripts. The PSI of a junction is the number of reads mapping to the junction, divided by the maximum number of reads mapped to any junction in the same gene. The junction with the highest number of reads in a given gene has a junction PSI of 100%. SRSF genes (red) are well-known examples^13^ of genes with high gene expression and high unproductive junction PSI.

**Fig S8.**
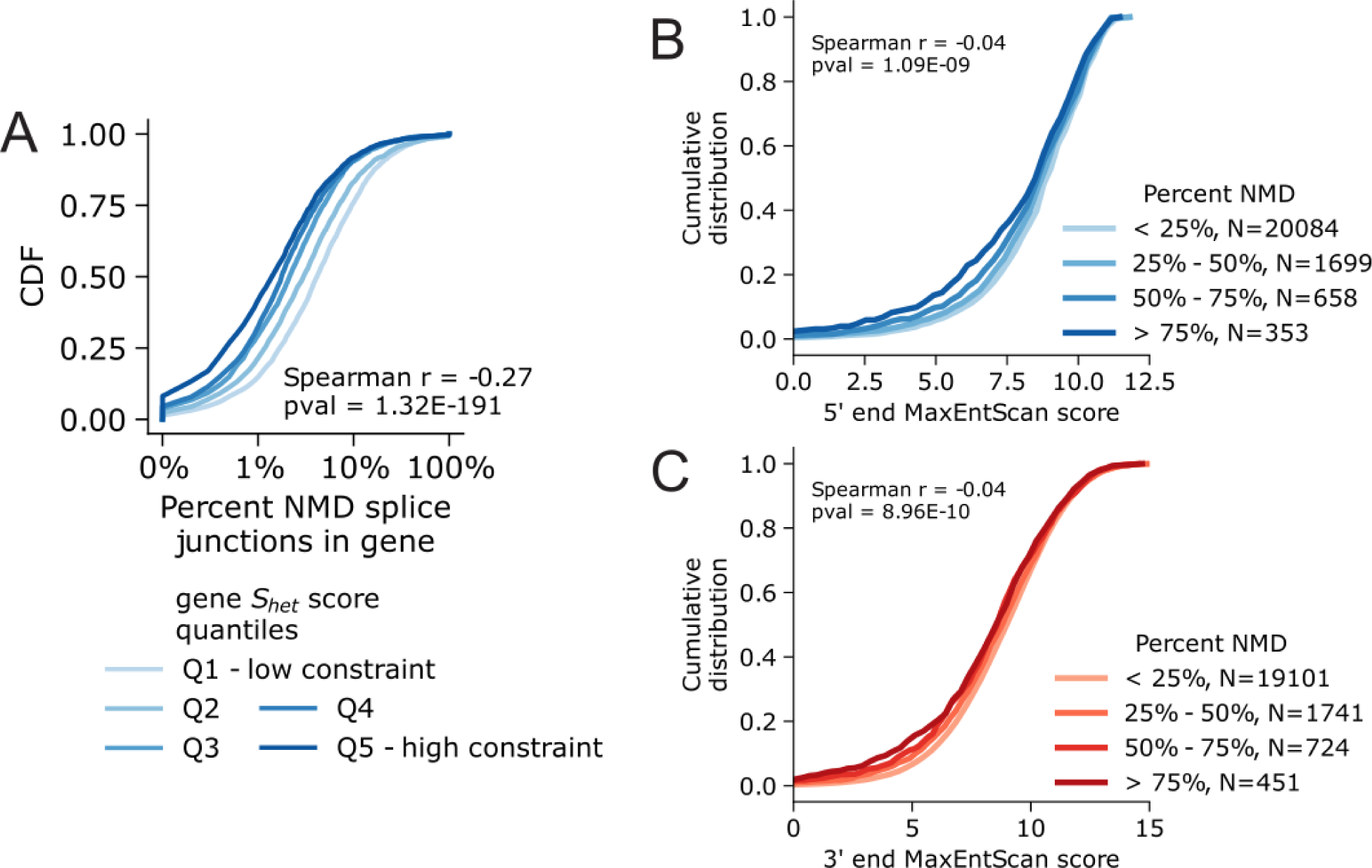
Gene constraint and splice site strength correlate with percent of unproductive splice junctions. (A) Rate of unproductive splicing negatively correlates with S_het_, a measure of haploinsufficiency. Correlation summarized with spearman correlation coefficient and P value, and visually presented as cumulative distribution of percent unproductive (NMD-inducing) splice junctions, grouped by S_het_, quintiles. (B) Rate of unproductive splicing overlapping annotated productive introns is negatively correlated with strength of the productive intron’s 5’ splice site (MaxEntScan). (C) Same as (B), but for 3’ splice sites.

**Fig S9.**
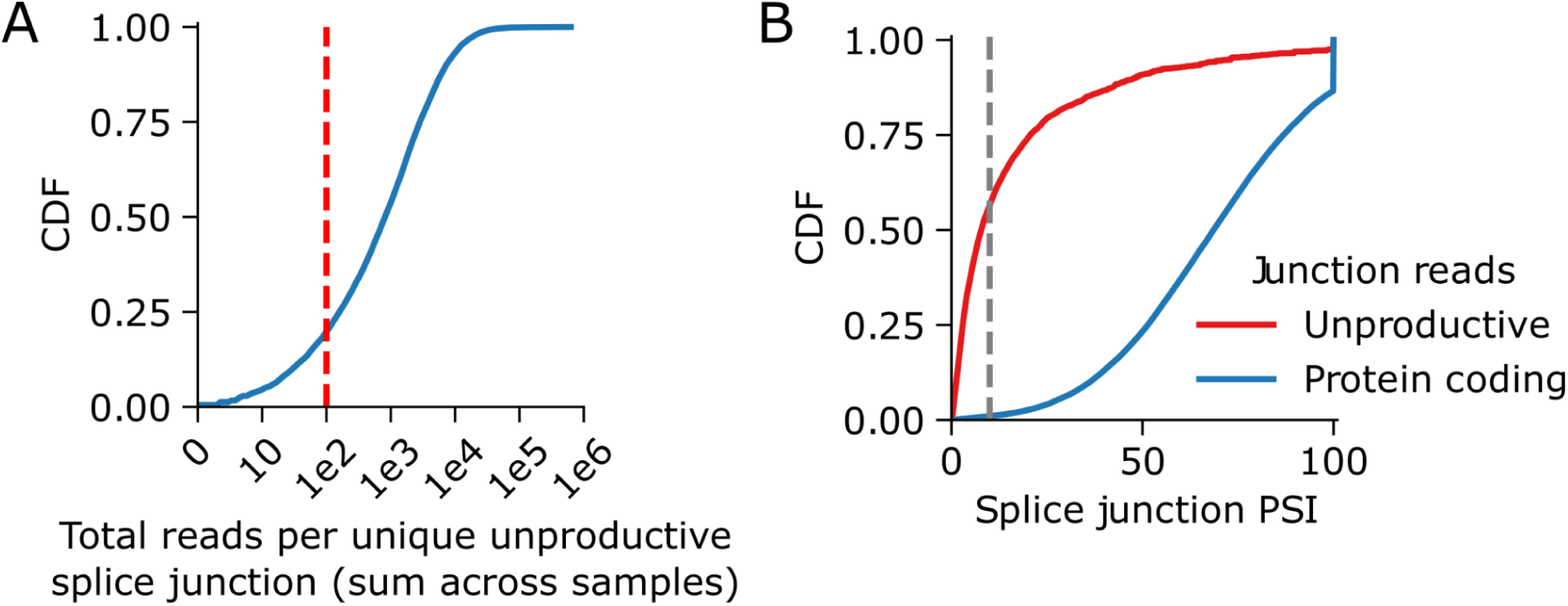
Distribution of unproductive splice junction usage. (A) Cumulative distribution of the total number of reads mapping to each unique unproductive splice junction across all naRNA-seq samples. We filtered out the ∼20% of splice junctions with fewer than 100 reads (red) for the unproductive junction contribution analysis in (B): Distribution of the splice junction PSI of unproductive and productive splice junctions, averaged across all naRNA-seq samples. The PSI of a junction is the number of reads mapping to the junction, divided by the maximum number of reads mapped to any junction in the same gene. The junction with the highest number of reads in a given gene has a junction PSI of 100%. More than 50% of unproductive junctions have a junction PSI of 10% or less (gray line).

**Fig S10.**
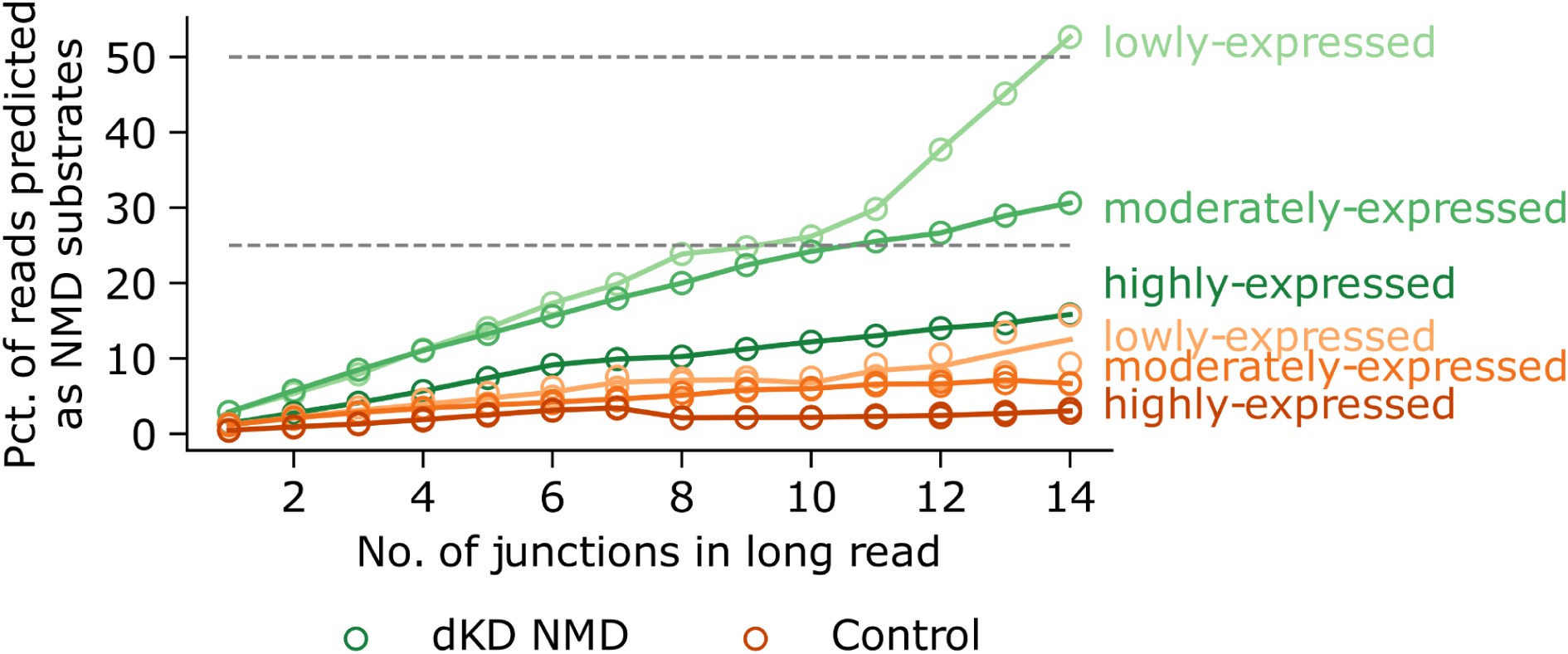
Over 50% of transcripts from lowly-expressed long genes are unproductive. Percent of full-length Nanopore reads that are targeted by NMD, as a function of the number of splice junctions in the read, and stratified by gene expression. dKD NMD corresponds to the shRNA SMG6/SGM7 double knockdown data in HeLa cells^29^, and control corresponds to the shRNA scrambled controls from the same dataset. Expression rankings are based on the RPKM expression of shRNA *SMG6*/*SGM7* double knockdown in HeLa cells using short-read data^28^. Lowly-expressed corresponds to genes in the bottom quartile (0.01-1.3 RPKM), moderately-expressed are genes in the second and third quartiles (1.3-15.6 RPKM), and highly-expressed are genes in the top quartile of expression (>15.6 RPKM). The gray lines mark the 25% and 50% thresholds of mRNA molecules targeted for NMD.

**Fig S11.**
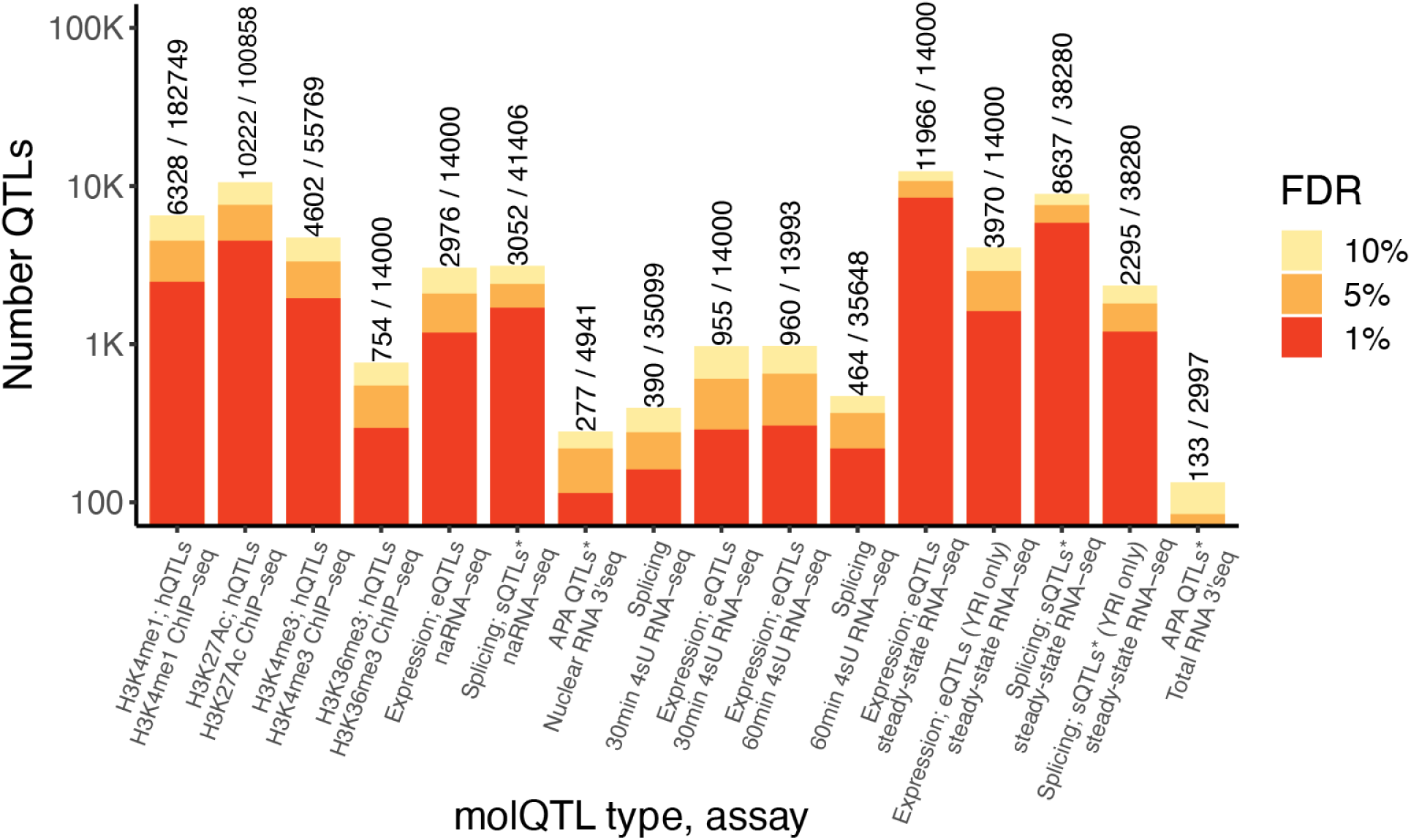
Number of QTLs. The number of QTLs at various false discovery rate thresholds. Numbers indicate the number of QTLs at 10% FDR, and the total number of test features. Numbers of *sQTLs and apaQTLs are counted once per local QTL. That is, a sQTL that affects multiple introns in the same LeafCutter cluster is only counted once.

**Fig S12.**
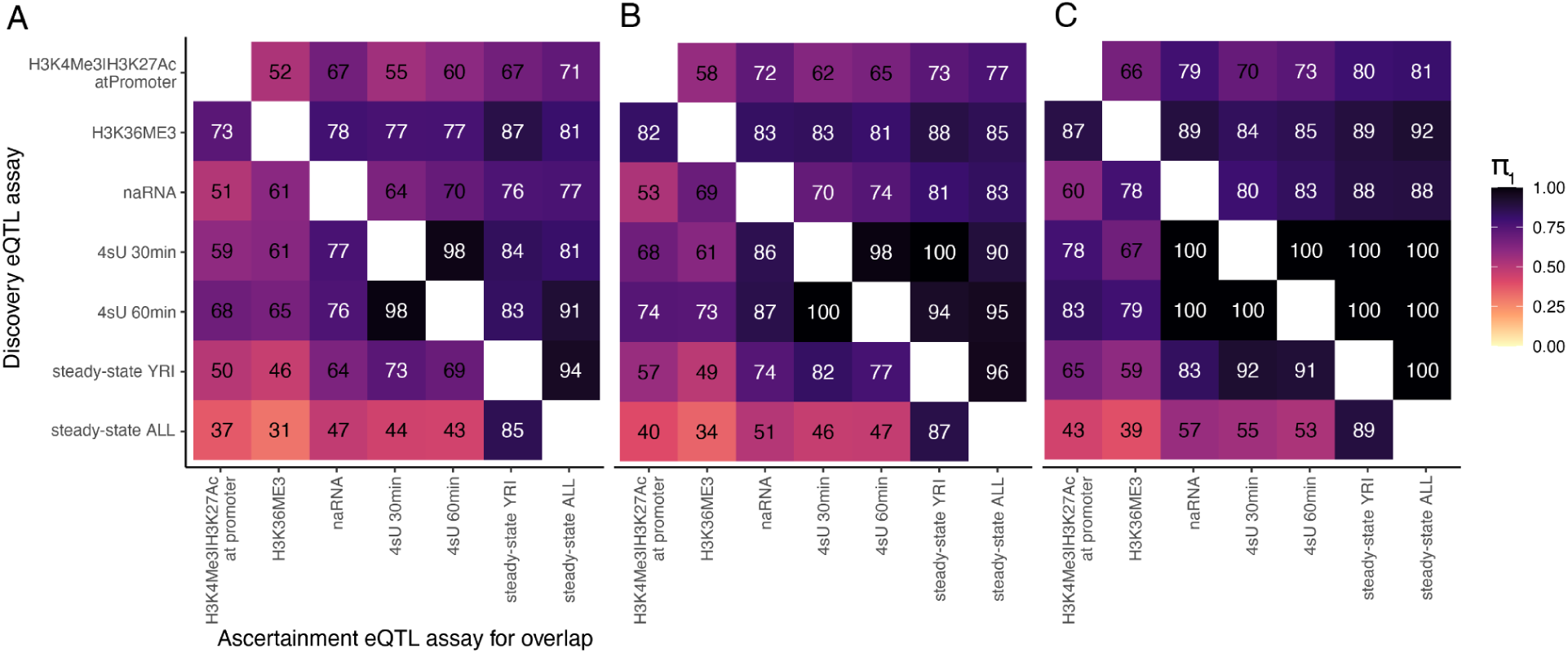
π_1_ sharing across phenotypes that measure gene expression. (A) QTLs discovered at 10% FDR (rows) were assessed for overlap by measuring the π_1_ statistic for the corresponding SNP:gene pair in different assays (columns). Promoter marks include H3K27ac, and H3K4me3, and consider any annotated promoter for the corresponding gene (STAR Methods for details). (B, C) same as A, using 5% and 1% FDR threshold for discovery QTLs, respectively. Overall, the effect of promoter hQTLs are apparent throughout subsequent datasets (upper right of square). By contrast, effects discovered in steady-state RNA are less likely to be apparent as promoter hQTL signals (lower left of square), suggesting the existence of post-transcriptional regulation in determining steady-state eQTL signals. Steady-state eQTLs were mapped using either n=89 (similar sample size as other assays) ancestry-matched samples (YRI), or using n=453 mixed ancestry samples.

**Fig S13.**
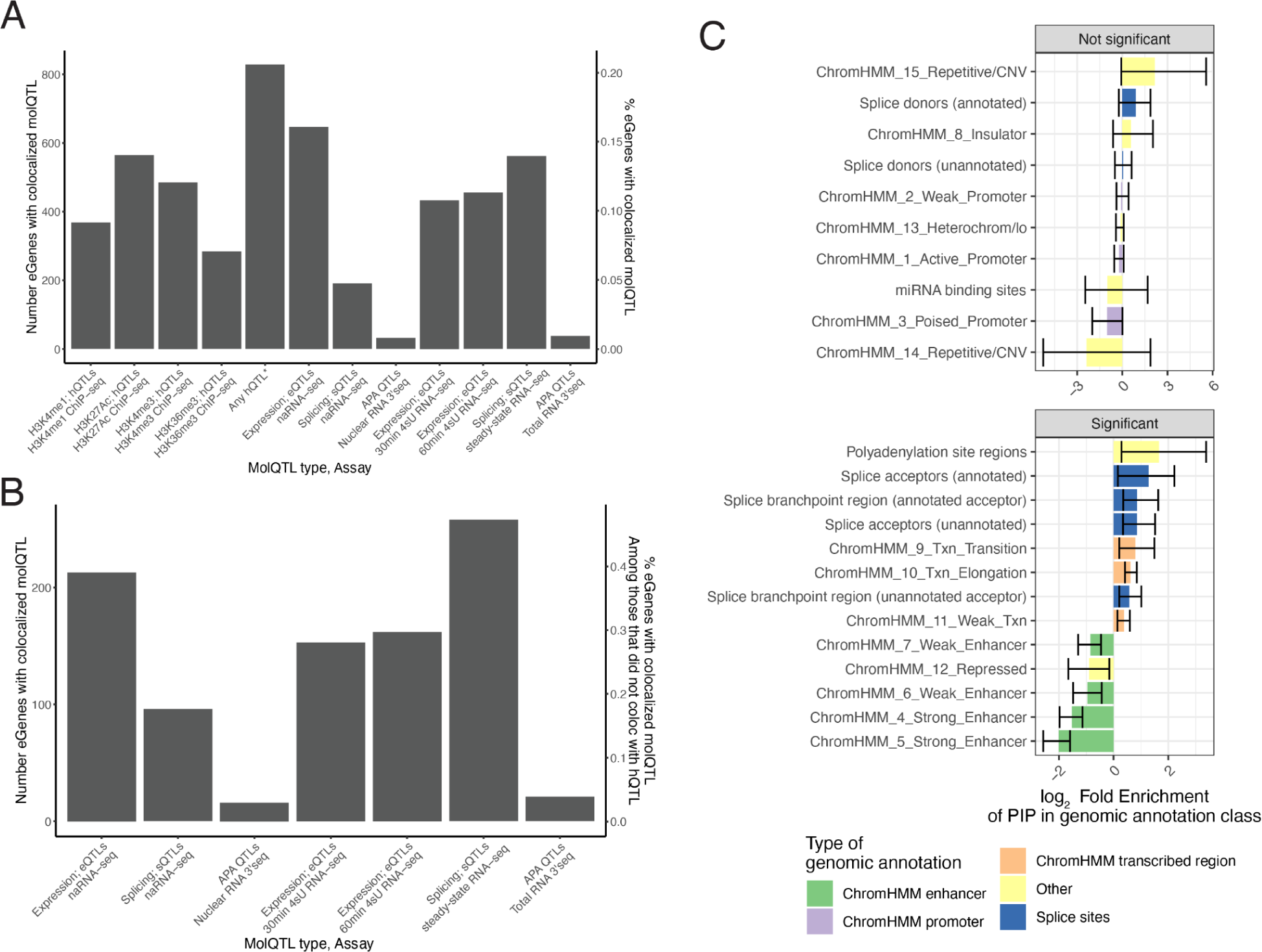
Multitrait colocalization used to identify post-transcriptionally functioning eQTLs. (A) The number (and fraction) of eGenes (steady-state eQTLs, mapped in n=89 Yoruban-ancestry samples) with at least one colocalizing molQTL for each type of molQTL. (C) The number (and fraction) of eGenes with at least one colocalizing molQTL, among the post-transcriptionally-regulated eGenes, defined as those colocalize with a molQTL but not with any hQTL (H3K4me1, H3K4me3, H3K27Ac, H3K36me3 QTLs). (C) enrichment of eQTL signal in genomic annotations, comparing enrichment in transcriptionally regulated eQTLs versus post-transcriptionally regulated eQTLs. Enrichment is defined as the relative fraction of fine-mapping posterior inclusion probability (PIP), in the given genomic annotation in post-transcriptionally regulated eQTLs compared to transcriptionally-regulated eQTLs. Boottrapped 95% confidence intervals used to determine the enrichments that are significant (bottom facet). Splice site related annotations make up many of the significantly enriched annotation classes. STAR Methods for details.

**Fig S14.**
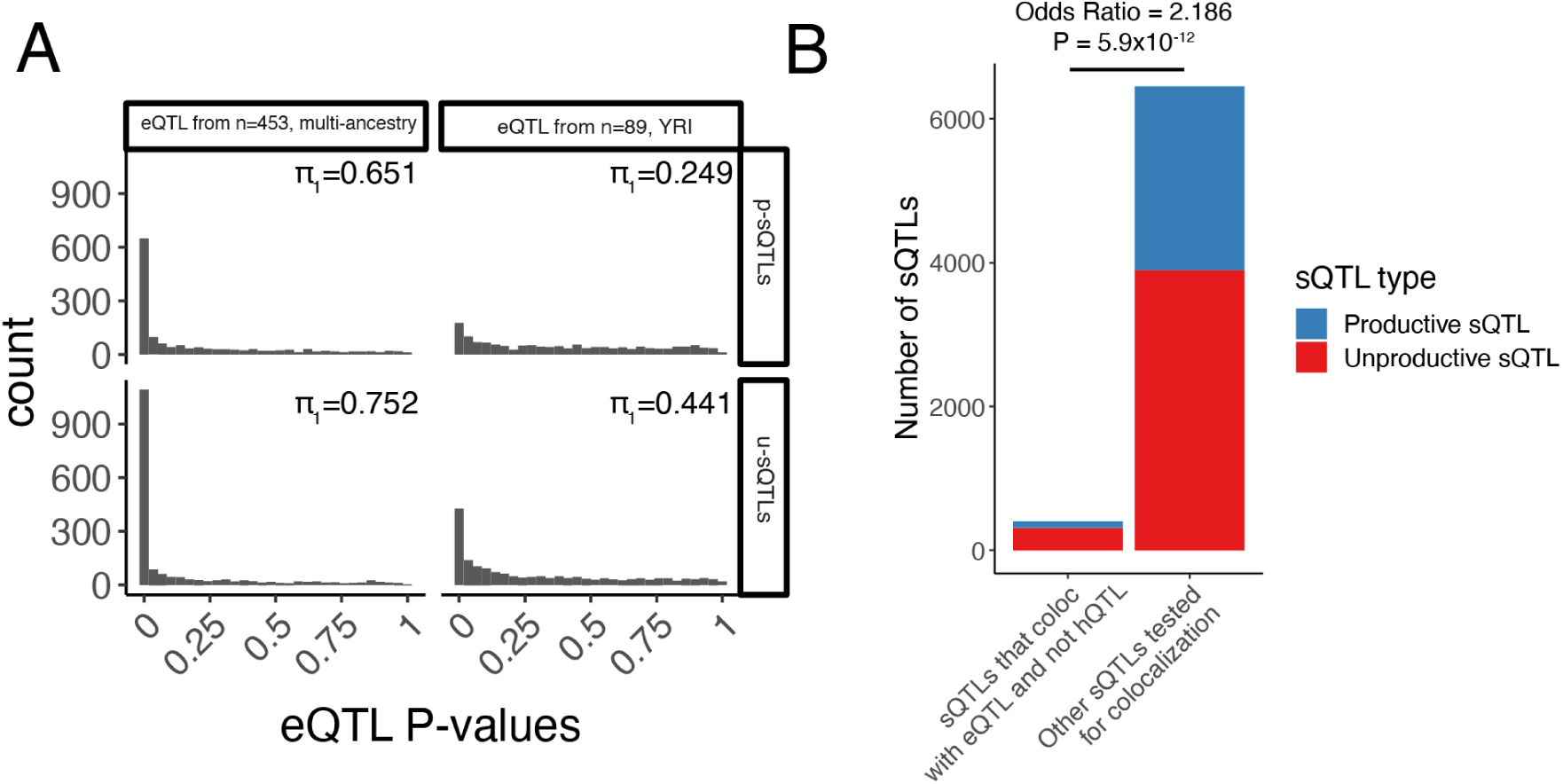
Enrichment of eQTL signals among sQTLs that affect unproductive splice junctions. (A) u-sQTLs (sQTLs that alter the balance of unproductive and productive isoforms, defined as sQTLs in LeafCutter clusters that contain an sQTL in an unproductive splice junction and are not nominally any hQTL) are enriched for eQTL signal, as evidenced by inflation for small P-values, compared to p-sQTLs (which contain significant sQTLs only in productive splice junctions). π_1_ statistic labeled in top right of P-value histograms. Importantly, unlike colocalization analysis in (B), the general inflation of small eQTL P-values implicitly accounts for sQTLs that may explain non-primary eQTLs. The effect is observed when estimating eQTL signal from all n=452 samples, as well as in the more power-limiting case when n=89 ancestry-matched samples. (B) sQTLs that colocalize with an eQTL are enriched in u-sQTLs compared to p-sQTLs. If a sQTL was tested for eQTL colocalization in multiple genes (which both overlap the splice junction), it is tallied multiple times. P value from hypergeometric test.

**Fig S15.**
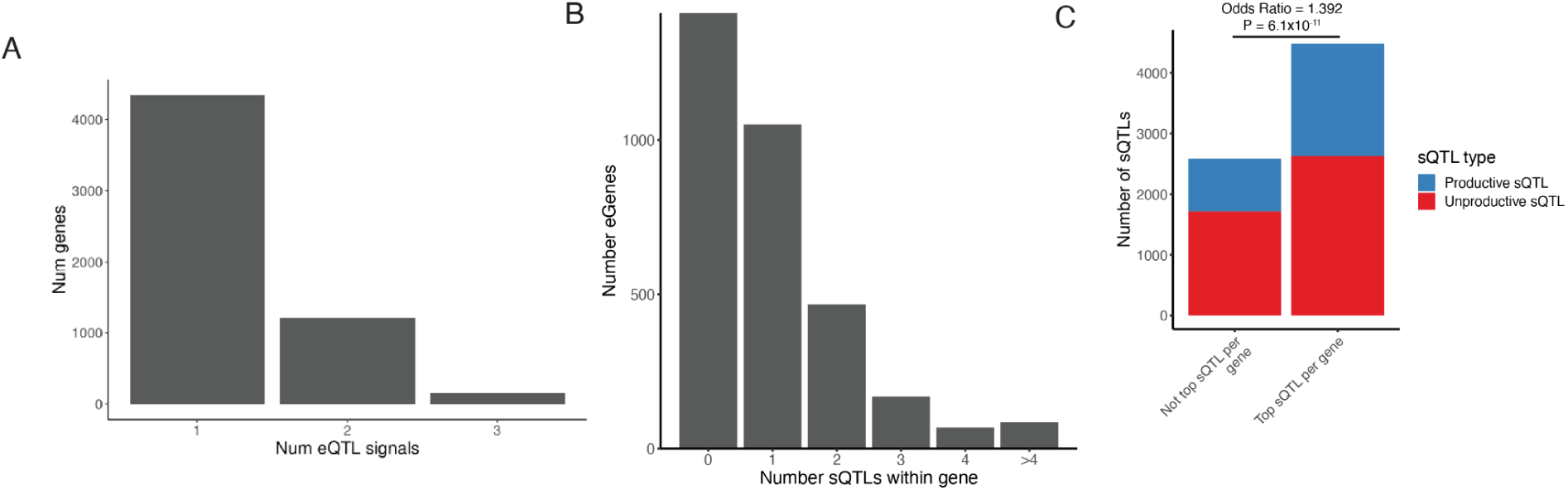
Importance of considering secondary eQTLs, hQTLs, and sQTLs when assessing overlap amongst molQTLs. (A) Number of eGenes with more one, two, or three independent effects identified by SuSiE. While more independent eQTL signals may biologically exist for a single gene, we only searched for a maximum of three signals per gene due to power limitations and linkage disequilibrium. (B) Number of sQTLs (distinct sQTL clusters) discovered amongst the n eGenes. (C) Primary sQTLs (most significant sQTL cluster per gene) are enriched for p-sQTLs over u-sQTLs, compared to non-primary sQTLs. P value from hypergeometric test.

**Fig S16.**
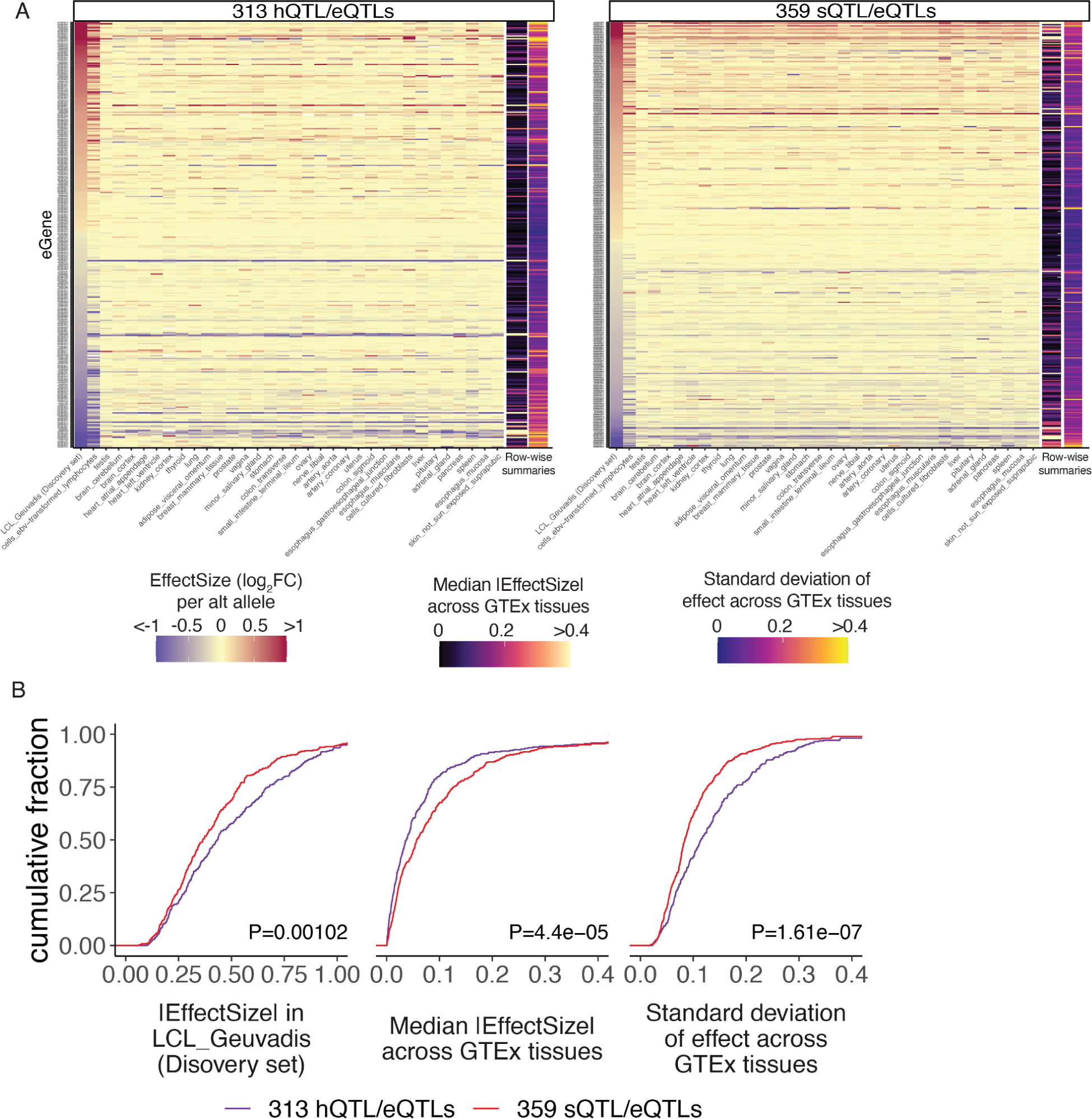
Transcription-mediated eQTLs (hQTL/eQTLs) are more tissue-specific than splicing-mediated eQTLs (sQTL/eQTLs). (A) A set of transcription-mediated eQTLs (hQTL/eQTL colocalization) identified in our source dataset was compared to a set of splicing-mediated eQTLs (sQTL/eQTL colocalization without nominal hQTL signal), by estimating the SNP:gene effect across 38 GTEx tissues (columns) for each eGene (rows). Row-wise summary statistics were calculated and plotted as extra columns. (B) Row-wise summary statistics are plotted as cumulative distributions for visual contrast. From left to right, we see that (1) the absolute effect size in the Geuvadis LCL discovery dataset is slightly greater for hQTL/eQTLs than sQTL/eQTLs. Despite this, the sQTL/eQTLs have a (2) larger median effect size across GTEx tissues, and (3) have a smaller standard deviation of effect size across tissues. P values from two-sided Mann-Whitney test.

**Fig S17.**
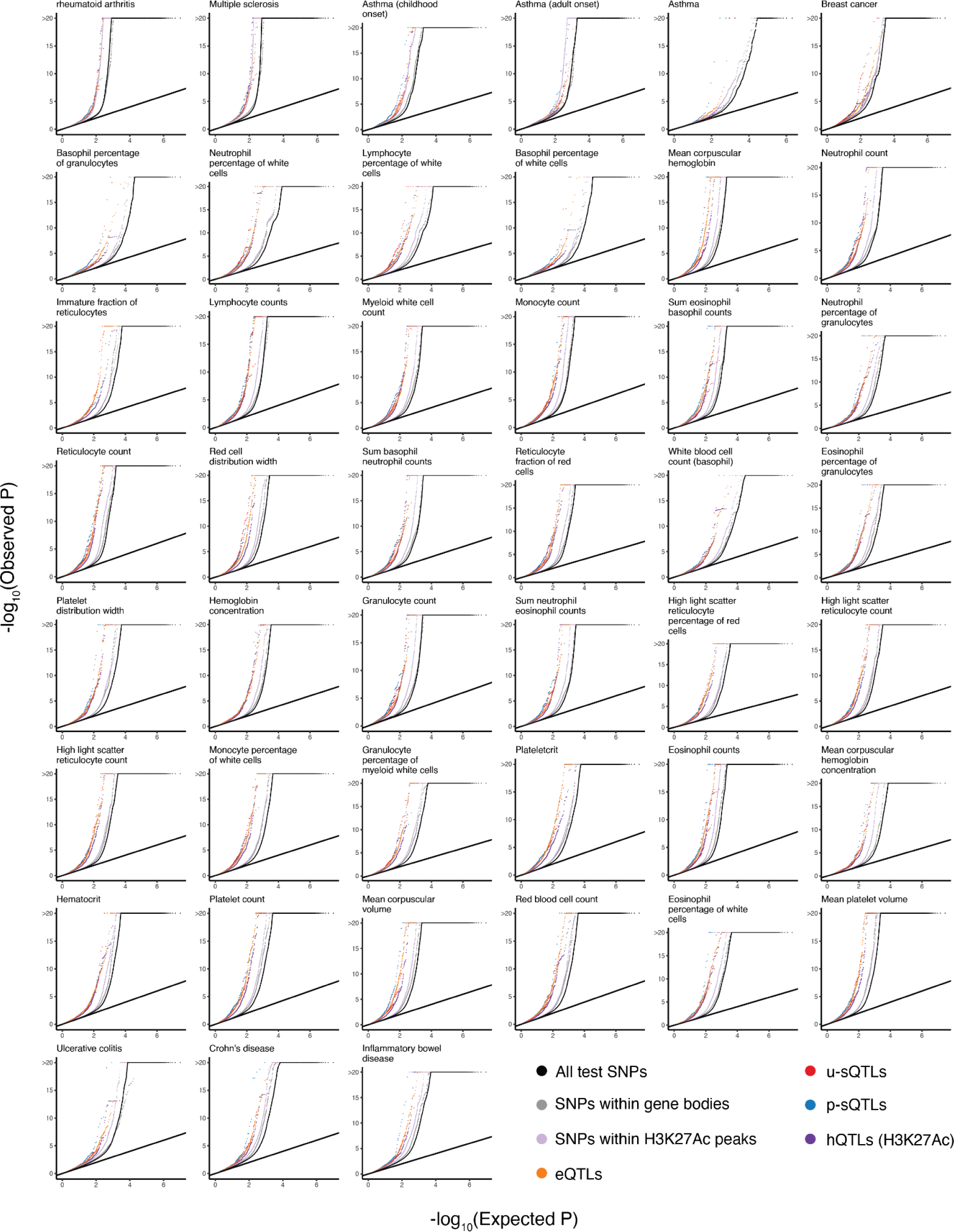
Enrichment of GWAS signal amongst molQTLs. QQ-plot of GWAS association signals for 45 blood and immune related traits, for different sets of SNPs. QTLs refer to the top SNP for each class of molQTLs. u-sQTLs and p-sQTLs represent the sQTLs that affect an unproductive splice junction or only productive (protein-coding) splice junctions, respectively.

**Fig S18.**
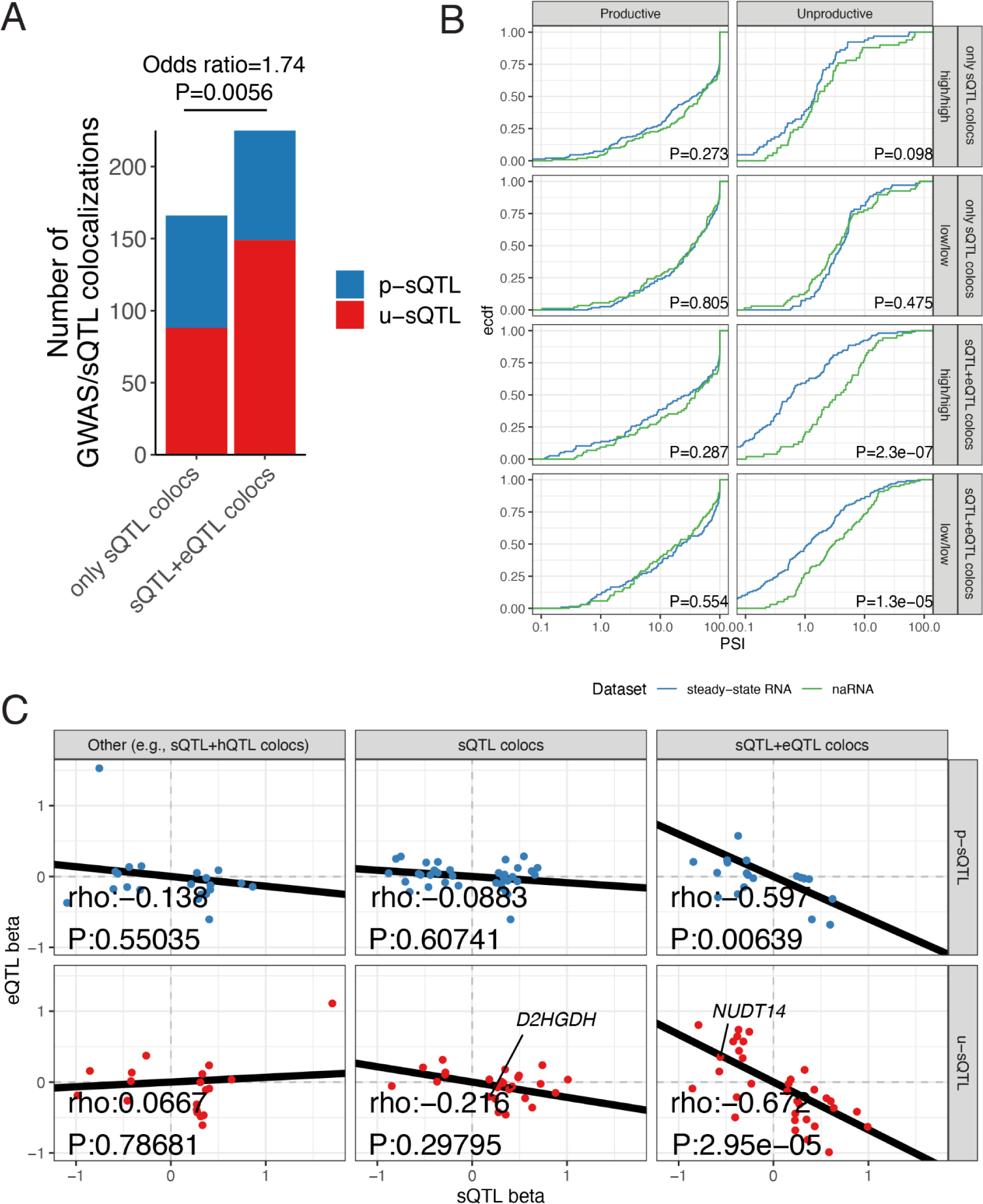
GWAS/sQTL colocalizations that also colocalize with eQTL have characteristics consistent with AS-NMD. (A) sQTLs/GWAS colocalizations are more likely to come from u-sQTLs than p-sQTLs if the host gene eQTL also colocalizes in multi-trait colocalization analysis. P value from hypergeometric test. (B) PSI distribution of introns as cumulative distribution plot for sQTLs that colocalize with eQTL and GWAS (sQTL+eQTL colocs) versus those that only colocalize with GWAS (sQTL colocs). We estimate PSI by averaging across samples with shared genotypes, either high/high genotypes or low/low genotypes (thus, avoiding confounding PSI estimates with different allele frequencies between datasets). sQTLs in unproductive introns that are sQTL+eQTLs have smaller PSI in steady-state RNA than naRNA, consistent with splicing-mediated decay at these GWAS loci transcripts. (C) Effect sizes of sQTLs that colocalize with GWAS, grouped by whether the GWAS signal also colocalizes with an eQTL (sQTL+eQTL colocs), whether it solely colocalizes with sQTL, or whether it also colocalizes with some other combination of traits (i.e., sQTL + hQTL) in multi-trait colocalization analysis. Each junction is plotted once, even if it colocalizes with multiple GWAS loci across multiple traits. sQTLs in *NUDT14* and *D2HGDH* are highlighted as examples with effect directions consistent with AS-NMD (wherein u-sQTL allele that associates with increase in splicing of unproductive junction also associates with decrease in gene expression levels).

**Fig S19.**
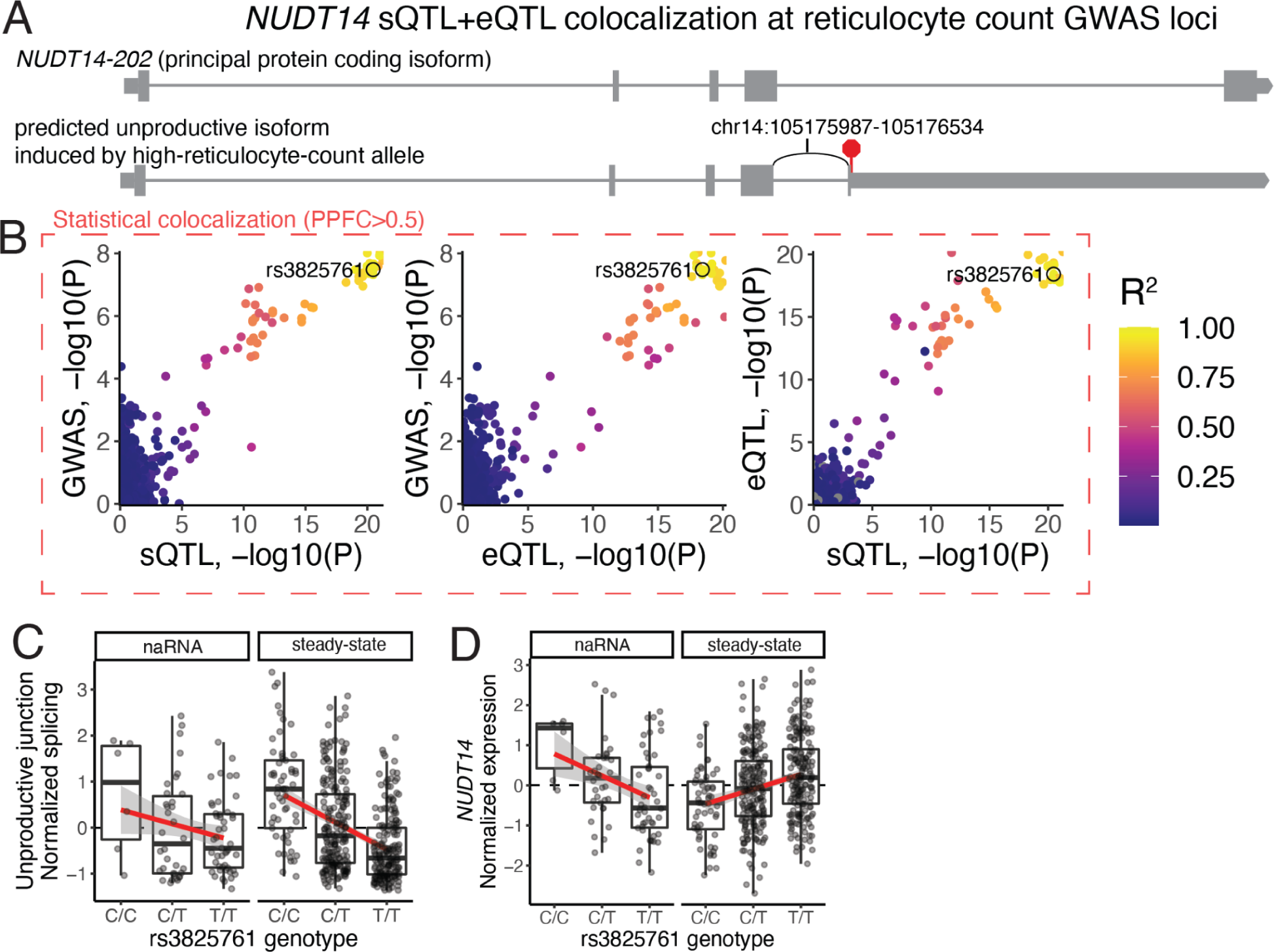
*NUDT14* u-sQTL regulates NUDT14 expression, likely contributing to reticulocyte count. *NUDT14* eQTL and sQTL. (A) Gene structure of *NUDT14-202*, the protein-coding isoform marked as the principal isoform by Gencode. Thick exonic regions mark the open reading frame. Using that isoform as a reference, we predicted the u-sQTL splice junction (labeled arc) that colocalizes with reticulocyte-count GWAS signal to introduce a premature stop codon (red octagon), created a transcript with a long 3’ UTR, inducing NMD. (B) Pairwise scatter plots depict the association between the GWAS signal, *NUDT14* eQTL signal, and chr14:105175987-105176534 junction sQTL signal. Each point is a SNP. All three traits colocalize in a single trait cluster in multi-trait colocalization (posterior probability of full colocalization, PPFC > 0.5). SNPs colored according to linkage disequilibrium relative to the top fine-mapped SNP (rs3825761). (C) *NUDT14* sQTL boxplots grouped by genotype show that the up-regulating effect of the C allele on the unproductive splice junction is present in steady-state RNA and naRNA, while the (D) down-regulating effect of the C allele on *NUDT14* expression is present in steady-state RNA but not naRNA, consistent with co-transcriptional splicing and post-transcriptional regulation by NMD.

**Fig S20.**
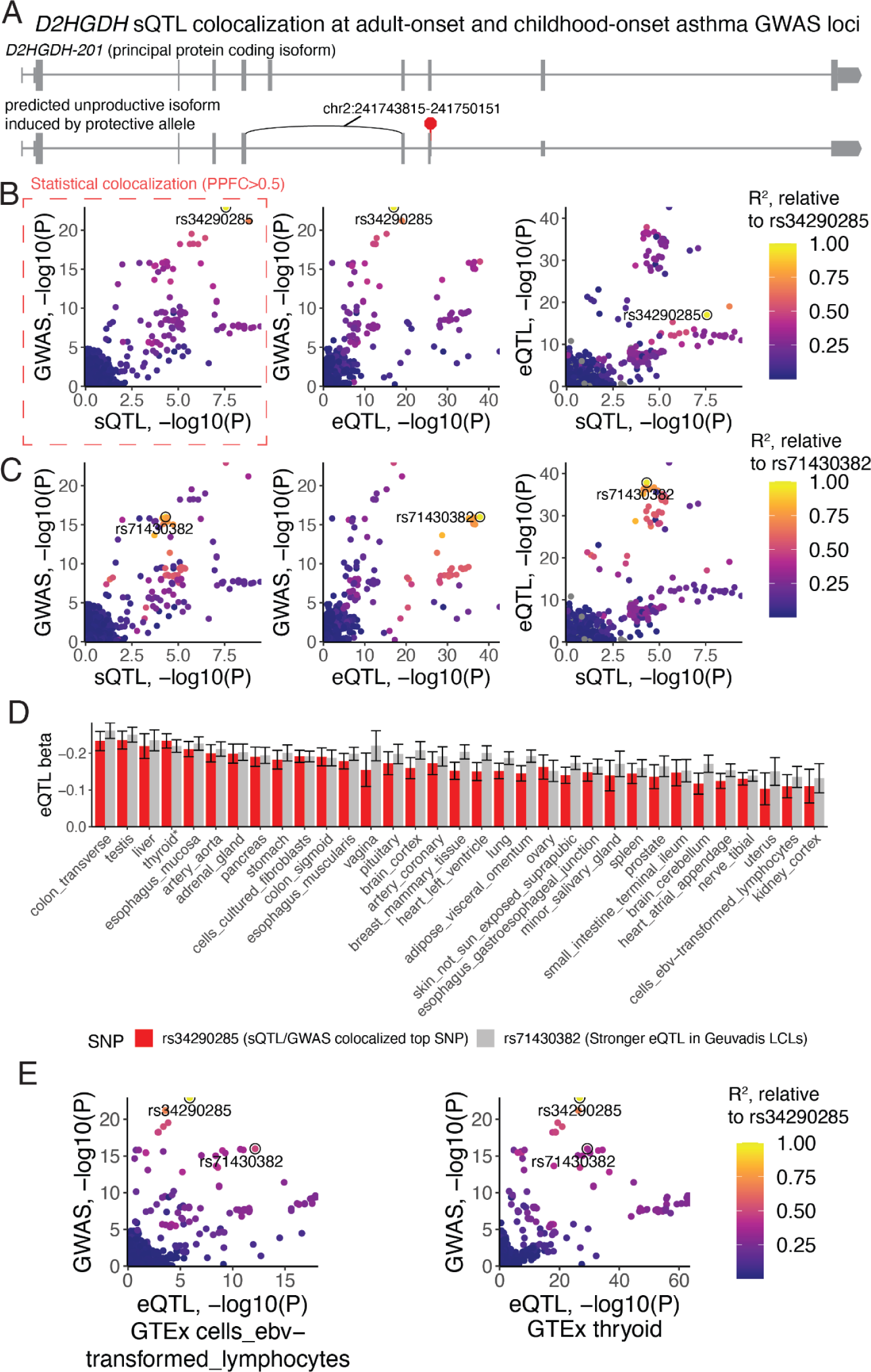
*D2HGDH* u-sQTL likely underlies asthma risk GWAS signal, as opposed to the stronger independent eQTLs in LCLs. (A) Gene structure of *D2HGDH-201*, the protein-coding isoform marked as the principal isoform by Gencode. Thick exonic regions mark the open reading frame. Using that isoform as a reference, we predicted the u-sQTL splice junction (labeled arc) that colocalizes with reticulocyte-count GWAS signal to frameshift, creating a premature stop codon (red octagon), inducing NMD. (B) Pairwise scatter plots depict the association between the GWAS signal, *D2HGDH* eQTL signal, and chr2:241743815-241750151 junction sQTL signal. Each point is a SNP. Only the GWAS signal and the sQTL trait cluster in multi-trait colocalization (posterior probability of full colocalization, PPFC > 0.5). SNPs colored according to linkage disequilibrium relative to the top fine-mapped SNP for the colocalized traits (rs34290285). Inspection of the GWAS by eQTL scatter plot revealed stronger eQTL SNPs with considerable GWAS signal, including rs71430382 and tightly linked SNPs which we highlight in (C): Same as (B) but SNPs colored by linkage relative to rs71430382. (D) The relative eQTL effects (standardized units scaled within each tissue separately) of rs34290285 and rs71430382 on *D2HGDH* expression estimated across tissues from GTEx data. *Thyroid is the tissue in which the rs34290285 has the strongest estimated effect relative to rs71430382. (E) Similar to B, with but utilizing eQTL signals from GTEx tissues: cells_ebv_transformed_lymphocytes (LCLs) and thyroid, highlighting that the colocalization between GWAS signal and *D2HGDH* eQTL signal due to rs34290285 is more plausible in thyroid.

**Fig S21.**
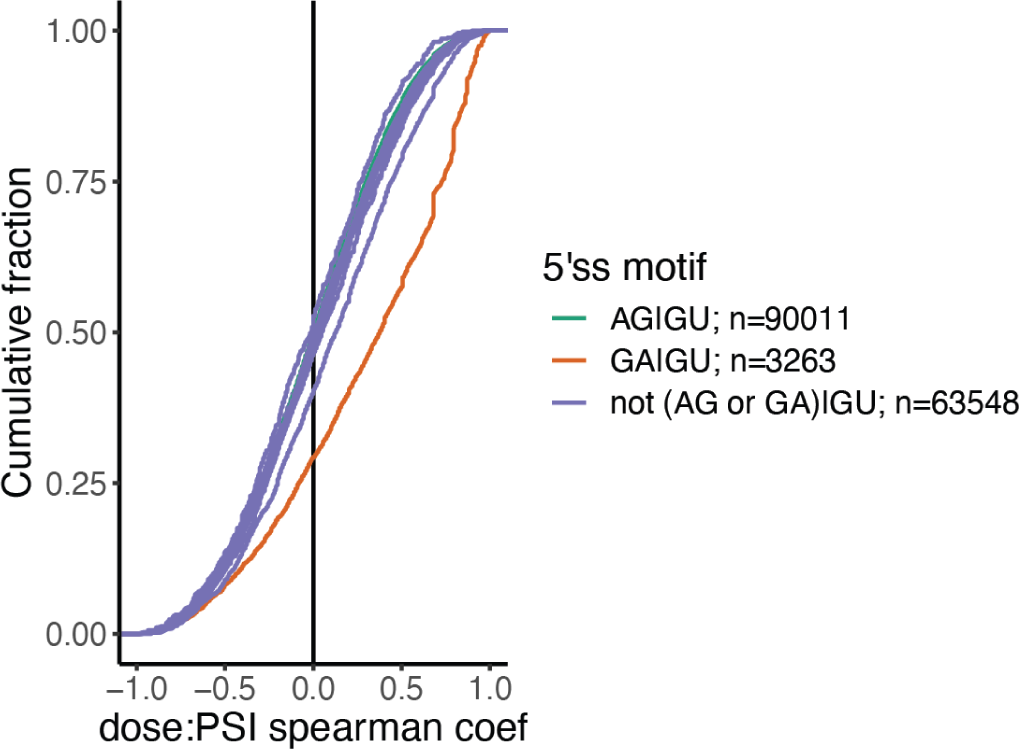
Genomewide activation of GA|GU 5’ splice sites. Cumulative distribution of dose vs splicing PSI spearman correlation coefficients amongst introns, grouped by their 5’ splice site motif. All 5’ splice site motifs contain GU at position +1:+2. 16 possible dinucleotides at position −2:-1 are plotted. The most common sequence (AG|GU, which forms Watson-Crick base-pairs with U1 snRNA) has a similar number of positive and negative correlations. Only GA|GU introns as a group have a noticeable shift in distribution towards positive correlations between dose and splicing rate. The 641 introns with significant (FDR<10%) dose-response correlation coefficients, corresponding with a spearman coefficient >0.775) were chosen for further analysis.

**Fig S22.**
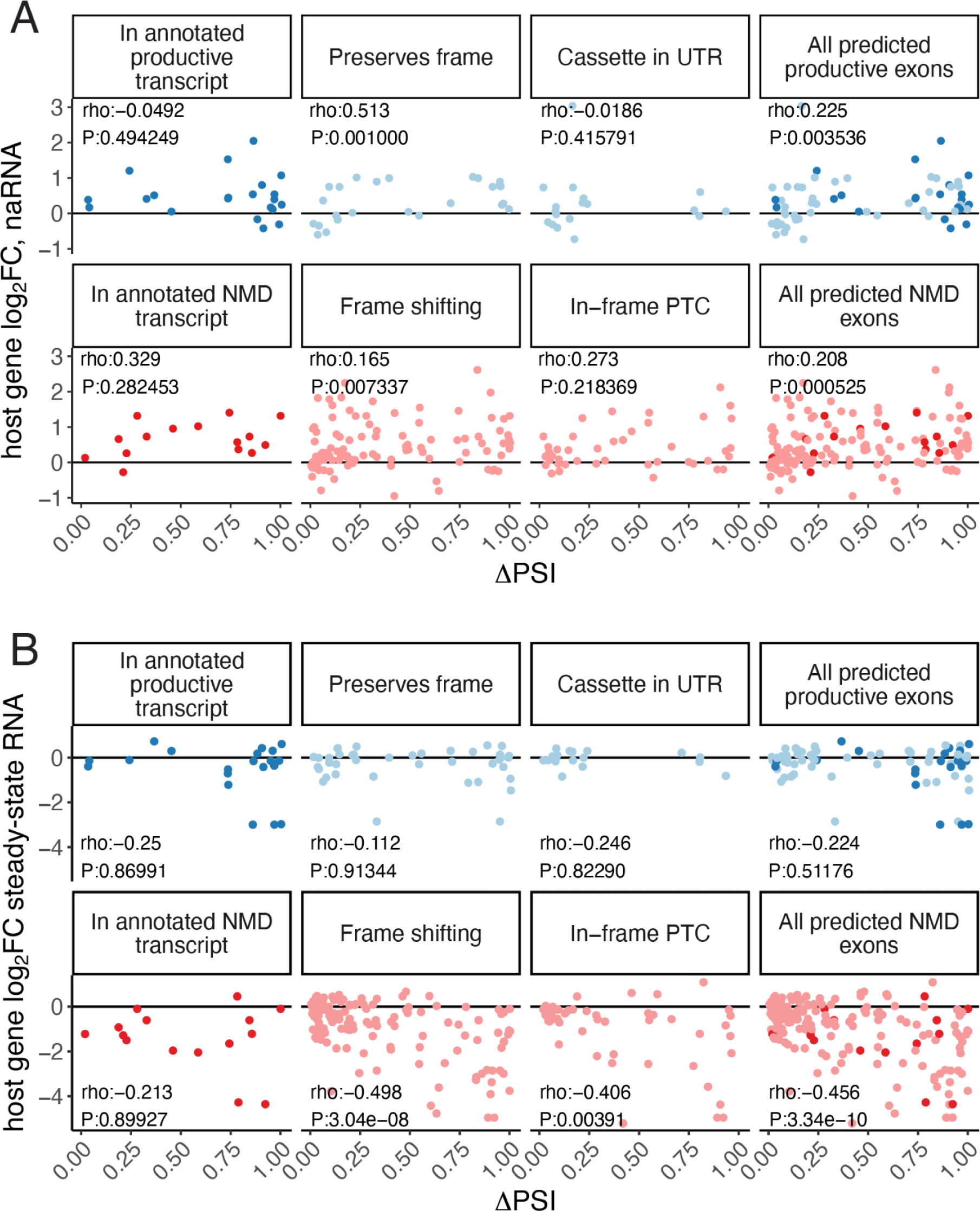
Expression and splicing effects of risdiplam-induced exons. (A) The change in percent cassette exon spliced-in (ΔPSI) vs the host-gene expression as measured in naRNA at 3160nM risdiplam dose for 305 induced cassette exons. Exons are grouped into facets for each possible reason the exon is predicted as either NMD-inducing (red) or stable (blue). Annotated exons are darker colors. In general, no correlation is observed in between splicing of cassette exons and host-gene expression in either NMD-inducing or stable exons. (B) Same as A, but effects measured in steady-state RNA. In general, a negative correlation is observed for NMD-inducing-, but not stable cassette exons. Correlations summarized with Spearman’s rho correlation coefficient and P value.

**Fig S23.**
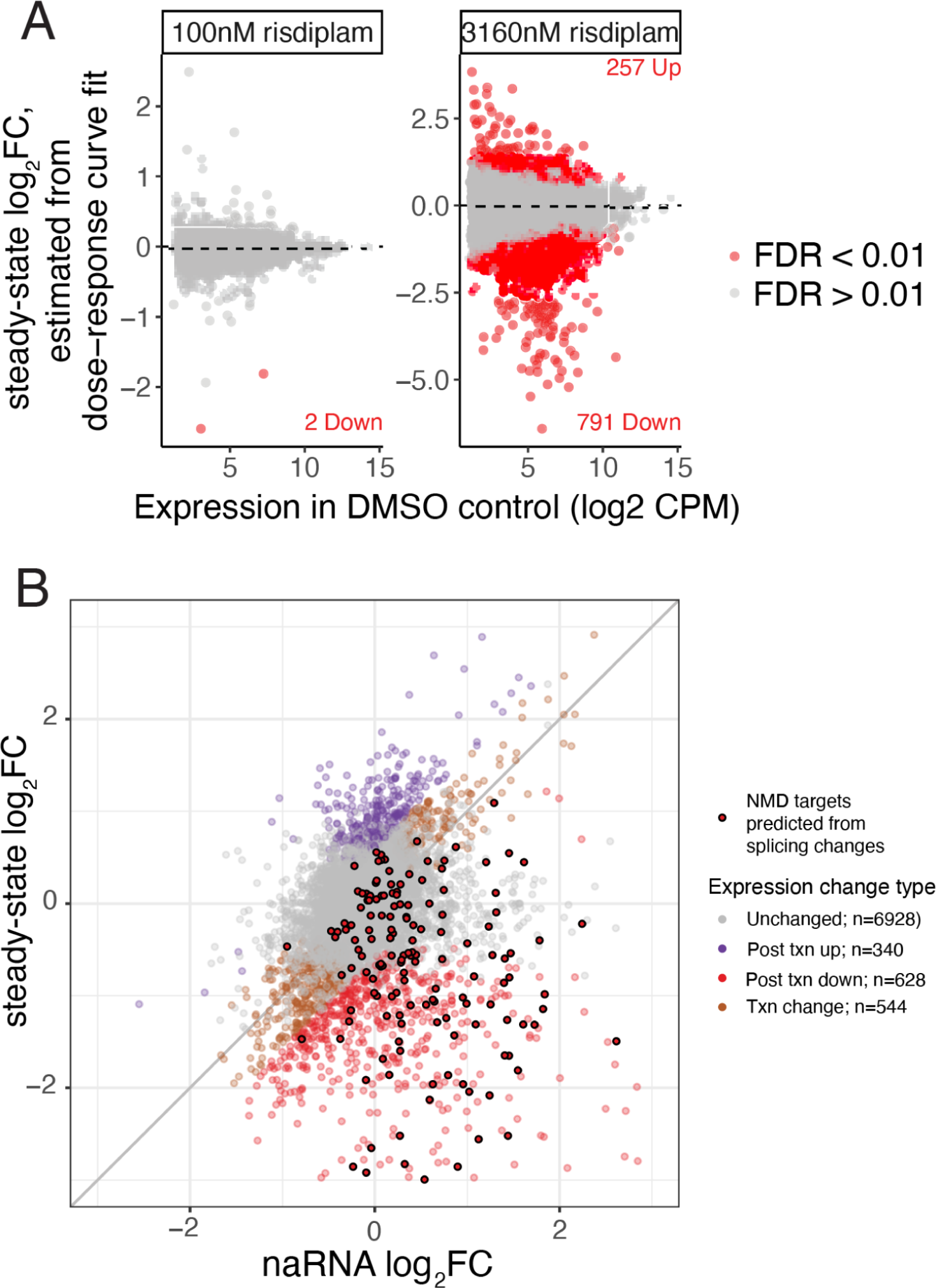
Risdiplam primarily induces down-regulating-post-transcriptional changes. (A) MA-plot of differential expression as estimated from steady-state RNA upon treatment with low or high dose of risdiplam. Number of significant up- or down-regulated genes is labeled. (B) Genes are classified by their effect size and significance (FDR<10%, STAR Methods) of expression changes as measured in naRNA or steady-state RNA-seq after being treated with 3160nM risdiplam. Transcription (Txn)-based gene expression changes are defined as having similar and significant effects as measured in naRNA and polyA RNA. Genes regulated post-transcriptionally have stronger effects in steady-state RNA. There are more post-transcriptionally down-regulated genes than up-regulated genes, suggesting risdiplam-induced splice sites more often destabilize than stabilize host transcripts. There is significant overlap of 219 NMD targets predicted from annotation of induced cassette exons among the post txn down regulated genes (Odds Ratio=14.0, P<2×10^-16^, hypergeometric test).

**Fig S24.**
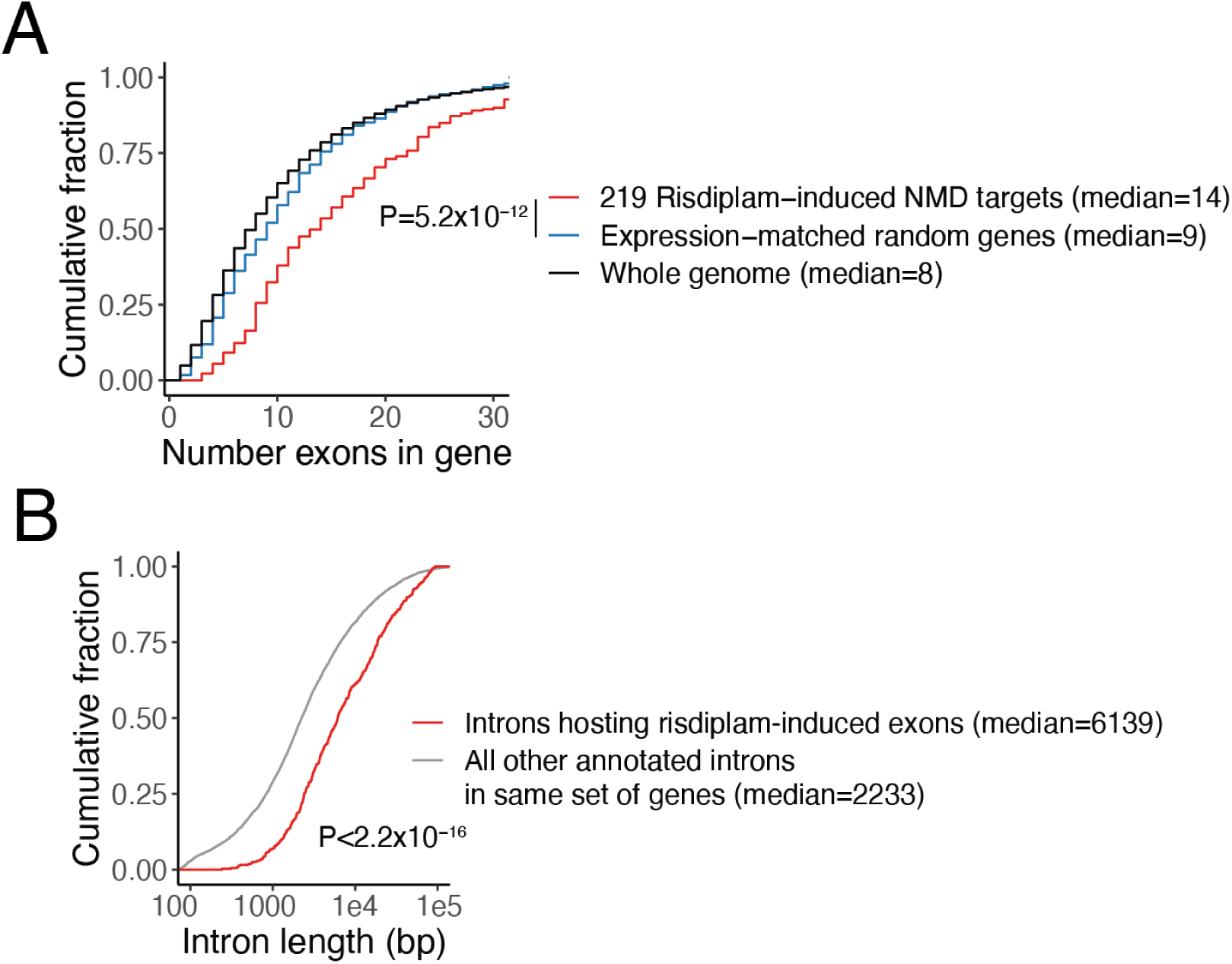
Risdiplam induced NMD exons are more likely to occur in long genes and introns. (A) Cumulative distribution of the number of exons in the gene (using the highest expressed protein_coding transcript isoform as a reference) for genes with a risdiplam-induced exon, a similarly sizes set of expression-matched set of genes, or all protein_coding genes in the genome. P value from Mann Whitney U-test. (B) Cumulative distribution of the length of introns that host risdiplam-induced exons, or as a control, all other annotated introns in the same set of genes. P-value from Mann Whitney U-test.

**Fig S25.**
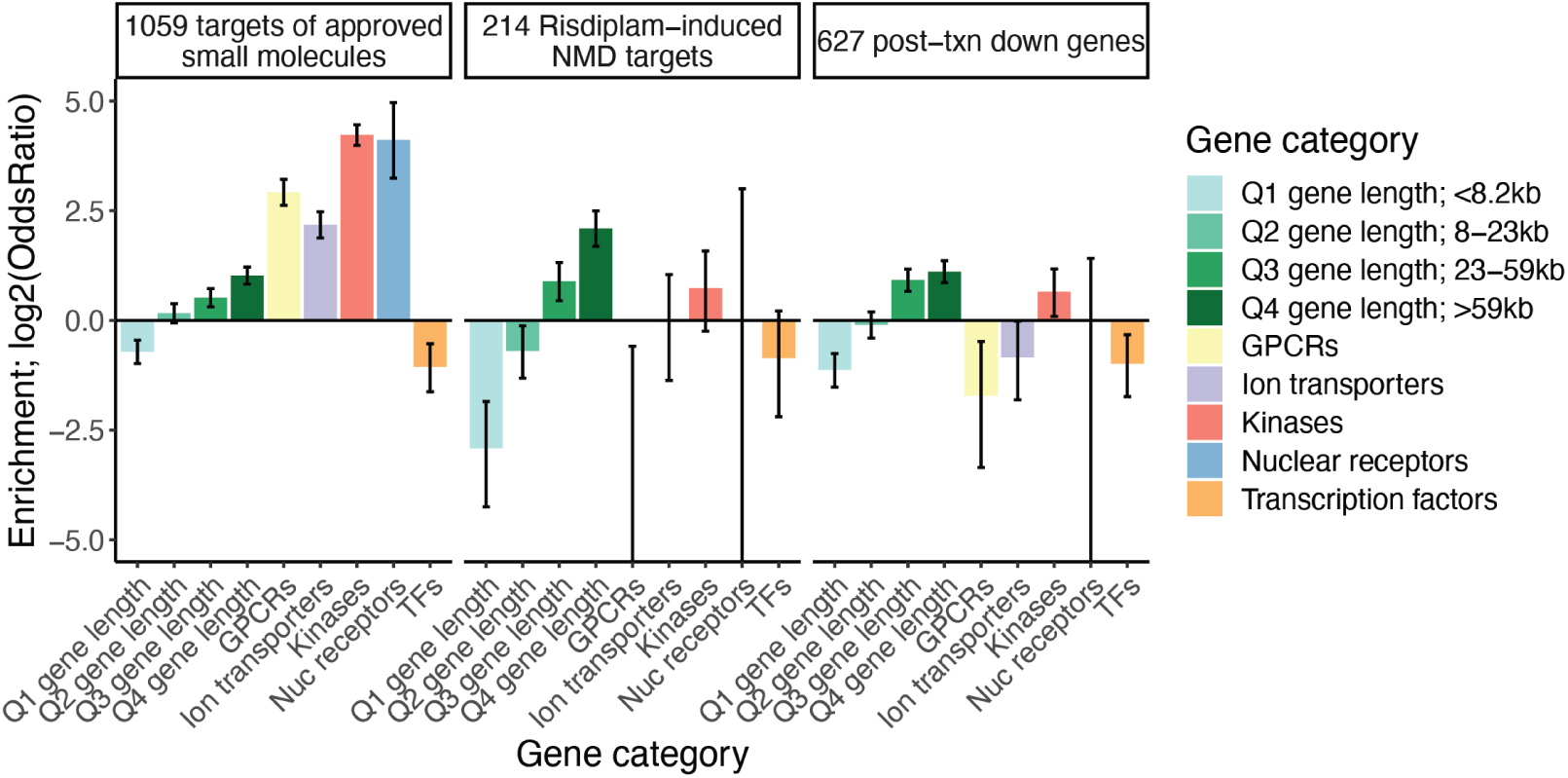
Risdiplam induced NMD targets are enriched in long genes, rather than particular protein classes. The enrichment of sets of genes among approved small molecule targets (small molecule drugs that primarily function at the level of protein binding), risdiplam-induced predicted NMD targets (defined by identification of and annotation of splicing changes), and a larger set of risdiplam-induced post-transcriptionally down-regulated genes (defined by changes in gene expression in naRNA and steady-state RNA. Gene sets defined by quartiles of gene length, and gene ontology categories (STAR Methods). Error bars represent 95% Confidence intervals.

**Fig S26.**
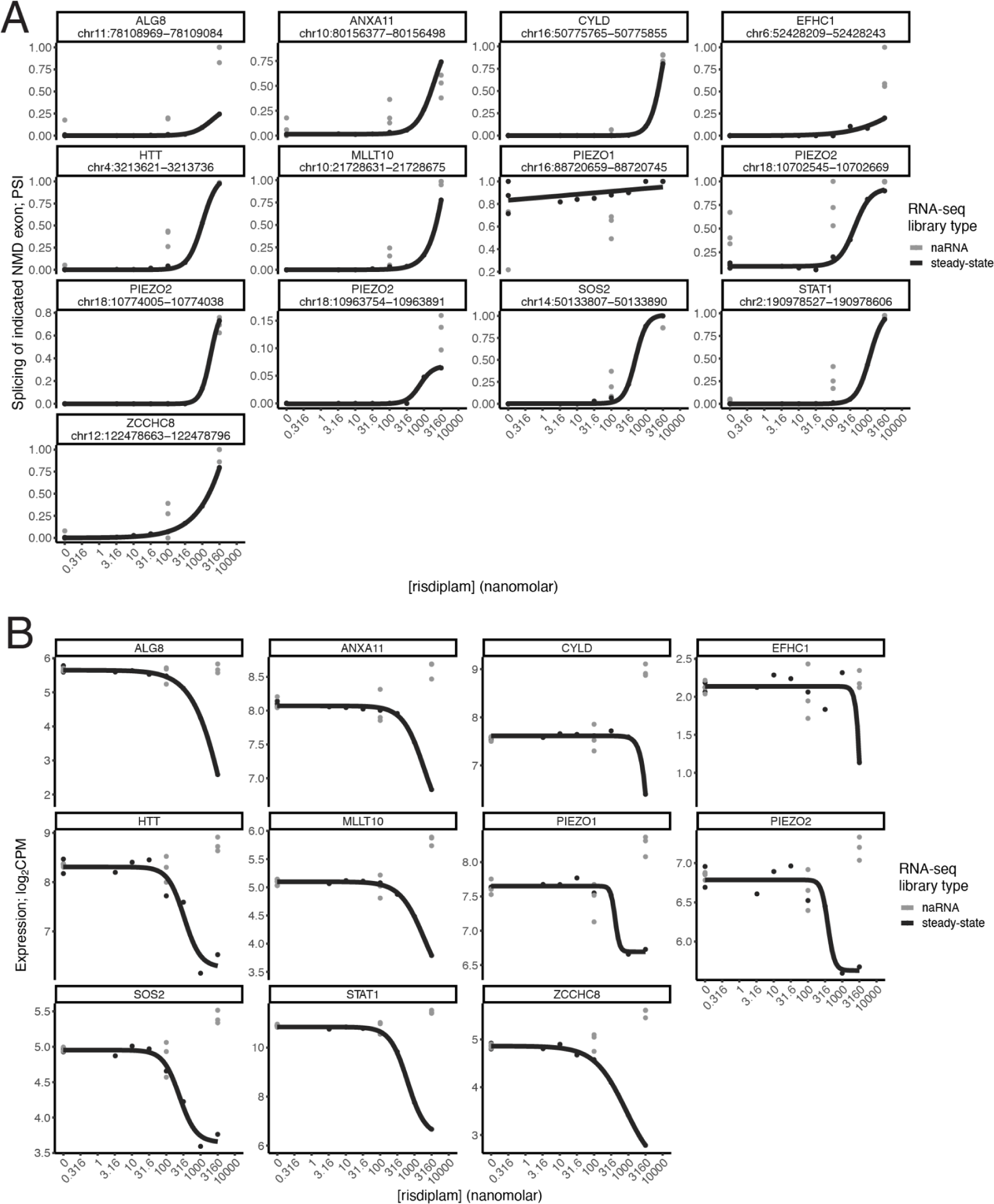
Dose-response curves of disease genes. (A) The set of 214 significantly down-regulated genes which have an identifiable splicing change predicted to induce NMD were intersected with a set of n disease genes which may produce therapeutic benefit if down-regulated. Disease genes are defined by the presence of dominant negative alleles in the OMIM database^108^. Dose-response curve shown for the splicing of the predicted NMD-inducing cassette exon. Dose-response log-logistic model fit based on PSI estimates in polyA RNA-seq, while PSI estimates from naRNA-seq samples are shown as lone points. (B) similar to (A), but measuring host-gene expression for the induced cassette exons. As expected, the down-regulating effect is polyA-specific, consistent with post-transcriptional regulation by NMD.

## Notes

### Competing Interest Statement

The authors have declared no competing interest.

### Summary of Updates

We updated the title, abstract, as well as a small part of our manuscript based on feedback from our initial biorxiv post. We also added an author that we previously omitted.

