## Supplementary Notes 1-2 for "Global impact of aberrant splicing on human gene expression levels"

### Supplementary Note 1: Contribution of alternative splicing coupled with nonsense-mediated decay to post-transcriptional regulation of gene expression levels

**In this supplementary note, we use linear regression to explore the impact of NMD in gene expression. We find that at least ~9% of the variance in post-transcriptional gene regulation in RNA levels across genes is explained by AS-NMD.**

Standard RNA sequencing measures polyadenylated steady-state mRNA molecules. These mRNA have undergone splicing, and most transcripts targeted by nonsense-mediated decay (NMD) have already been degraded[1]. In contrast, chromatin-associated RNA sequencing captures nascent RNA before degradation[2]. and during co-transcriptional splicing. We hypothesize that transcripts targeted by NMD explains part of the expression differences between standard RNA-seq, and nascent RNA sequencing (naRNA-seq). 4sU labeled RNA sequencing captures recently transcribed cytosolic RNA[3]. We expect recently transcribed RNA to have a higher portion of NMD junctions than standard RNA-seq, but less than naRNA-seq. Indeed, we observe that genes with high % NMD junction reads in naRNA have a higher log2FC between naRNA and RNA-seq, than genes with low % NMD junction reads in naRNA (Main Text, Figure 1G). This effect still exists but it is greatly diminished when comparing 4sU labeled RNA-seq with standard RNA-seq. A lingering question is whether this higher percent of NMD junction reads has a global effect in gene expression as RNA matures. Here we implement a simple regression model to show that at least ~9% of the variance in degradation rates observed across genes can be attributed to NMD activity levels as measured by % of unproductive junctions. This is likely an underestimate due to confounding factors and regression dilution. Additionally, technical differences are also expected between naRNA and steady-state RNA measurements further biasing our estimate downwards.

##### Correlation in gene expression across different RNA-seq assays

We normalized gene expression in the naRNA-seq, 4sU labeled RNA-seq, and standard RNA-seq data to RPKM as described in the methods. For each protein coding gene, we consider the median log2 RPKM expression across all the samples in each assay. We observed that gene expression is highly correlated among all RNA-seq assays, with a Pearson r of 0.85 between naRNA-seq and standard RNA-seq (Supplementary Note 1 Figure 1A). Gene expression in 4sU labeled RNA-seq has a Pearson r of 0.87 and 0.97 with naRNA-seq and standard RNA-seq, respectively (Supplementary Note 1 Figure 1B,C). This corroborated the notion that 4sU labeled RNA-seq captures RNA at a stage between nascent RNA and steady-state mature mRNA.

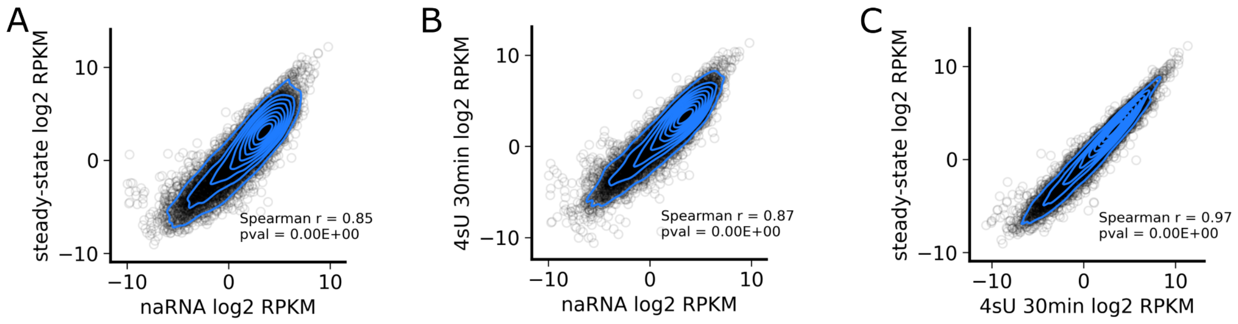


Supplementary Note 1 Figure 1: Correlation in gene expression levels measured using different RNA sequencing assays. (A) naRNA-seq vs standard RNA-seq. (B) naRNA-seq vs 4sU labeled (30 minutes) RNA-seq. (C) 4sU labeled (30 minutes) RNA-seq vs standard RNA-seq.

#### Gene expression and gene length are correlated with NMD transcript levels

The percent of NMD splice junction reads in a nascent RNA is negatively correlated with a gene's expression (Main Text, Figure 1D). This correlation strengthens across the RNA lifetime (Supplementary Note 1 Figure 2), as the effects of degradation become increasingly more prominent in increasingly mature RNA (Main Text, Figure 1G).

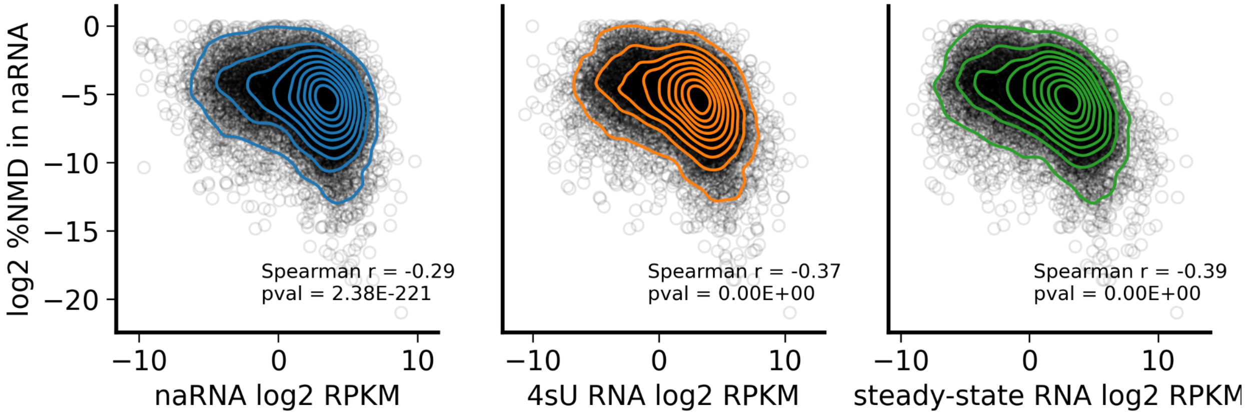


Supplementary Note 1 Figure 2: Correlation between the percent of NMD junction reads in naRNA-seq, and gene expression across the multiple RNA-seq assays.

Standard RNA sequencing and 4sU labeled RNA sequencing were both performed using polyA selection, which is known to introduce a 3' sequencing bias. Accordingly, gene length is negatively correlated with gene expression in both 4sU labeled and standard RNA sequencing, but this correlation is not observed in naRNA-seq (Supplementary Note 1 Figure 3A). Conversely, the percent of NMD junction reads in naRNA is slightly positively correlated with gene length (Supplementary Note 1 Figure 3B). For this reason, gene length is a potential confounder for the effect of NMD in the differences in gene expression between naRNA-seq and standard RNA-seq. To account for this, we used gene length as an extra covariate in our regression.

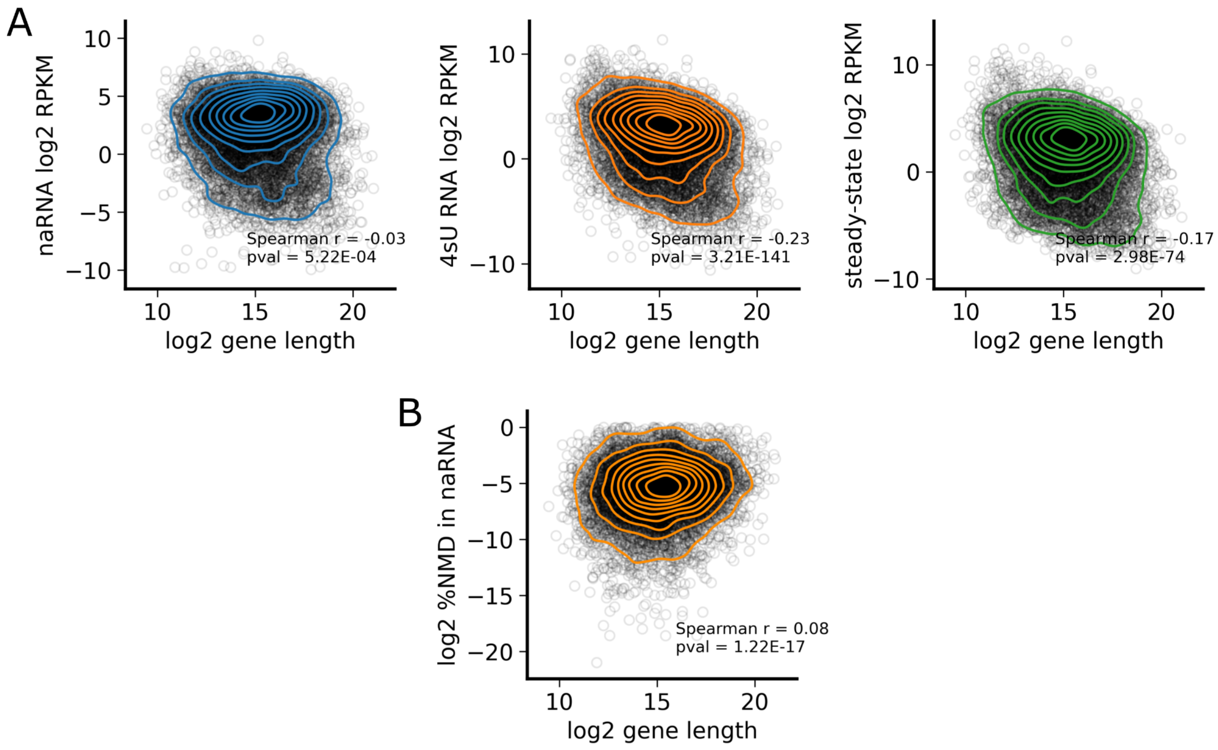


Supplementary Note 1 Figure 3: (A) Correlation between gene length and gene expression. (B) Correlation between gene length and percent of NMD junction reads in naRNA-seq.

#### Linear regression reveals global effect of NMD in gene expression

To determine the contribution of NMD to the post-transcriptional regulation of gene expression, we obtained the residuals of linear regressions of the measured expression levels of all genes at each stage of RNA maturity, versus the measured expression at earlier stages, adding the log2 of gene length as a covariate:

$$\text{steady-state RNA} \sim \text{4sU RNA} + \text{gene length},$$

$$\text{steady-state RNA} \sim \text{naRNA} + \text{gene length},$$

$$\text{4sU RNA} \sim \text{naRNA} + \text{gene length},$$

Then we performed a second regression of the residual on the log2 percent of NMD junction reads in naRNA-seq data:

$$\text{resid}_{\text{steady-state RNA v 4sU}} \sim \text{log2 \%NMD naRNA},$$

$$\text{resid}_{\text{steady-state RNA v naRNA}} \sim \text{log2 \%NMD naRNA},$$

$$\text{resid}_{\text{naRNA v 4sU}} \sim \text{log2 \%NMD naRNA},$$

To ensure that the percent of variance attributed to NMD occurs post-transcriptionally, we also performed a regression on the coverage of H3K27ac and H3K4me3 at the transcription start site of each gene, and the H3K36me3 coverage across the gene body:

$$\text{RNA-seq} \sim \text{H3K27ac} + \text{H3K4me3} + \text{H3K36me3} + \text{gene length},$$

In all cases, nonsense-mediated decay levels, as measured by % of unproductive junctions, were negatively correlated with the residuals, implying that excess NMD reduces expression levels as the RNA matures (Supplementary Note 1 Figure 4A). The slope is stronger between assays that are further apart in the regulatory cascade. Using $R^2$ scores, we found that 8.7% of the residual variance of regressing the standard RNA-seq expression versus naRNA-seq expression and gene length is explained by the percent of unproductive junction reads in naRNA-seq. This percentage is smaller (6.6%) for the residual variance between 4sU labeled RNA-seq and naRNA-seq, and for that between standard RNA-seq and 4sU labeled RNA-seq (2.3%, Supplementary Note 1 Figure 4B).

Moreover, the percent of residual variance attributed to NMD after regressing the RNA-seq assays on histone modifications is small (2.3%) in naRNA-seq, as expected. The variance explained is bigger for 4sU labeled RNA-seq, and for standard RNA-seq (7.2% and 9.5%, respectively) (Supplementary Note 1 Figure 4B), which is consistent with the residual between chromatin marks and 4sU-seq and standard RNA-seq capturing RNA degradation levels.

**Discussion**

The results of this analysis confirm that NMD is an important post-transcriptional regulatory mechanism of gene expression level. NMD has global effects across the transcriptome, suggesting that splicing plays a greater role in gene regulation than previously thought. Due to technical biases, regression dilution and other confounding factors, our analysis underestimates the percent of variance attributable to NMD.

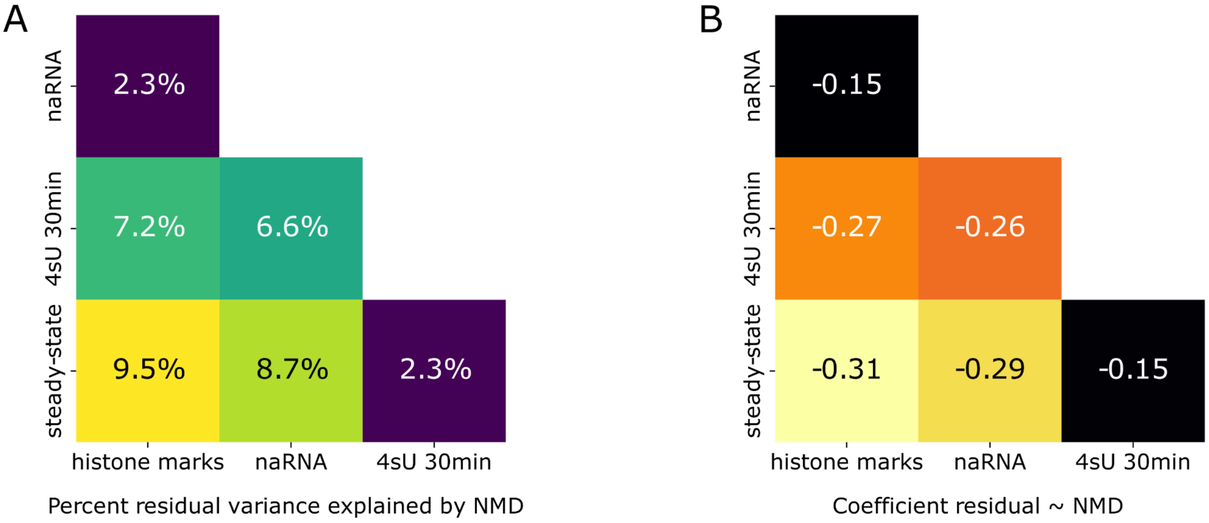


Supplementary Note 1 Figure 4:(A) Regression coefficient of the residual variance explained by the percent of NMD junction reads in naRNA-seq. (B) Percent of residual variance from Y ~ X + *gene length* explained by the percent of NMD junction reads in naRNA-seq.

#### Supplementary Note 2: Splicing of unproductive isoforms is upregulated in splicing factors and downregulated in translation factors

**Alternative splicing (AS) of transcripts targeted for nonsense-mediated decay (NMD) is a mechanism that affects gene expression post-transcriptionally. NMD is largely considered a quality control mechanism to remove mRNA molecules that encode truncated proteins. Some AS-NMD events are highly conserved, which suggests that these AS-NMD events have a regulatory role. In this Supplementary Note we use polynomial regression and Gene Ontology analysis to identify what genes produce the most or least transcripts subject to NMD. We find evidence of excess of AS-NMD in genes encoding splicing factors, and depletion in genes encoding translation factors.**

Prior to the present study, the regulated alternative splicing of NMD targeted isoforms was considered a relatively uncommon occurrence, affecting mostly splicing factors[4–7]. Our analysis of nascent RNA-sequencing data revealed that AS leading to transcripts targeted for NMD represent 2.4% of all splice junction reads. We estimated that ~20% of all mRNA molecules produced are subject to NMD. The majority of protein coding genes present NMD isoforms, with 11,328 out of 14,000 protein coding genes analyzed having at least one NMD junction read, and 6,549 genes with at least 1% and less than 20% of reads predicted to induce NMD (we consider genes with less than 20% NMD splice junction reads because higher numbers could correspond to genes with very low expression or to genes with little to no protein-coding isoforms). These results put together suggest that AS-NMD events are widespread across the transcriptome, in greater abundance than previously described. Still, one remaining question is to what extent AS-NMD events play a regulatory function in gene expression versus a mere consequence of erroneous RNA splicing.

##### Genes with highly conserved poison exons have high percentage of NMD transcripts

Arguably the best studied instance of regulated AS-NMD are the genes in the Serine/arginine-rich splicing factors (SRSF) family. The majority of genes in this family can produce isoforms with alternative splicing events - usually cassette exons, that introduce premature termination codons (PTC), known as poison exons - that lead to NMD[5,6,8,9]. Cross-linking and immunoprecipitation (CLIP) studies have shown that proteins of the SRSF family bind to their own poison exons, promoting their inclusion[8,10–12]. As a result, AS of NMD isoforms has been proposed as a post-transcriptional mechanism of gene expression auto-regulation in these genes. In addition to the SRSF family, AS events leading to NMD transcripts are enriched in splicing factors and chromatin factors[7]. These include the poison exons of the key spliceosome component SNRNP70[13], and in SMNDC1, a splicing factor involved in the assembly of the mature spliceosome[13,14]. Other splicing factors autoregulate their own activity through other means, such as MBNL1, which represses inclusion of its own exon carrying a nuclear localization signal protein domain[15].

In this paper we show that two factors are highly correlated with the PSI of NMD junctions: (i) gene expression and (ii) the evolutionary constraint of the gene (Main text Figure 2A, Figure S6-2A). As a result, lowly expressed genes dominate the highest quantiles of NMD junction PSI. To account for these confounding factors, we performed a quadratic polynomial regression of each gene's maximum junction PSI on the gene's expression in log2 RPKM, and the gene's constraint score in log2 s_het[16] as follows:

max(junction PSI) ~ a*RPKM + b*s_het + c*RPKM*s_het + d*RPKM^2 + e * s_het^2 + const

The residual of this regression indicates the deviation in the usage of NMD junctions (as measured by the maximum PSI of a NMD junction) from the expected usage based on the gene's expression and evolutionary constraint. Nine out of the eleven genes in the SRSF family with annotated splice junctions had a positive residual, indicating that they have higher NMD junction PSI than expected (Supplementary Note 2 Figure 1). Three of them: SRSF4, SRSF10 and SRSF11 had more than three standard deviations above the average residual. MBNL1, SNRNP70 and SMNDC1 also had positive residuals, with MBNL1 having more than two standard deviations above the average residual. SRSF5 and SRSF9 had a negative residual within one standard deviation from the mean. SRSF8 was excluded given that it does not have annotated protein coding splice junctions.

With the exception of SRSF12 and SMNDC1, all aforementioned splicing factors are in the highest quartile of gene expression level. And with the exception of SRSF12, all of them are also in the highest quartile of evolutionary constraint. The combination of high expression, high evolutionary constraint and high production of NMD transcripts likely played a role in why AS-NMD events in these genes have been extensively described. These results show that the highly conserved AS-NMD events in splicing factors are used significantly more frequently than in genes with similar levels of gene expression and evolutionary constraint. This is consistent with the notion that the ability of splice factors to regulate their own expression is an important example of regulated AS-NMD.

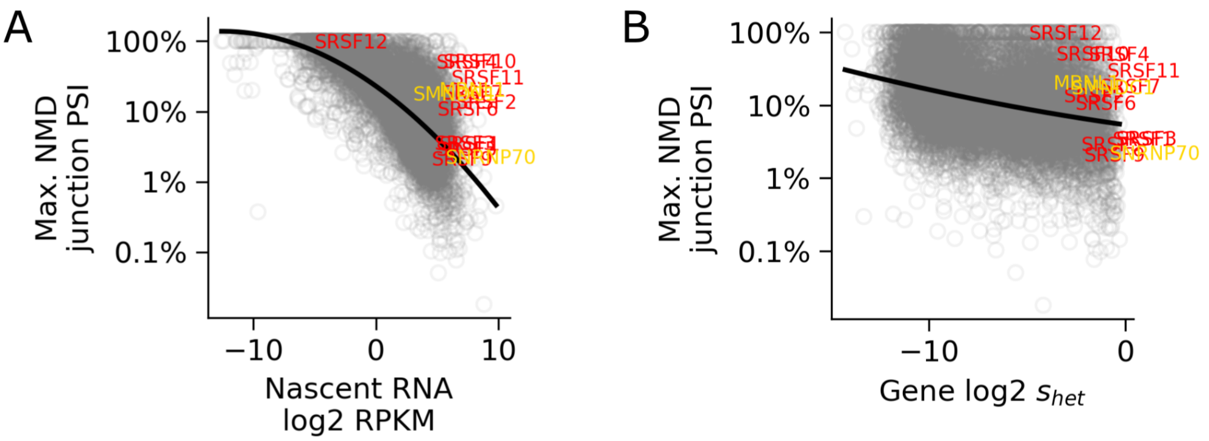


Supplementary Note 2 Figure 1: Splicing factors with highly conserved cassette exons tend to have more AS-NMD splicing than predicted by (A) gene expression and (B) evolutionary constraints alone. Gene names in red are from the SRSF genes, while names in orange correspond to MBNL1, SMNDC1 and SNRNP70. Black lines correspond to the polynomial regression of the maximum junction PSI on the corresponding X-axis feature. It is worth noticing that the polynomial regression used for the rest of the analyses was done simultaneously on the gene expression level and evolutionary constraint score.

.

#### Splicing factors have high AS-NMD, while cell cycle genes have low AS-NMD

To expand our previous results, we investigated what type of genes are the most affected by AS-NMD events, and which genes are the least affected. We reasoned that genes with the lowest residuals from the polynomial regression are the least affected by AS-NMD, possibly due to AS-NMD splicing suppression. Meanwhile, genes with the highest residual score could be genes whose AS-NMD is upregulated.

From the 14,000 protein coding genes that we selected for our main analysis, 11,328 had AS-NMD events in naRNA-seq data. We performed Gene Ontology enrichment analysis in the bottom and top decile of genes ranked by the residual from the polynomial regression (i.e., the 1,133 genes with the lowest regression residual, and the 1133 genes with the highest regression residual). For this, we used the Enrichr[17–19] implementation in GSEApy[20]. We used the Biological Process subset of the C5: Gene Ontology signature collection from the MSigDB database[21,22]. We found that the top enriched tags on the bottom decile are associated with peptide and amide biosynthesis, ribosome biogenesis, and translation (Supplementary Note 2 Figure 2A). In contrast, the most enriched tags on the top decile are associated with RNA metabolism, processing, and splicing, as well as protein modification (Supplementary Note 2 Figure 2B).

To corroborate these results, we also explored Gene Ontology enrichment in groups of highly expressed genes with high evolutionary constraint. From these, we selected 1,225 genes that are both in the top quartile of expression in our naRNA-seq data, and in the top quartile of the s_het evolutionary constraint metric from GeneBayes[16]. From these genes, we selected the genes at the bottom or top quartile of NMD junction PSI (306 genes on each quartile). Once again, we found that genes in the bottom quartile of NMD splicing junctions are enriched for translation and protein biosynthesis tags (Supplementary Figure Figure 2C), while genes on the top NMD quartile are enriched for tags in mRNA metabolism and splicing (Figure 2D).

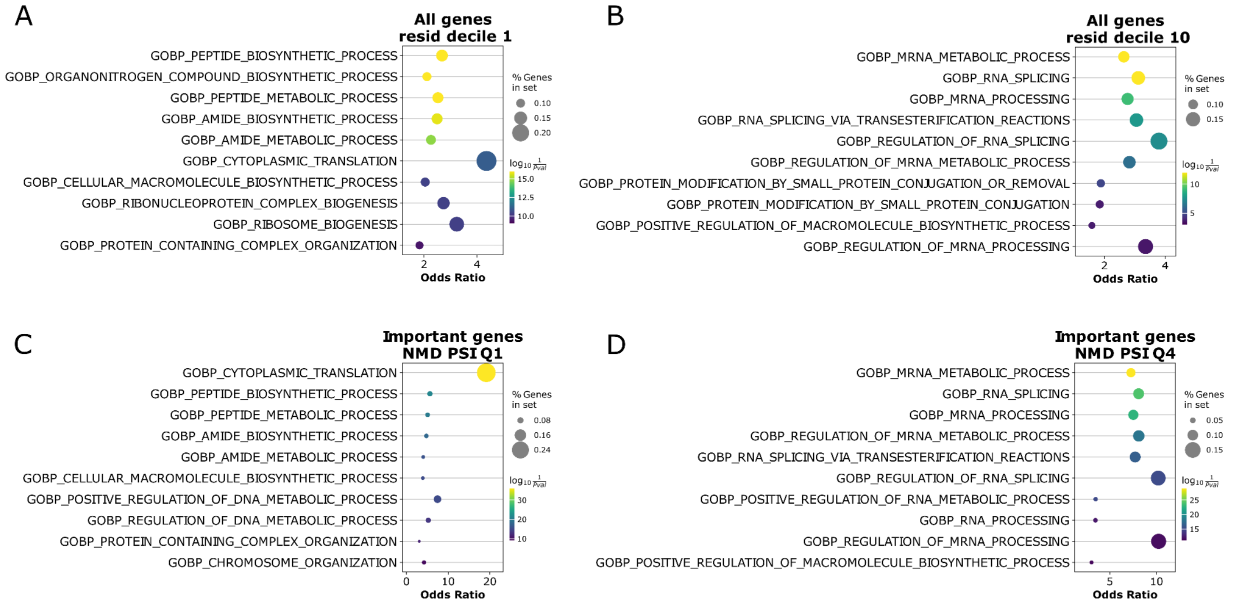

Supplementary Note 2 Figure 2: Gene Ontology analysis using MSigDB's C5 collection of tags, selected for Biological Process events. (A) All genes in the bottom decile of polynomial residual. (B) top decile of residual. (C) Tags in top expressed and evolutionary conserved genes, in the bottom NMD junction PSI. (D) Tags in top expressed and evolutionary conserved genes, in the top NMD junction PSI.

To further explore the contribution of specific types of genes, we selected genes with Gene Ontology Biological Process tags associated with transcription (transcription factors), RNA splicing (splicing factors), chromatin remodeling (chromatin factors) and RNA translation into proteins (translation factors), from the 1,225 genes in the top RPKM and s_het score quartiles. Genes in the top decile of the regression residual were enriched for splicing factors (enrichment = 2.43, hypergeometric test p-value = $1.9 \times 10^{-7}$), while genes in the bottom decile were enriched for translation factors (enrichment = 2.64, hypergeometric test p-value = $2.1 \times 10^{-8}$; Supplementary Note 2 Figure 3A). Previous studies reported that regulated AS-NMD events are more common in splicing factors, RNA binding proteins, and chromatin factors. Our results further show that indeed splicing and chromatin factors have a larger percent of NMD junction reads (Supplementary Note 2 Figure 3B) and a larger NMD junction PSI (Supplementary Note 2 Figure 3C) than other types of genes.

On average, the most abundant NMD junction of a gene contributes 45.1% of all NMD junction reads, while 16.4% are contributed by reads in the fifth and lower rank (Main figure 2B). This suggests that splicing is error-prone and that genes tend to produce multiple NMD junction reads. Splicing factors have a higher contribution from the top ranked NMD junction, with 52.3%, and a smaller contribution from the fifth and lower ranked NMD junctions, with 9.6% (Supplementary Note 2 Figure 3D). The difference is even higher in SRSF genes, with 62.3% of the NMD junction reads on average coming from the top junction, and only 4.1% from the fifth and lower ranked NMD junctions. Translation factors also have a higher than average contribution from the top ranked NMD junction, while transcription and chromatin factors have a higher than average NMD contribution from junctions ranked fifth and lower. Interestingly, coding transcripts from splicing factors and translation factors tend to have fewer introns than chromatin and transcription factors (Supplementary Note 2 Figure 3E).

These results show that, despite both splicing and chromatin factors having more NMD junction reads overall, splicing factors have a larger contribution from a single AS-NMD event, while chromatin factors have a more uniform contribution of splice junctions. This suggests that there are AS-NMD events in splicing factors that are consistently spliced, which implies that the process is regulated. In contrast, in chromatin factors there are multiple AS-NMD events contributing to the overall production of NMD isoforms. These could be the result of errors in the splicing of a larger number of introns per transcript. Finally, our results also provide evidence that AS-NMD events are suppressed in translation factors.

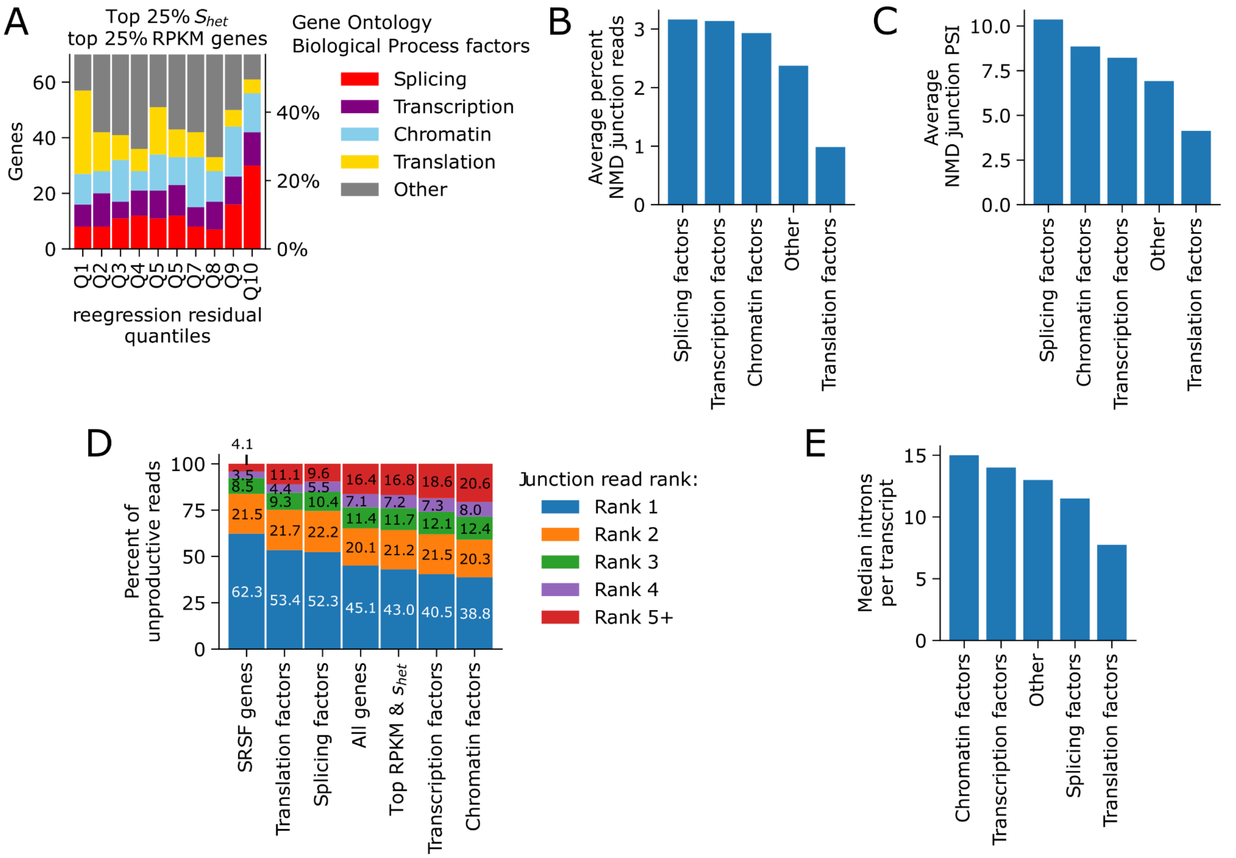


Supplementary Note 2 Figure 3: AS-NMD in splicing and translation factors. (A) percent of genes from each tag present in polynomial regression residual. For clarity, the barplot shows up to 70 genes per decile, out of a total of 123. (B) Average percent of NMD junction reads on multiple gene categories. (C) Average percent of NMD junction PSI on multiple gene categories. (D) distribution of NMD junction ranks across multiple gene categories. (E) Median number of introns per coding transcript by each gene category.

#### Haploinsufficient genes have lower levels of AS-NMD

Genes are considered haploinsufficient if deletion or loss-of-function mutations on one single copy lead to reduction of fitness. This means that for haploinsufficient genes, the dosage or functional RNA and/or protein produced by one single copy is not enough to maintain the gene's proper function[23]. Since haploinsufficient genes are likely more sensitive to changes in gene dosage than genes that can maintain proper function with one single functional copy. We reasoned that haploinsufficient genes may exhibit fewer AS-NMD events than haplosufficient genes. To test this hypothesis, we used the haploinsufficiency prediction scores from Huang et al. 2010[24] to sort genes according to their probability of being haploinsufficient. We found that genes with high probability of being haploinsufficient in general have a lower NMD splice junction PSI (Supplementary Note 2 Figure 4A). However, this effect could be influenced by higher expression levels of haploinsufficient genes compared to haplosufficient genes (Supplementary Note 2 Figure 4B). Indeed, when we analyze genes in the top expression and evolutionary constraint quartile, the difference in splice junction PSI for the most used NMD junction is reduced, although haploinsufficient genes still have a lower PSI (Supplementary Note 2 Figure 4C).

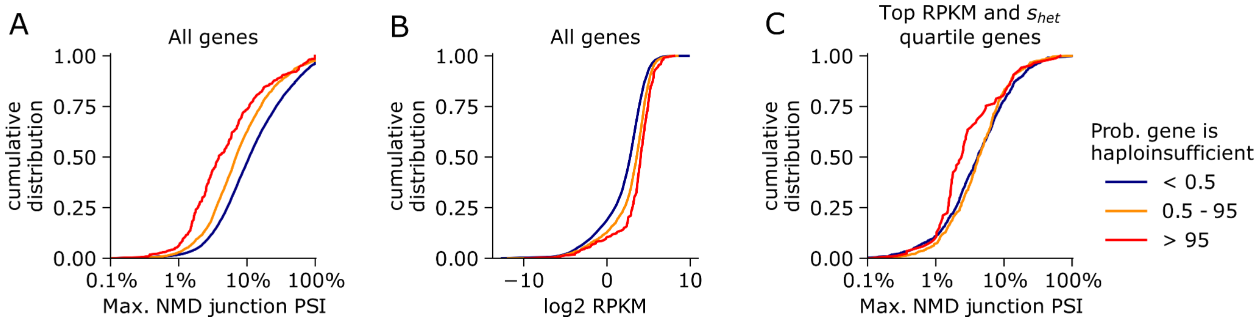


Supplementary Note 2 Figure 4. AS-NMD in haploinsufficient genes. (A) Cumulative distribution of the percent spliced-in of the highest used NMD junction in genes, stratified by their probability of being haploinsufficient. Red lines correspond to genes with high probability of being haploinsufficient. Orange lines are genes that are more likely than not to be haploinsufficient. Blue lines correspond to genes that are likely not haploinsufficient. (B) Cumulative distribution of gene expression in log2 RPKM in genes. (C) Cumulative distribution of the percent spliced-in of the highest used NMD junction in genes in the top quartiles of expression and evolutionary constraint.

In our main analysis, we demonstrate that splicing QTLs (sQTLs) affecting NMD associated splice junctions are more likely to have an effect on the gene's expression level, than sQTLs affecting protein coding junctions. Given that AS-NMD events affect haploinsufficient genes, we asked what is the impact that sQTLs affecting NMD splice junctions in haploinsufficient genes have on gene expression. We found that, although sQTLs affecting NMD splice junctions in haploinsufficient genes have a similar strength of effect than their counterparts in genes that are not haploinsufficient (Supplementary Note 2 Figure 5A), their effect on gene expression is weaker (Supplementary Note 2 Figure 5B). Interestingly, sQTLs affecting NMD junctions in haploinsufficient genes tend to decrease intron splicing rather than increase it, when comparing them with sQTLs affecting NMD junctions in genes that are not haploinsufficient (Supplementary Figure 5C). NMD sQTLs in haploinsufficient genes also tend to result in smaller changes in gene expression levels, when compared to genes that are not haploinsufficient (Supplementary Note 2 Figure 5D).

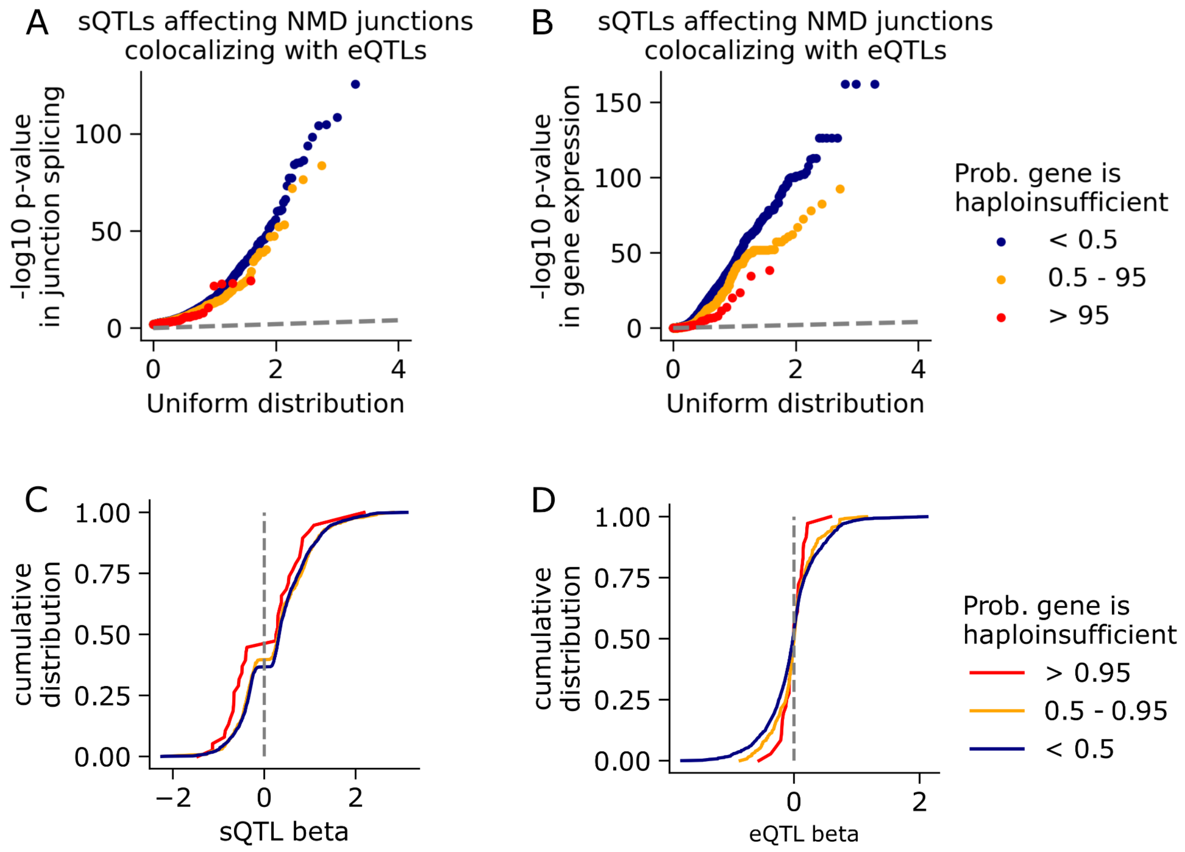


Supplementary Note 2 Figure 5: sQTLs affecting NMD junctions in haploinsufficient genes. (A) QQ plot of the p-values for sQTLs affecting NMD splice junctions, stratified by the gene's probability of being haploinsufficient. (B) QQ plot of the p-values of the effect of these sQTLs on the expression of the gene containing the NMD splice junction. (C) Cumulative distribution of the sQTL effect size on the affected NMD splice junctions. (D) Cumulative distribution of the sQTL effect size on the expression of the genes containing the NMD splice junctions.

#### Discussion

AS-NMD events are pervasive, affecting the majority of protein coding genes. Functional and evolutionary evidence suggest that some AS-NMD events are regulated and they have a role in maintaining homeostasis in the levels of gene expression, as it is the case of the highly conserved poison exons in the SRSF genes and other splicing factors. Here we show that splicing factors have a higher percentage of NMD splice junction reads, and a higher NMD splice junction PSI, than other types of genes with similar levels of gene expression and evolutionary constraint. This increase in NMD splicing comes primarily from one or two NMD splice junctions, which supports the hypothesis that AS-NMD events actively regulate the expression levels of splicing factors. In contrast, translation factors have smaller percentages of NMD splicing and lower NMD splice junction PSI than other genes with similar levels of expression and evolutionary constraint. This suggests that AS-NMD events are suppressed in translation factors.

1. Brogna S, Wen J. Nonsense-mediated mRNA decay (NMD) mechanisms. *Nat Struct Mol*

*Biol*. 2009;16(2):107-113. doi:10.1038/nsmb.1550
2. Wissink EM, Vihervaara A, Tippens ND, Lis JT. Nascent RNA analyses: tracking transcription and its regulation. *Nat Rev Genet*. 2019;20(12):705-723. doi:10.1038/s41576-019-0159-6
3. Li YI, van de Geijn B, Raj A, et al. RNA splicing is a primary link between genetic variation and disease. *Science*. 2016;352(6285):600-604. doi:10.1126/science.aad9417
4. Pickrell JK, Pai AA, Gilad Y, Pritchard JK. Noisy Splicing Drives mRNA Isoform Diversity in Human Cells. *PLOS Genet*. 2010;6(12):e1001236. doi:10.1371/journal.pgen.1001236
5. Ni JZ, Grate L, Donohue JP, et al. Ultraconserved elements are associated with homeostatic control of splicing regulators by alternative splicing and nonsense-mediated decay. *Genes Dev*. 2007;21(6):708-718. doi:10.1101/gad.1525507
6. Lareau LF, Inada M, Green RE, Wengrod JC, Brenner SE. Unproductive splicing of SR genes associated with highly conserved and ultraconserved DNA elements. *Nature*. 2007;446(7138):926-929. doi:10.1038/nature05676
7. Yan Q, Weyn-Vanhentenryck SM, Wu J, et al. Systematic discovery of regulated and conserved alternative exons in the mammalian brain reveals NMD modulating chromatin regulators. *Proc Natl Acad Sci*. 2015;112(11):3445-3450. doi:10.1073/pnas.1502849112
8. Jumaa H, Nielsen PJ. The splicing factor SRp20 modifies splicing of its own mRNA and ASF/SF2 antagonizes this regulation. *EMBO J*. 1997;16(16):5077-5085. doi:10.1093/emboj/16.16.5077
9. Lareau LF, Brenner SE. Regulation of Splicing Factors by Alternative Splicing and NMD Is Conserved between Kingdoms Yet Evolutionarily Flexible. *Mol Biol Evol*. 2015;32(4):1072-1079. doi:10.1093/molbev/msv002
10. Änkö ML, Müller-McNicoll M, Brandl H, et al. The RNA-binding landscapes of two SR proteins reveal unique functions and binding to diverse RNA classes. *Genome Biol*. 2012;13(3):R17. doi:10.1186/gb-2012-13-3-r17
11. Brugiolo M, Botti V, Liu N, Müller-McNicoll M, Neugebauer KM. Fractionation iCLIP detects persistent SR protein binding to conserved, retained introns in chromatin, nucleoplasm and cytoplasm. *Nucleic Acids Res*. 2017;45(18):10452-10465. doi:10.1093/nar/gkx671
12. Leclair NK, Brugiolo M, Urbanski L, et al. Poison Exon Splicing Regulates a Coordinated Network of SR Protein Expression during Differentiation and Tumorigenesis. *Mol Cell*. 2020;80(4):648-665.e9. doi:10.1016/j.molcel.2020.10.019
13. Saltzman AL, Kim YK, Pan Q, Fagnani MM, Maquat LE, Blencowe BJ. Regulation of Multiple Core Spliceosomal Proteins by Alternative Splicing-Coupled Nonsense-Mediated mRNA Decay. *Mol Cell Biol*. 2008;28(13):4320-4330. doi:10.1128/MCB.00361-08
14. Rappsilber J, Ajuh P, Lamond AI, Mann M. SPF30 Is an Essential Human Splicing Factor Required for Assembly of the U4/U5/U6 Tri-small Nuclear Ribonucleoprotein into the Spliceosome *. *J Biol Chem*. 2001;276(33):31142-31150. doi:10.1074/jbc.M103620200
15. Kino Y, Washizu C, Kurosawa M, et al. Nuclear localization of MBNL1: splicing-mediated autoregulation and repression of repeat-derived aberrant proteins. *Hum Mol Genet*. 2015;24(3):740-756. doi:10.1093/hmg/ddu492
16. Zeng T, Spence JP, Mostafavi H, Pritchard JK. Bayesian estimation of gene constraint from an evolutionary model with gene features. *bioRxiv*. Published online May 21, 2023:2023.05.19.541520. doi:10.1101/2023.05.19.541520
17. Chen EY, Tan CM, Kou Y, et al. Enrichr: interactive and collaborative HTML5 gene list enrichment analysis tool. *BMC Bioinformatics*. 2013;14:128. doi:10.1186/1471-2105-14-128
18. Kuleshov MV, Jones MR, Rouillard AD, et al. Enrichr: a comprehensive gene set enrichment analysis web server 2016 update. *Nucleic Acids Res*. 2016;44(W1):W90-97. doi:10.1093/nar/gkw377
19. Xie Z, Bailey A, Kuleshov MV, et al. Gene Set Knowledge Discovery with Enrichr. *Curr Protoc*. 2021;1(3):e90. doi:10.1002/cpz1.90
